# Lipid tail chemistry regulates selective membrane interactions with model DNA nanoprobes and DNA-based coacervates

**DOI:** 10.1101/2025.10.11.681803

**Authors:** Yuhan Li, Krishnakavya Thaipurayil Madanan, Simran Dhanwantri, Akika Altman-Chandler, Nicholas G. Horton, Derek K. O’Flaherty, Roger Rubio-Sánchez, Claudia Bonfio

**Author notes:** These authors contributed equally (Yuhan Li, Krishnakavya Thaipurayil Madanan, Simran Dhanwantri).

## Abstract

Biological membranes actively regulate their composition to fine-tune their packing, fluidity, phase and surface charge, key properties that influence biomolecular interactions driving essential cellular pathways. While membrane surface charge is often attributed to specific lipid headgroups, the role of acyl-chain chemistry in modulating the interplay between these biophysical membrane properties remains unexplored. Here, we systematically investigate how acyl-chain length and saturation modulate lipid packing, fluidity, and membrane surface charge in zwitterionic lipid membranes. Using amphiphilic DNA nanoprobes as model charged biomolecules, we describe the interplay between packing, fluidity, phase and charge, identifying a packing-dependent guiding principle for membrane interactions that persists in the presence of anionic lipids. We also demonstrate that the identity and hydrophobicity of membrane anchors in nanoprobes significantly influence their binding to membranes. By integrating acyl-chain chemistry and membrane biophysical properties into design criteria for biomolecular attachment, our findings provide a mechanistic framework for engineering membrane interactions with both DNA nanoprobes and DNA-based coacervates. Beyond direct application to biomimetic platforms and synthetic cell engineering, these insights are relevant to lipid-based vaccine nanotechnologies and a fundamental understanding of membrane-biomolecule interactions in living cells.

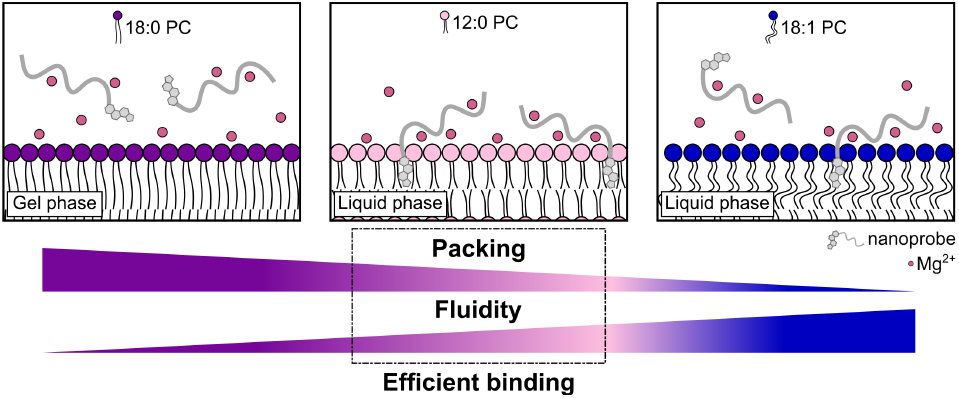

## Introduction

Biological membranes coordinate a vast array of essential cellular functions, from environmental sensing to molecular transport.^1^ Membrane-mediated functions rely on selective interactions between lipid bilayers and surrounding biomolecules. Such interactions, governed by electrostatic and hydrophobic forces, depend on the diverse charge profiles of biomolecules, their hydrophobic modifications, and the molecular heterogeneity of membranes, particularly the presence of charged lipids and bilayer-bound proteins and receptors.^1^

Cells actively regulate interactions between membranes and biomolecules, influencing diverse processes, such as protein insertion,^2,3^ transmembrane transport,^1^ and virus binding.^4^ The modulation of such interactions is traditionally ascribed to the diversity of lipid headgroups and their contribution to the overall surface charge of cell membranes.^5^ In contrast, lipid hydrophobic chains are often viewed mainly as structural regulators, affecting membrane packing, fluidity, and phase.^6^ However, variations in lipid tail structure have been shown to induce differences in lipid headgroup orientation,^7^ which may alter surface charge distribution and membrane propensity for interactions with biomolecules. Notably, the molecular structures of both lipid hydrophobic chains and lipophilic anchors of membrane proteins are highly diverse and often evolutionarily conserved,^8,9^ suggesting functional roles for these hydrophobic moieties beyond structural association. For instance, subtle adjustments to acyl chain length or saturation, which alter membrane packing and fluidity, could affect molecular recognition and binding.^10^ A refined understanding of how the molecular features of lipid hydrophobic chains influence biomolecular interactions with membranes would thus provide new insights into the principles of modern membrane biology and the potential to engineer membrane recognition and interactions with biomolecules for applications in synthetic biology, vaccine technologies, and gene delivery.

Because electrostatic forces strongly influence interactions between nucleic acids and lipid membranes,^7^ DNA-based nanoprobes, often covalently modified with lipophilic moieties like cholesterol, have been successfully used to elucidate phase- and cation-dependent binding of nucleic acids to membranes,^7,11^ and to monitor the presence of charged lipids in model membranes.^12^ These recent studies, however, have predominantly focused on tuning the structural properties of the DNA constructs, including their size, hydrophobicity, and flexibility. However, how the intrinsic physico-chemical properties of the target membrane govern these biomolecular interactions remains poorly understood. Amphiphilic DNA nanoprobes are therefore a powerful tool for unravelling how membrane packing, fluidity, phase, and surface charge collectively shape biomolecular interactions.

Here, we show that membrane recognition by amphiphilic nucleic acid probes is governed by a packing-dependent balance between insertion and retention, revealing a design rule that depends on both the membrane phase state and the hydrophobic character of the probe. Using amphiphilic nanoprobes comprising short single-strand DNA oligonucleotides conjugated with cholesterol or fatty acid motifs, we investigate how lipid chain length and unsaturation influence the packing, fluidity and surface charge of synthetic membranes. Our results suggest that nanoprobe-membrane interactions are influenced not only by the electrostatic profile of membranes but also strongly modulated by the hydrophobic chains of their constituent lipids. Generalised polarisation, fluorescence anisotropy, zeta potential, single-particle Raman spectroscopy, and confocal microscopy studies confirm that nanoprobe attachment to model membranes can be tuned by varying lipid packing and lateral diffusivity, even when the membrane surface charge remains constant. Importantly, we show that this packing-dependent rule underlies selective probe recruitment across mixed membrane populations and extends to higher-order biomolecular assemblies, including coacervates. Our findings demonstrate that the hydrophobic core of lipid bilayers is not merely a passive component of cell membranes, but actively influences their biomolecular interactome, providing an indirect handle for regulating the recognition and binding of (charged) biomolecules. Our work offers mechanistic insights into membrane-biomolecule interactions with broad implications in membrane biophysics, programmable biomolecular recognition, and lipid-based nanotechnology.

## Results and Discussion

### Lipid packing influences membrane surface charge and DNA nanoprobes binding

We initially focused on single-component membranes comprising phosphatidylcholine (PC) lipids, which, despite possessing zwitterionic headgroups, yield membranes that are slightly negatively charged.^7,11,13^ PC membranes thus provide an optimal model to disentangle how lipid packing, membrane fluidity, and surface charge influence cation-mediated DNA nanoprobe binding (Fig. 1a). We first produced Large Unilamellar Vesicles (LUVs, d_H_ ~ 200 nm) employing either dilauroyl phosphatidylcholine (12:0 PC, T_m_= −2°C), distearoyl phosphatidylcholine (18:0 PC, T_m_= 55°C), or dioleoyl phosphatidylcholine (18:1 PC, T_m_= −17°C), and measured their zeta ( ζ) potential *via* electrophoretic light scattering (Fig. 1b). Despite all phospholipids having the same headgroup, differences in surface charge across membranes readily emerged, with ζ-potential values following the trend 18:0 PC > 12:0 PC > 18:1 PC, suggesting a modulating role of lipid chain length and unsaturation on membrane surface charge.

**Figure 1.**
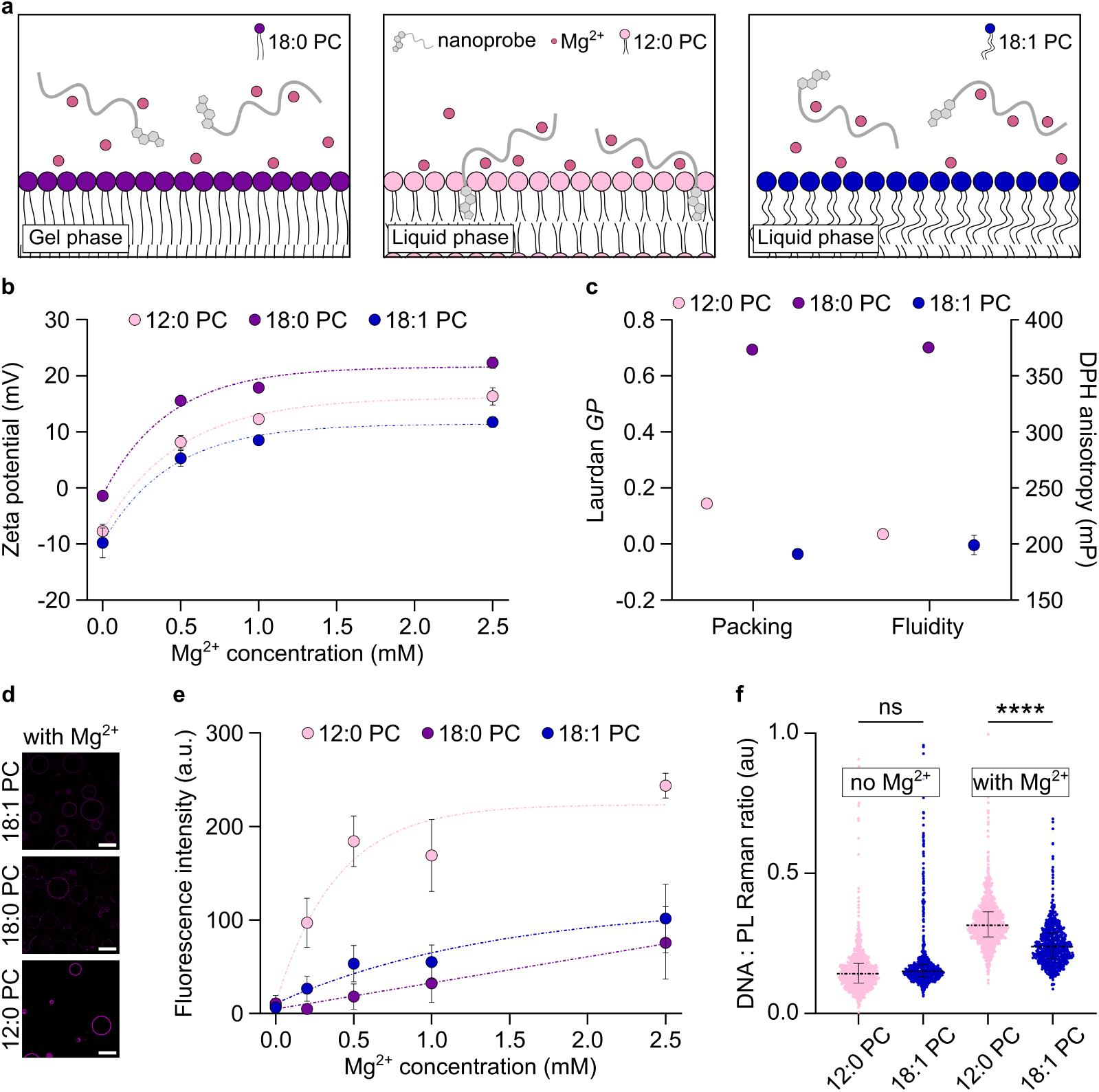
Influence of membrane biophysical properties on DNA nanoprobe binding. **a.** Schematic illustrating the differential interactions observed between amphiphilic DNA nanoprobes with membranes comprising 18:0 (left), 12:0 (middle), and 18:1 (right) PC lipids, as enabled by charge screening with cations, lipid packing, and membrane fluidity. **b**. ζ-potential of Large Unilamellar Vesicles (LUVs) composed of 12:0, 18:0, or 18:1 PC lipids and dispersed in buffering solutions supplemented with increasing concentrations of MgCl_2_. **c**. Membrane packing (left) and fluidity (right) probed *via* Laurdan Generalised Polarisation (*GP*) and anisotropy of 1,6-Diphenyl-1,3,5-hexatriene (DPH), respectively. **d**. Representative confocal micrographs of 12:0, 18:0, or 18:1 PC Giant Unilamellar Vesicles (GUVs) showing the differential attachment of fluorescent DNA nanoprobes (signal in magenta) at 0.5 mM MgCl_2_. Scale bars = 20 µm. **e**. Membrane attachment of fluorescent DNA nanoprobes at increasing concentrations of MgCl_2_determined by sampling membrane fluorescence intensity from confocal micrographs. **f**. Quantification of the Cy3-DNA Raman signal of individual particles for distinct membranes in the absence or presence of Mg^2+^ (0.5 mM), normalised to the choline signal of PC lipids (1385-1405/700-730 cm^−1^) (median ± IQR, n ≥ 893). The Cy3 Raman peak was chosen as the binding reporter because its signal does not overlap with phospholipid signals.

At room temperature, 18:0 PC forms gel-phase membranes,^14^ while shorter acyl-chain lipids, e.g., 12:0 PC, and unsaturated acyl-chain lipids, e.g., 18:1 PC, generate liquid-phase membranes.^15,16^ Through fluorescence anisotropy measurements with 1,6-Diphenyl-1,3,5-hexatriene (DPH) (Fig. 1c), we confirmed that 18:0 PC membranes exhibit anisotropy values consistent with gel phases (375.29 mP),^17^ while 12:0 PC and 18:1 PC membranes (208.78 mP and 202.23 mP, respectively) exhibit anisotropy values consistent with liquid-phase membranes. Generalised Polarisation (*GP*) measurements using the solvatochromic Laurdan dye^18^ revealed subtle differences in lipid packing between the two liquid-phase membranes (*GP*_12:0 PC_= 0.14 *vs GP*_18:1 PC_= −0.03). Specifically, 12:0 PC membranes, despite the shorter acyl chains of their constituent lipids, exhibited tighter packing than 18:1 PC membranes, yet lower than 18:0 PC membranes (*GP*_18:0 PC_= 0.69) (Fig. 1c). Note that spectroscopy measurements on LUVs indicate that Laurdan *GP* is agnostic to overall lipid concentration (Fig. S1). The variations in lipid packing observed across the three model membranes are ascribed to differences in acyl-chain unsaturation and length, parameters that could collectively influence the overall membrane surface charge.

We next supplemented our buffered solution (300 mM sucrose + 5 mM HEPES, pH 7.5) with increasing concentrations of magnesium chloride (MgCl_2_). Upon Mg^2+^ titration, ζ-potential measurements revealed similar bilayer-cation interactions among all three types of membranes (Fig. 1b). Specifically, membrane surface charge monotonically increased upon Mg^2+^ titration, eventually becoming positive, while preserving the relative order imposed by acyl-chain length and unsaturation ( ζ_18:0 PC_> ζ_12:0 PC_> ζ_18:1 PC_). Notably, the addition of as little as 0.5 mM Mg^2+^ (equimolar to the lipid concentration) is sufficient to achieve a positive surface charge for all three types of LUVs studied herein.

To assess how the identified differences in lipid packing and surface charge influence membrane interactions with charged biomolecules, we employed a fluorescently labelled (Cy3) amphiphilic DNA nanoprobe, comprising a 12-nucleotide single-strand oligonucleotide covalently 3′-linked to a hydrophobic cholesterol moiety *via* a tetraethylene glycol spacer. This simple, unstructured amphiphilic ssDNA nanoprobe, whose binding is highly sensitive to variations in ionic strength,^7^ allowed us to investigate how bilayer biophysical properties govern membrane attachment. We produced non-fluorescently labelled Giant Unilamellar Vesicles (GUVs) and incubated them with DNA nanoprobes (upon conditions optimisation, Figs. S2-S5) for one hour at room temperature to allow for membrane-nanoprobe interaction in the presence of Mg^2+^. We observed clear differences in membrane fluorescence intensity, reflecting variations in nanoprobe binding. Control experiments with a Cy3-labelled DNA nanoprobe lacking the hydrophobic modification showed no detectable membrane accumulation (Fig. S4), ruling out non-specific electrostatic interactions between the DNA backbone and the lipid bilayer and confirming that cholesterol insertion into the membrane enables binding. Moreover, dynamic light scattering of the amphiphilic DNA nanoprobe in solutions with or without Mg^2+^ (Fig. S6) showed no evidence of probe aggregation at the experimental concentration (100 nM), confirming that membrane fluorescence intensity reflects probe insertion into bilayers rather than surface accumulation of aggregates. Importantly, zeta-potential measurements indirectly validate membrane association of nanoprobes, as shown by changes in LUV surface charge before and after adding Mg^2+^ and DNA (Fig. S7).

Quantitative image analysis confirms that, consistent with differences in surface charge, 12:0 PC membranes interact more efficiently with DNA nanoprobes than 18:1 PC membranes (Figs. 1d-e). Fluorescence recovery after photobleaching (FRAP) of membrane-bound nanoprobes demonstrates that, upon association, DNA nanoprobes remain laterally mobile and diffusive within fluid bilayers (Fig. S8). Moreover, higher Mg^2+^ concentrations lead to stronger nanoprobe binding, likely because divalent cations screen negative charges and enable electrostatic interactions between phosphate groups on both oligonucleotides and lipids.^7^ Surprisingly, DNA nanoprobes exhibited a lower affinity for 18:0 PC membranes (Fig. 1d-e), despite their less negative surface charge relative to their liquid counterparts. We attribute this result to the low lateral diffusivity of the 18:0 PC gel-phase bilayers. We argue that, by limiting nanoprobe translational motion when membrane-anchored compared to its unrestricted diffusion in solution, membrane-DNA interactions entail a significant entropic penalty.

We hypothesise that membrane attachment entails a significant entropic penalty by restricting nanoprobe translational and conformational degrees of freedom relative to its unrestricted diffusion in solution.^19–21^ Anchoring a probe to a fluid membrane preserves 2D translational entropy *via* lateral diffusion,^19^ whereas binding to a gel-phase membrane with restricted lateral mobility would result in a loss of both 3D and 2D translational freedom. Additionally, the tight packing of gel-phase membranes likely limits access to the bilayer hydrophobic core for amphiphilic DNA, thereby hindering insertion of cholesterol anchors into the membrane.

To independently validate the membrane-binding trends observed by confocal microscopy, we quantified nanoprobe association using single-particle Raman spectroscopy by monitoring DNA-associated Raman signatures of liposomes incubated with the cholesterol-DNA probe in the presence of 0.5 mM Mg^2+^. We observed a distinctive band at approximately 1394 cm^-1^,^22^ consistent with the symmetric deformation of methyl groups in the Cy3-labelled DNA construct (Figs. S9-S10). Importantly, the intensity of this band was substantially higher for 12:0 PC membranes than for 18:1 PC membranes, indicating approximately 1.5-fold greater probe association with the more tightly packed liquid-phase bilayer (Figs. 1f and S11). These measurements establish single-particle Raman spectroscopy as an orthogonal readout for membrane recruitment of amphiphilic DNA constructs and independently confirm that DNA nanoprobe binding is enhanced on tightly packed liquid-phase membranes.

### Acyl-chain length modulates lipid packing and surface charge in zwitterionic membranes

After identifying lipid chain-dependent membrane-DNA nanoprobe interactions, we systematically explored how chain length affects lipid packing, surface charge, and, ultimately, binding. We first compared the biophysical properties of 12:0 PC and 18:0 PC membranes with those of bilayers comprising dimyristoyl phosphatidylcholine (14:0 PC), dipalmitoyl phosphatidylcholine (16:0 PC), and diarachidoyl phosphatidylcholine (20:0 PC). To minimise lipid degradation (e.g., hydrolysis, oxidation, and polymerisation) catalysed by heat exposure, we prepared membranes from high-melting-temperature lipids using a non-extrusion protocol (see Methods). We thus confirmed that differences in sample polydispersity do not bias our membrane packing measurements with Laurdan *GP* (Fig. S12). While 12:0 PC forms liquid-phase membranes at room temperature, the remaining lipids form gel-phase membranes,^6^ as confirmed by DPH anisotropy (Fig. S13). A monotonic increase in Laurdan *GP* values was observed as a function of acyl-chain length, consistent with the different phases of 12:0 PC and 14:0 PC membranes and the progressively tighter lipid packing regime within the gel phase (Fig. S14). Across these gel-phase membranes, a trend also emerges in zeta potential values ( ζ_18:0 PC_> ζ_16:0 PC_> ζ_14:0 PC_), with shorter chain lipids yielding more negatively charged membranes (Fig. S15). Upon addition of Mg^2+^ ions, all membranes exhibited a shift towards positive ζ-potential values, consistent with divalent cation binding and charge screening (Fig. S15). Differences in membrane surface charge across the tested zwitterionic PCs likely reflect chain-length-dependent variations in lipid packing and fluidity.

To understand how chain-length-induced changes in membrane properties influence membrane interactions with DNA nanoprobes, we explored relationships among nanoprobe binding, lipid packing, membrane fluidity, and surface charge. We incubated GUVs composed of the saturated PCs mentioned above with DNA nanoprobes and quantified their interactions. We observed differences in membrane fluorescence intensity and, thus, nanoprobe binding, with higher binding efficiency for membranes composed of shorter-chain lipids (Figs. 2a and S16). Increasing Laurdan *GP* values, indicative of tighter lipid packing, are associated with decreasing nanoprobe attachment (coefficient of determination, R^2^ = 0.99) (Fig. 2b). Although membrane fluidity showed a non-monotonic, convex dependence on chain length, decreasing from 12:0 PC to 16:0 PC and then increasing again for 18:0 PC and 20:0 PC, nanoprobe binding did not follow this trend (Fig. S17). Instead, nanoprobe attachment decreased monotonically with increasing chain length, with 18:0 PC and 20:0 PC both showing very low binding, consistent with their tighter packing relative to shorter-chain PCs. Interestingly, 12:0 PC membranes showed stronger binding, with lower lipid packing and a less positive surface charge in the presence of Mg^2+^ ions (Fig. S18). However, no correlation was observed for gel-phase membranes between packing and charge (R^2^ = 0.28) (Fig. S19). We thus conclude that tight lipid packing in gel-phase membranes, even with membrane fluidity and surface charges favourable for binding, hinders nanoprobe attachment, likely by restricting access to the hydrophobic core for the cholesterol-based anchor in DNA nanoprobes, in a manner that scales with lipid acyl-chain length.

**Figure 2.**
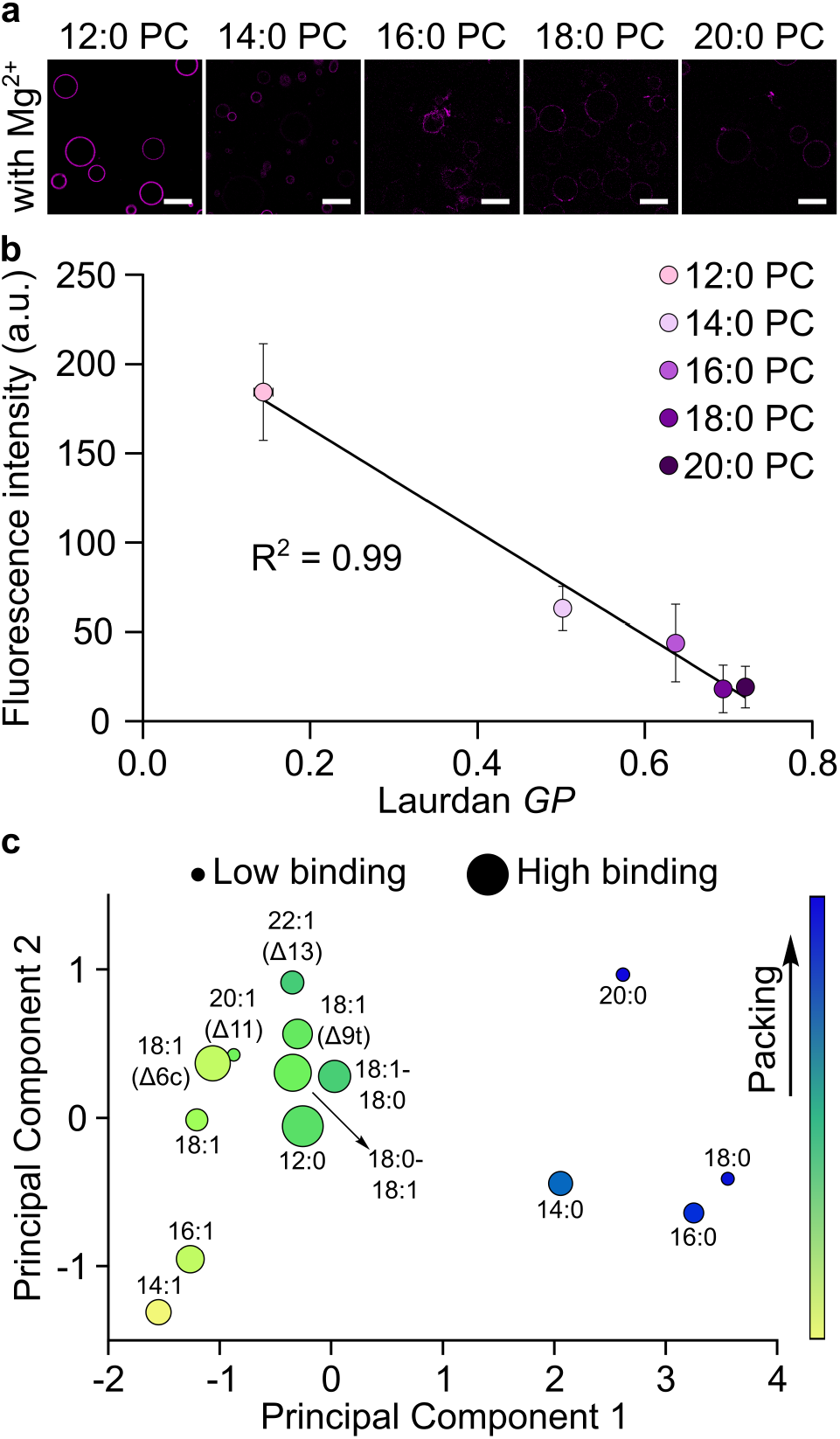
Lipid packing influences DNA nanoprobes binding to zwitterionic membranes. **a.** Representative confocal micrographs of Giant Unilamellar Vesicles (GUVs) comprising phospholipids of increasing acyl-chain length showing the differential attachment of fluorescent DNA nanoprobes in the presence of 0.5 mM MgCl_2_(in magenta). Scale bars = 20 µm. **b**. Membrane attachment of nanoprobes in the presence of 0.5 mM MgCl_2_, as observed in confocal micrographs, is negatively correlated with lipid packing, as measured by Laurdan Generalised Polarisation (*GP*) (R^2^ = 0.99), with tighter packing corresponding to lower fluorescence intensities. **c**. Principal Component Analysis (PCA) of membrane biophysical descriptors across the tested lipid library (Table S1). PCA was performed using lipid packing, membrane fluidity, surface charge and melting temperature as input variables. Points are coloured from low to high membrane packing (inferred from Laurdan *GP*), and symbol size indicates nanoprobe binding (inferred from fluorescence intensity), with smaller dots corresponding to less efficient binding and larger dots to more efficient binding.

We next examined how membrane-DNA nanoprobe interactions vary with increasing acyl chain length in unsaturated phospholipids, carrying a single unsaturation per acyl chain, within two selected lipid libraries. Liposomes were prepared using either Δ9 PCs, including dimyristoleoyl phosphatidylcholine (14:1 D9 PC), dipalmitoleoyl phosphatidylcholine (16:1 D9 PC), or Δn-9 PCs (where n corresponds to the number of carbons in the acyl chain), including dieicosenoyl phosphatidylcholine (20:1 D11 PC) and dierucoyl phosphatidylcholine (22:1 D13 PC). We used 18:1 Δ9 PC in both screenings for comparison. Except for 22:1 PC, all lipid compositions yield liquid-phase membranes with comparable fluidities (~200 mP) (Fig. S20); 22:1 PC membranes exhibit fluidity comparable to that of 12:0 PC membranes. Laurdan *GP* measurements reveal a monotonic increase in lipid packing with lipid chain elongation (Fig. S21), with 22:1 PC membranes exhibiting tighter packing than liquid-phase 12:0 PC membranes. ζ-potential studies showed differences in membrane surface charge across the tested lipid library (Fig. S22). These differences were reduced upon addition of Mg^2+^ ions (0.5 mM, equimolar to the lipid concentration), which resulted in a decrease in surface charge with increasing chain length. Still, all membranes exhibited a positive ζ-potential value in the presence of divalent cations, reflecting efficient charge screening. Interestingly, in contrast to the results for gel-phase membranes, we observed a mild negative correlation between packing and surface charge for liquid-phase membranes composed of unsaturated PCs (R^2^ = 0.77) (Fig. S23). However, nanoprobe attachment varied non-monotonically with acyl chain length, with 16:1 PC showing the strongest membrane-DNA interactions, 20:1 PC the weakest among unsaturated PC membranes, and 18:1 PC and 22:1 PC exhibiting intermediate behaviour (Fig. S24).

We ascribe the differences we observed in DNA nanoprobe binding to variations in double-bond position. It has recently been demonstrated that cholesterol interacts with and condenses acyl chains to a differential extent depending on the exact position of unsaturation along the acyl chain.^23^ The differential binding observed within the two unsaturated-lipid series suggests that nanoprobe-membrane interactions are not solely dictated by acyl chain length but are also influenced by the position of unsaturation, which would likely affect the insertion and positioning of the cholesterol anchor.

Consistent with their higher fluidity compared to the fully saturated PC library (Figs. S13 and S17), all liquid-phase membranes composed of unsaturated PCs exhibited more efficient nanoprobe binding than their gel-phase counterparts, which comprised lipids with the same acyl-chain length, except for both 20:0 PC and 20:1 PC, which exhibited near-zero binding. However, we observed no correlation between nanoprobe binding and lipid packing, membrane fluidity, or surface charge in unsaturated PC membranes (Figs. S25-S27). Notably, nanoprobe binding to liquid-phase membranes comprising 12:0 PC was substantially higher than that to membranes composed of unsaturated PCs. Despite having a comparable surface charge to membranes composed of 14:1, 16:1, 18:1, and 20:1 PCs, liquid-phase 12:0 PC-based membranes exhibit tighter packing and lower fluidity, seemingly favouring the incorporation of the cholesterol-functionalised DNA nanoprobe.

Altogether, our results demonstrate that membrane surface charge is not the sole parameter regulating membrane-DNA nanoprobe interactions. In gel-phase membranes, looser packing observed with shorter-chain phospholipids improves nanoprobe insertion. In contrast, in liquid-phase membranes, tighter packing affords more efficient nanoprobe binding, with 20:1 and 22:1 PCs notably deviating from the trend. Our findings reflect a highly intricate interplay between lipid acyl-chain length and membrane packing, and their influence on membrane surface charge. By modulating the hydrophobic core of model phospholipid membranes and thereby altering its packing within the bilayer, we show that cation-mediated membrane-nanoprobe binding can be finely tuned, even without changes to the headgroups of the constituent lipids.

To integrate the multiple membrane descriptors measured across our full lipid library rather than comparing only subsets of said library based on individual parameters (e.g., presence of double bonds, position or conformation of double bonds, and length of the lipid acyl chains), we next performed principal component analysis (PCA)^24^ using lipid packing, membrane fluidity, surface charge and melting temperature as input variables (Figs. 2c and S28). The first two principal components captured the majority of the variance in the dataset (PC1 = 85.3% variance, PC2 = 10.7% variance), indicating that these biophysical descriptors are not independent but instead define a limited number of coupled membrane-state dimensions. Along the two principal components, the data cluster by membrane properties representing lipid packing and membrane phase, which appear to be the major determinants of DNA-binding trends. Specifically, the more tightly packed, gel-phase membranes cluster separately from the less packed membranes. However, we observed no sub-clustering within the liquid-phase space, despite higher DNA-binding efficiencies primarily observed in well-packed liquid-phase membranes by single-particle Raman spectroscopy and fluorescence microscopy (Fig. S28).

Overall, these results collectively indicate a design rule in which efficient membrane association requires an intermediate lipid packing state: packing must be sufficient to allow efficient insertion of the cholesterol anchor, but not so extreme as to suppress bilayer accessibility and lateral mobility.

### Membrane packing governs nanoprobe binding, even when anionic lipids modulate the membrane surface potential

To test whether the packing-dependent membrane-binding trends observed for zwitterionic PC bilayers also persist when membrane charge is intentionally altered, we prepared mixed membranes containing 12:0 PC or 18:1 PC doped with 10% or 20% anionic lipids of matching chain length. We considered both fatty acids and phosphatidic acids, which differ in charge density and membrane-incorporation behaviour, to probe how electrostatic and hydrophobic contributions combine to determine nanoprobe recruitment.

We expected anionic lipid incorporation to alter membrane surface charge by introducing negatively charged groups in the lipid headgroups. Phosphatidic acids, bearing one to two negative charges at neutral pH (pK_a_ 7.2), were expected to render the membrane surface more negative than fatty acids, bearing zero to one negative charge (apparent pK_a_= 7.5-8.5, depending on acyl chain length) (Figs. S29-30). As expected, phosphatidic-acid-containing membranes exhibit more negative z-potentials (between −50 and −70 mV) than the corresponding fatty acid-containing systems (between −25 and −45 mV). In parallel, spectroscopic measurements showed that phosphatidic acids perturb the bilayer hydrophobic core more strongly than fatty acids (Figs. S31-34), consistent with fatty acids being present in non-negligible amounts as free in solution rather than membrane-bound.^25–27^ These observations indicate that adding anionic lipids modulates both the electrostatic and structural properties of the membrane in a composition-dependent manner.

Upon incubation with the DNA nanoprobes, membrane binding remained sensitive to both charge and packing: 12:0 PC-based membranes generally enabled stronger recruitment than 18:1 PC-based membranes, and fatty acid-containing membranes supported more efficient nanoprobe binding than phosphatidic acid-containing membranes (Figs. S34-S40). Consistently, nanoprobe attachment positively correlated with surface charge (Figs. 3a and S41) and negatively correlated with lipid packing (Figs. 3b and S42), in line with the observed inverse correlation between packing and charge (R^2^ = 0.83 for 12:0 PC-based membranes, Figs. S43-44), thereby providing a useful framework to tune membrane-nanoprobe interactions.

**Figure 3.**
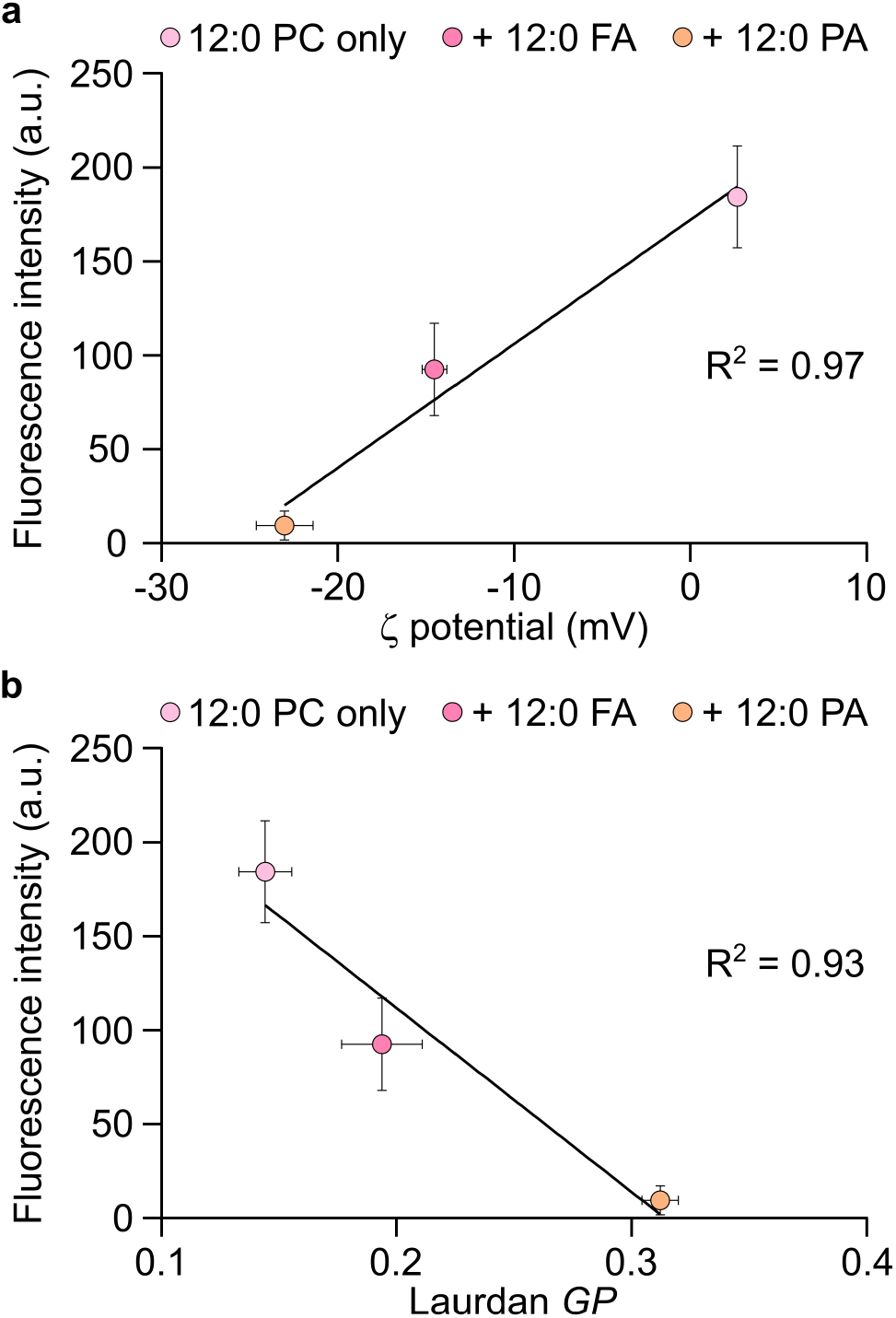
Anionic inclusions influence the membrane attachment of DNA nanoprobes by altering lipid packing and surface charge. **a.** Nanoprobe binding to membranes doped with 20% anionic lipids in the presence of 0.5 mM MgCl_2_is governed by surface charge, with a positive correlation between membrane fluorescence intensity and zeta potential (R^2^ = 0.97). **b**. Nanoprobe binding to membranes doped with anionic lipids in the presence of 0.5 mM MgCl_2_is modulated by lipid packing, showing negatively correlated fluorescence intensities (sampled from confocal micrographs) with their corresponding Laurdan Generalised Polarisation (*GP*) values (R^2^ = 0.93).

Overall, these data show that the packing-dependent binding rule established for zwitterionic membranes also applies to negatively charged bilayers. Even in the presence of anionic lipids, efficient recruitment appears to require a membrane that is sufficiently packed to support efficient insertion, but not so tightly packed as to compromise accessibility or lateral diffusion of the amphiphilic DNA probe.

### Nanoprobe attachment is governed by phase-dependent lipid packing and the identity of the lipophilic anchor

Having established that lipid packing and membrane surface charge vary with acyl chain length in both liquid- and gel-phase membranes, we next sought to disentangle the specific contribution of membrane phase state. To do so, we used lipids of identical chain length and varying cholesterol levels to generate membranes in distinct physical states, enabling us to evaluate the role of membrane phases in mediating interactions with DNA nanoprobes.

Specifically, 18-carbon phospholipids were employed to produce membranes in the gel phase (18:0 PC), liquid-ordered phase (*L*_o_; 60% 18:0 PC + 40% cholesterol), or liquid-disordered phase (*L*_d_; 60% 18:1 PC + 40% cholesterol or 18:1 PC).^14^ Membrane phase was evaluated by DPH anisotropy and Laurdan *GP* measurements, reflecting differences in membrane fluidity (Fig. 4a) and lipid packing (Fig. S45). ζ-potential measurements revealed that all liquid-phase membranes exhibited a more negative surface potential than gel-phase membranes (Fig. S46). Upon Mg^2+^ addition, all systems shifted towards positive ζ-potentials, but the 18:0 PC membranes remained the most positively charged, confirming that Mg^2+^ interacts more readily with more tightly packed bilayers.^7^

**Figure 4.**
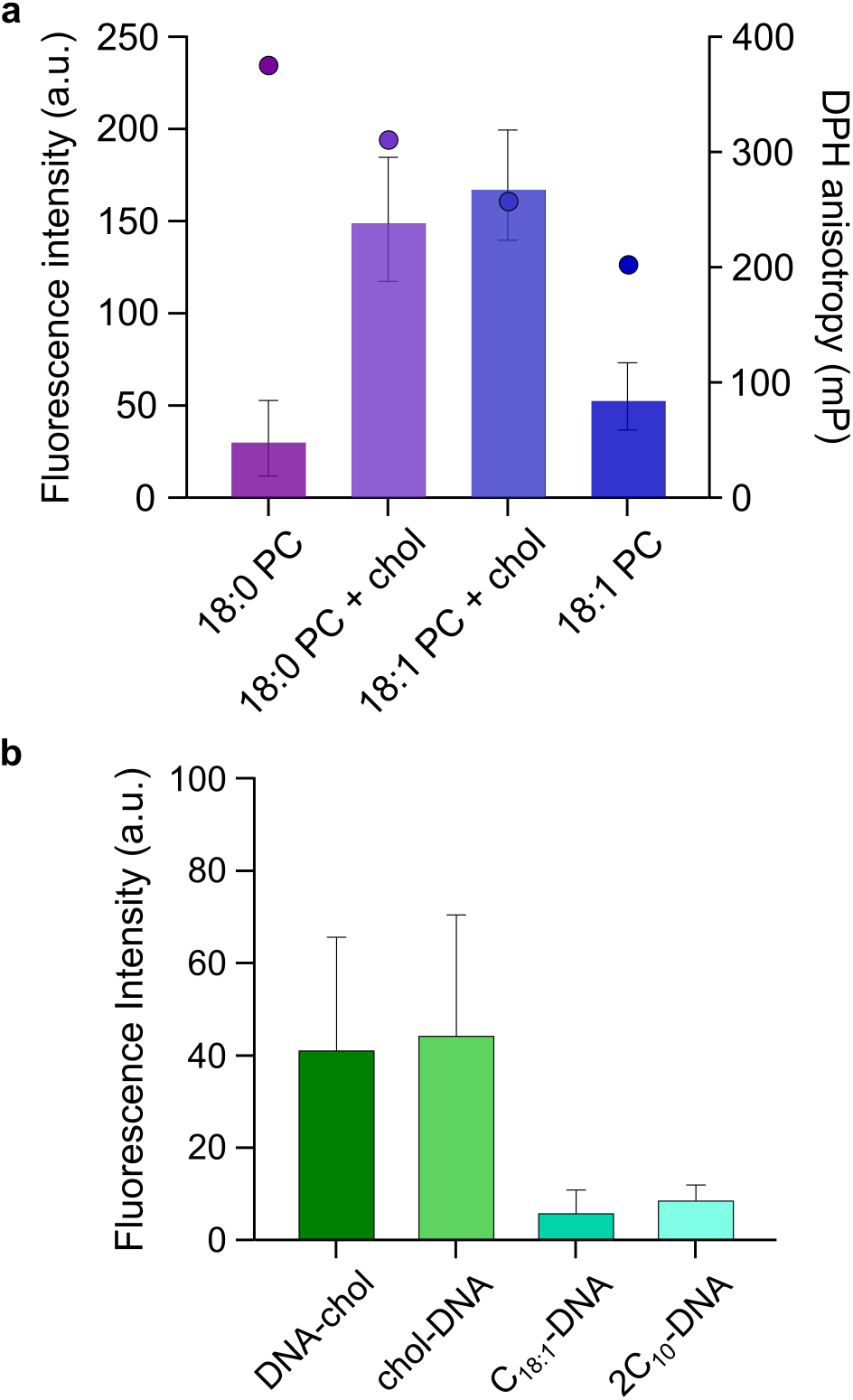
Phase-dependent membrane packing and anchor identity regulate DNA nanoprobe binding. **a.** Membrane fluorescence intensities of DNA nanoprobes in the presence of 0.5 mM MgCl_2_(bar plot, sampled from confocal micrographs) overlaid with anisotropy of 1,6-Diphenyl-1,3,5-hexatriene (DPH) measured in Large Unilamellar Vesicles (LUVs) (circles), showing the differences in membrane fluidity as a function of phase: gel (18:0 PC), liquid-ordered (18:0 PC + cholesterol), liquid-disordered (18:1 PC), and liquid-disordered supplemented with cholesterol (18:1 PC + cholesterol). **b**. Membrane intensity of nanoprobes featuring different anchor groups in the presence of 0.5 mM MgCl_2_: cholesterol on the 3′ as well as 5′ end, C_18:1_or double-C_10_(no spacer).

The strongest nanoprobe attachment occurred in the cholesterol-enriched membranes, with the two cholesterol-containing systems, 18:0 PC + cholesterol and 18:1 PC + cholesterol (Figs. 4a and S47), showing similarly high binding despite their different packing and phase properties. In contrast, the gel-phase 18:0 PC membranes bound substantially less nanoprobe (Figs. 4a and S47), consistent with the previously identified requirement for an intermediate packing regime. Thus, while gel-phase membranes are too rigid to support efficient recruitment, the comparable binding of the two cholesterol-enriched membranes suggests that membrane association is not determined by packing alone. Instead, cholesterol-rich bilayers appear to provide a membrane environment that remains compatible with the cholesterol-based anchor, as previously observed for longer single-strand DNA sequences,^28^ even when lipid packing differs. We hypothesise that cholesterol-rich membranes enhance incorporation of cholesterol-tagged DNA probes because pre-existing cholesterol promotes favourable packing in liquid-phase membranes, thereby increasing the effective affinity of additional cholesterol-bearing species.^29–31^

Importantly, for *L*_o_, *L*_d_, and cholesterol-enriched *L*_d_membranes, nanoprobe attachment showed a weak inverse correlation with membrane surface charge (R^2^ = 0.60) (Fig. S48). Analogously, mildly stronger positive correlations were observed with lipid packing (R^2^ = 0.68) and membrane fluidity (R^2^ = 0.64) (Figs. S49-50). Collectively, these data indicate that membrane packing and phase still modulate nanoprobe recruitment in cholesterol-enriched membranes. Yet, membrane-embedded cholesterol can introduce an additional layer of cholesterol-tagged DNA-binding specificity beyond packing or fluidity alone.

Having explored how the membrane hydrophobic core affects membrane-nanoprobe interactions, we next investigated how the identity of the lipophilic anchor influences binding to membranes with different packing and surface charge. We synthesised a small library of FAM-labelled DNA nanoprobes featuring distinct 5′-hydrophobic motifs. After confirming that substitution of the DNA fluorophore does not affect membrane-nanoprobe interactions (Figs. S51-52), we first asked whether the position of the cholesterol anchor, at either the 3′ or 5′ end of the oligonucleotide, alters attachment to 12:0 PC membranes; no statistically significant difference was observed (Fig. 4b).

We then compared nanoprobes bearing different hydrophobic modifications: a single 10-carbon (C10) tail, with or without a hexaethylene glycol spacer, two C10 tails, and a single unsaturated 18-carbon (C18:1) tail (Figs. S53-56). Quantification of membrane attachment revealed that only the double-C10- and C18:1-acylated nanoprobes moderately interacted with 12:0 PC membranes, whereas single C10-anchored nanoprobes showed no detectable membrane association, irrespective of the spacer (Figs. 4b and S57). These differences are consistent with the distinct hydrophobic character of the anchors, with cholesterol and long acyl chains being more hydrophobic than short acyl chains. Indeed, C10 anchors are less likely to partition efficiently into lipid bilayers, since short-chain fatty acids can exchange between membrane-bound and soluble states more readily than cholesterol.^26,27,32^ As a result, short acyl-chain anchors, particularly C10, may reduce DNA recruitment by leaving a larger fraction of nanoprobes in solution at equilibrium.

Overall, these findings show that cholesterol-enriched membranes are unexpectedly permissive to cholesterol-based probes, suggesting that nanoprobe recruitment depends not only on lipid packing but also on compatibility between the membrane environment and the probe anchor. Membrane interactions thus appear to be tuneable independently through both lipid composition (*via* packing, surface charge, and phase) and the chemical identity of the hydrophobic anchor. Moreover, both membrane and probe design provide tuneable, independent handles to regulate interactions. The same membrane can favour different probes depending on the hydrophobic handle, and conversely, the same probe can behave differently across lipid states.

### Lipid packing enables selective recruitment of nanoprobes and coacervates to synthetic membranes

Having established that membrane packing and phase state govern nanoprobe recruitment across zwitterionic, anionic, and cholesterol-containing bilayers, we next asked whether these biophysical differences could drive selective binding within coexisting membrane populations and with higher-order biomolecular assemblies. We first compared mixed membrane systems differing in packing and phase and observed preferential association of the amphiphilic DNA probes with the composition that displayed the most favourable balance of packing and fluidity. Specifically, cholesterol-based nanoprobes preferentially localised to liquid-phase membranes, consistent with their higher affinity for fluid membranes. Importantly, when incubated with 18:0 PC or 18:1 PC membranes, C18:1-based nanoprobes showed only weak and mostly unselective binding (Fig. S58). In contrast, cholesterol-DNA conjugates bound most efficiently to cholesterol-enriched phases and, secondarily, to *L*_d_ membranes (Fig. 5a-b). To directly probe selectivity, we co-incubated cholesterol- and C18:1-functionalised DNA nanoprobes with gel-phase membranes (18:0 PC) or cholesterol-enriched *L*_d_ membranes (60% 18:1 PC + 40% cholesterol). As expected, cholesterol-based nanoprobes preferentially localised to liquid-phase membranes, consistent with their differences in packing and fluidity (Figs. 5a-b and S59). Conversely, C18:1-based nanoprobes showed a moderate preference for gel-phase membranes, indicating that the hydrophobic anchor identity can drive membrane selectivity.

**Figure 5.**
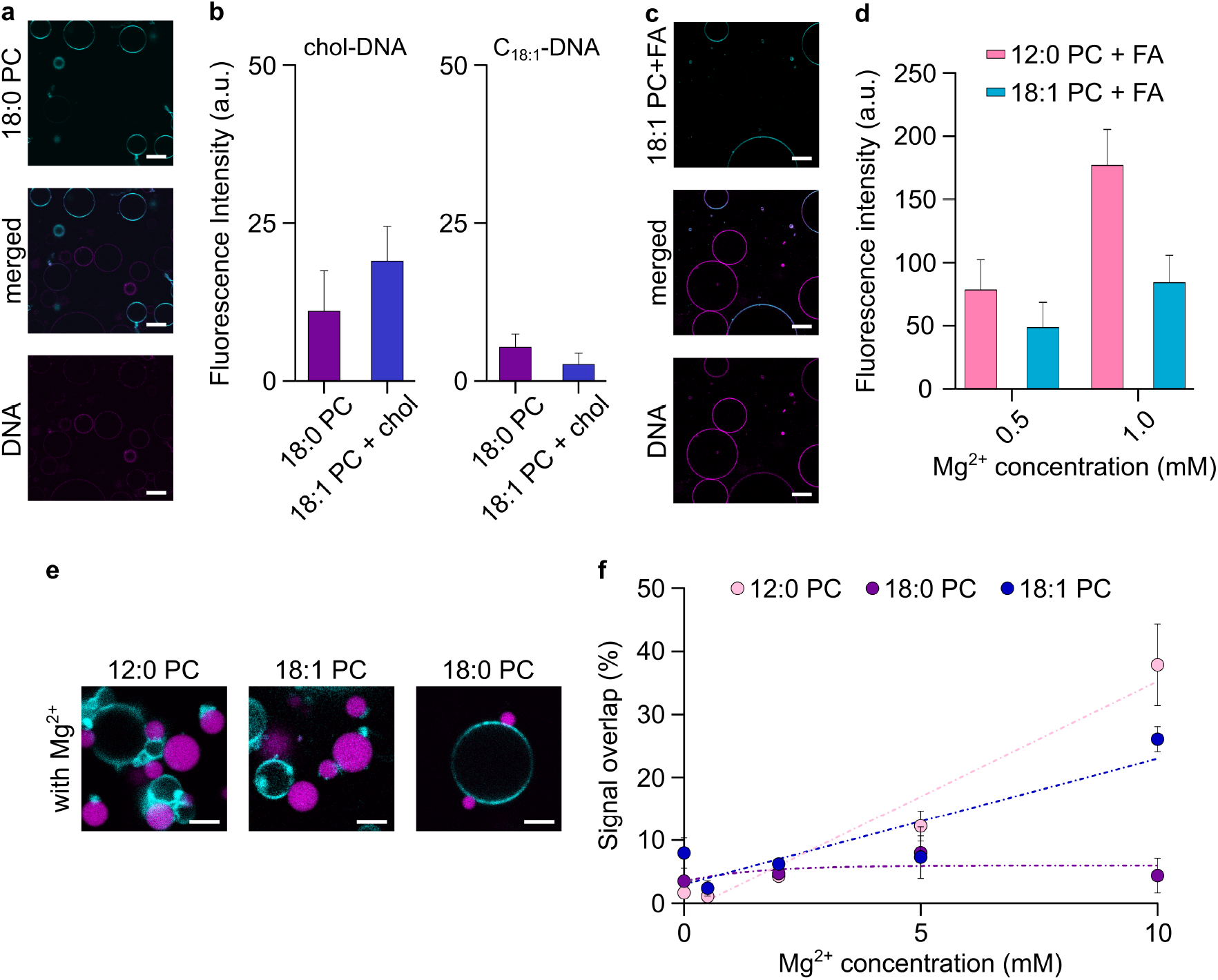
Lipid acyl chain composition regulates selective membrane interactions with DNA nanoprobes and DNA-based coacervates. **a.** Representative confocal micrographs of nanoprobe binding selectivity assay between 12:0 PC + FA and 18:1 PC + FA Giant Unilamellar Vesicles (GUVs) in the presence of 0.5 mM MgCl_2_, showing higher FAM-DNA nanoprobe attachment (signal in magenta) to 12:0 PC + FA membranes (non-fluorescent) relative to 18:1 PC + FA GUVs (stained with TexasRed-DHPE, signal shown in cyan). Scale bars = 20 µm. **b**. Nanoprobe fluorescence intensity in membranes sampled from confocal micrographs, showing the preferential attachment to 12:0 PC + FA GUVs in the presence of 0.5 mM MgCl_2_. **c**. Representative confocal micrographs of the nanoprobe selectivity binding assay, showing preferential attachment of cholesterol-conjugated nanoprobes to liquid-disordered membranes (60% 18:1 PC + 40% chol) enriched with cholesterol relative to gel-phase GUVs (18:0 PC) in the presence of 0.5 mM MgCl_2_. Scale bars = 20 µm. **d**. Membrane-associated fluorescence intensities of nanoprobes, featuring either cholesterol (left) or C18:1 (right) anchors, in a selectivity assay in the presence of 0.5 mM MgCl_2_. Preferential binding of the chol-DNA nanoprobes is observed for liquid-phase membranes, while C18:1 shows weak preferential attachment to gel-phase membranes. **e**. Representative confocal micrographs of GUV-coacervate interactions, showing preferential binding of DNA-based coacervates (FAM-labelled DNA signal in magenta) to 12:0 PC membranes relative to 18:1 PC or 18:0 PC GUVs (stained with TexasRed-DHPE, signal shown in cyan) in the presence of 5 mM MgCl_2_. Scale bars = 5 µm. **f**. Coacervate-membrane interactions at increasing concentrations of MgCl_2_determined by quantification of the fluorescence signal overlap between coacervates and GUVs, extracted from confocal micrographs.

We then extended our selectivity studies to anionic membranes exhibiting different packing. When nanoprobes were incubated with mixed populations of 12:0 PC + fatty acid and 18:1 PC + fatty acid membranes, we observed preferential attachment of the DNA nanoprobe to the more tightly packed 12:0 PC-containing membranes, in agreement with the stronger binding observed for this composition in the single-population experiments (Fig. 5c-d). These observations show that membrane selectivity can be driven by the hydrophobic core rather than by headgroup charge alone, and that the same packing-dependent rule persists even in the presence of anionic lipids.

Finally, we moved from composition-dependent selection in mixed-membrane populations to preferential membrane recruitment of coacervates. To test whether the membrane-recognition principles identified for simple DNA nanoprobes extend to DNA oligonucleotides that are initially pre-organised as core constituents of biomolecular assemblies, we evaluated negatively charged nucleic acid/peptide coacervates^33^ and their interactions with differently packed PC-based membranes. In the absence of Mg^2+^ and cholesterol-functionalised DNA, the coacervates (comprising R_4_ and DNA_12_ in a 1:1 charge ratio) did not exhibit appreciable membrane association on their own, consistent with electrostatic repulsion between the two negatively charged supramolecular structures (Figs. 5e-f and S60). Upon addition of Mg^2+^, however, membrane association became measurable and followed the same compositional trend observed for DNA nanoprobes, with 12:0 PC membranes showing the strongest interaction, followed by 18:1 PC membranes and then 18:0 PC membranes (Figs. 5e-f and S50). These observations indicate that lipid packing, rather than surface charge alone, remains the dominant determinant of membrane recruitment, even in macroscopic, compositionally complex, charged coacervates. Notably, when cholesterol-functionalised DNA was pre-incorporated into the coacervates, its partitioning was reduced relative to the unmodified oligonucleotide (Fig. S61), suggesting that the hydrophobic modification perturbs the interactions that drive probe recruitment within the coacervate phase. Nevertheless, addition of 12:0 PC liposomes rapidly promoted membrane association even in the absence of Mg^2+^ (Figs. S62-S63). In contrast, 18:1 PC and 18:0 PC membranes showed only weak interactions with coacervates (Figs. S62-S63), indicating that a lipid bilayer with the favourable packing properties discussed above can efficiently drive coacervate-membrane association even when only a small fraction of the amphiphilic DNA probe is partitioned into the coacervate.

Together, these results demonstrate that membrane-packing principles can be used to engineer membrane selectivity and bias recruitment toward one population over another, across both simple lipid mixtures and higher-order coacervate-membrane assemblies.

## Conclusions

In this study, we examined how acyl chain length and unsaturation affect the biophysical properties of zwitterionic and anionic phospholipid bilayers, modulating the attachment of amphiphilic DNA nanoprobes to the membrane. By systematically varying acyl chain length, we show that both lipid packing and membrane surface charge govern nanoprobe binding, which can be tuned by regulating the lipid hydrophobic core rather than altering the net charges on the lipid headgroups. While previous studies have largely regulated membrane attachment through external ionic conditions or DNA-construct structural properties, our findings establish lipid acyl-chain chemistry as an additional independent tuning parameter. These chain-length-dependent effects persist when bilayers include anionic lipids, such as fatty acids and, to a lesser extent, phosphatidic acids, further confirming the role of lipid packing, in addition to surface charge, in enabling DNA nanoprobe binding. Because membrane association is enabled by a hydrophobic anchor on the single-strand DNA, unstructured nanoprobes effectively act as simple polyanionic polymers. Consequently, our findings should be agnostic to the DNA sequence used, provided the nanoprobe does not fold into secondary structures that could introduce additional steric factors or interaction modes.

Our results also show that amphiphilic DNA nanoprobes preferentially associate with liquid-phase membranes and exhibit reduced binding to gel-phase membranes. Importantly, the binding response is not simply monotonic with membrane packing or fluidity. Instead, efficient recruitment occurs only within an intermediate packing window, where membranes allow insertion of the cholesterol anchor while remaining fluid enough to permit stable partitioning, retention, and lateral diffusion. In this framework, gel-phase membranes are too rigid and tightly packed to accommodate insertion, whereas fluid membranes are too loosely packed to retain the amphiphilic probe. This membrane-packing-dependent binding relationship provides a design criterion for engineering selective membrane targeting and explains why the strongest interactions occur in tightly packed yet fluid membranes.

Together, our results identify an intermediate membrane-packing regime as optimal for amphiphilic DNA recruitment, providing a design principle in which productive binding requires membranes that are ordered enough to support insertion, yet fluid enough to permit stable retention and lateral diffusion. Orthogonal Raman spectroscopy measurements support these conclusions by independently confirming enhanced DNA-associated signal on tightly packed liquid-phase membranes. Coacervate assays show that the same packing-dependent preference extends to coacervate-membrane interactions. Finally, in line with recent studies comparing DNA nanoprobes carrying different hydrophobic anchors,^11,34^ we identify a strong dependence of membrane interactions on the identity of the lipophilic anchoring moiety. Collectively, our findings outline a framework for designing selective membrane-biomolecule interactions that do not rely exclusively on surface charge but instead on modulating the bilayer hydrophobic core by regulating lipid chain length, membrane phase, and lipophilic anchor identity. These insights have direct implications for bionanotechnology and bottom-up synthetic biology, where controlling and fine-tuning interactions in membrane-nucleic acid platforms are relevant to gene delivery,^35^ vaccine development,^36^ and the design of membrane-active DNA nanostructures.^37,38^ In particular, we anticipate that these design principles will enable the engineering of sophisticated responses in synthetic cells,^39,40^ ranging from membrane-hosted detection^41–43^ to targeted molecular transport^44^ and surface-assisted self-assembly.^45,46^

More broadly, the principles uncovered here, which link the molecular biophysics of the bilayer hydrophobic core to that of DNA nanoprobe binding, likely extend to other amphiphilic macromolecular structures, including lipid-anchored membrane proteins,^47^ SNARE-like membrane fusion systems,^48^ cytoskeletal filaments,^49^ protein fibrils,^50^ and viruses interacting with membranes during cell entry.^51^ In particular, the interactions and lateral distribution of the SNARE complex within membranes are highly sensitive to fatty acids, and these effects are regulated by acyl-chain length and unsaturation.^52–54^ Similarly, acyl-chain length and unsaturation modulate the binding affinity and lipid solvation of peripheral and lipid-anchored membrane proteins,^55,56^ while hydrophobic mismatch, driven by differences in acyl-chain length and transmembrane domain thickness, governs membrane protein insertion, packing, and distribution.^53,57^ Our approach thus provides a mechanistic framework for dissecting and controlling biomolecular interactions with biomimetic and biological membranes, with far-reaching implications for membrane biophysics, cell biology, and bionanotechnology.

## Acknowledgements

The authors acknowledge funding from the Human Frontier Science Program Organization (HFSPO) *via* an Early Career Research Grant (RGY00062/2022, https://doi.org/10.52044/HFSP.RGY00622022.pc.gr.153594, to C.B. and D.K.O.), the ERC (Starting Grant) under the European Union’s Horizon Europe research and innovation programme (GA 101162933 to C.B.), the Leverhulme Centre for Life in the Universe *via* a Joint Collaborative Programme Research Grant (to C.B.), the Federation of European Biochemical Society via a FEBS Excellence Award (to C.B.), the UKRI Future Leaders Fellowship (UKRI2316/G130052 to C.B.), the Agence Nationale de la Recherche *via* an ANR AAPG JCJC 2022 (to C.B.), the University of Strasbourg Institute for Advanced Study (USIAS) *via* a USIAS Fellowship (to C.B.), the Foundation Jean-Marie Lehn, the CSC Graduate School funded by the Agence Nationale de la Recherche (CSC-IGS ANR-17-EURE-0016 for doctoral funding to K.T.M.), the Biotechnology and Biological Sciences Research Council *via* a BBSRC Discovery Fellowship (BB/X010228/1 to R.R.S.), the NSERC *via* a NSERC Discovery Grant (RGPIN 2020-05043 to D.K.O.) and a NSERC Alliance Catalyst Grant (ALLRP 57555822 to D.K.O.).

