## Supporting Information for "Lipid tail chemistry regulates selective membrane interactions with model DNA nanoprobes and DNA-based coacervates"

**Materials and Methods**

**Supporting Table 1**

**Supporting Figures 1 to 63**

**References**

### Materials and Methods

1,2-dilauroyl-*sn*-glycero-3-phosphate (sodium salt) (12:0 PA), 1,2-dioleoyl-*sn*-glycero-3-phosphate (sodium salt) (18:1 PA), 1,2-didecanoyl-*sn*-glycero-3-phosphocholine (10:0 PC),1,2-dilauroyl-*sn*-glycero-3-phosphocholine (12:0 PC), 1,2-dimyristoyl-*sn*-glycero-3-phosphocholine (14:0 PC), 1,2-dipalmitoyl-*sn*-glycero-3-phosphocholine (16:0 PC), 1,2-distearoyl-*sn*-glycero-3-phosphocholine (18:0 PC), 1,2-dimyristoleoyl-*sn*-glycero-3-phosphocholine (Δ9-Cis, 14:1 PC), 1,2-dipalmitoleoyl-*sn*-glycero-3-phosphocholine (Δ9-Cis, 16:1 PC), 1,2-dioleoyl-*sn*-glycero-3-phosphocholine (Δ9-Cis, 18:1 PC) and cholesterol (Chol) were purchased from Avanti Polar Lipids. Capric acid (10:0 FA), lauric acid (12:0 FA), myristic acid (12:0 FA), palmitic acid (16:0 FA), stearic acid (18:0 FA), myristoleic acid (Δ9-Cis, 14:1 FA), palmitoleic acid (Δ9-Cis, 16:1 FA) and oleic acid (Δ9-Cis, 18:1 FA) were purchased from Larodan. All lipids mentioned above were received in powder form and dissolved in chloroform at 50 mM, except for: 12:0 PA, which was dissolved in methanol:chloroform 1:1 at 25 mM; 12:0 FA, which was dissolved in methanol:chloroform 1:1 at 50 mM; and 16:1 PC, which was received already dissolved in chloroform at 10 mg/mL (~13.7 mM). Texas Red 1,2-Dihexadecanoyl-sn-Glycero-3-Phosphoethanolamine Triethylammonium Salt (Texas Red-PE) was purchased from Invitrogen Thermo Fisher Scientific and dissolved in chloroform at 2 mM.

DNA oligonucleotides (ONs) were synthesised using an ABI-394 DNA synthesiser using *N*-benzoyl-dA, *N*-isobutyryl-dG, *N*-acetyl-dC, dT, 4-triazolyl thymidine CED phosphoramidites purchased from ChemGenes. All ONs were synthesised from FAM-CPG 500, 6-isomer, which was purchased from Lumiprobe. Amino-PEG7-alcohol and 1-decylamine were purchased from Combi-Blocks, and oleylamine was purchased from Sigma-Aldrich. *N,N′*-Disuccinimidyl carbonate (DSC) and all other solvents used were purchased from Fisher Scientific.

Modified and unmodified ONs were purchased from IDT or synthesised in-house (SiH). Seven different ONs were used in this study; the structure of the modification is shown only for SiH ONs:

| DNA sequence | Abbreviation | Source |
|---|---|---|
| 5′-Cy3-AGT AGT ATC CAT-TEG-Cholesterol-3′ | Cy3-DNA-chol | IDT |
| 5′-FAM-AGT AGT ATC CAT-TEG-Cholesterol-3′ | FAM-DNA-chol | IDT |
| 5′-Cholesterol-TEG-AGT AGT ATC CAT-FAM-3′ | Chol-DNA-FAM | IDT |
| 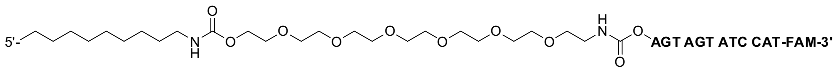  | C10*-DNA-FAM | SiH |
| 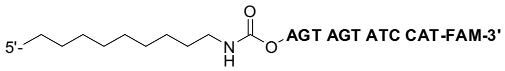  | C10-DNA-FAM | SiH |

| | | |
|---|---|---|
| 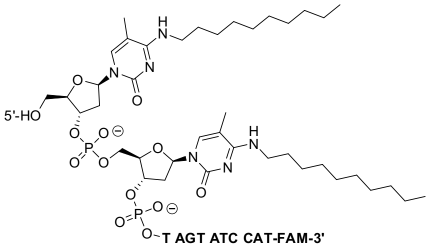 | 2xC10-DNA-FAM | SiH |
| 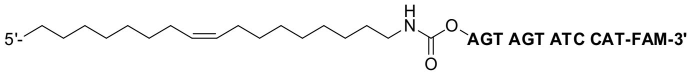 | C18:1-DNA-FAM | SiH |

All single-stranded oligonucleotides were received in dry form, rehydrated with Milli-Q water to a concentration of 100 μM, and diluted to 1 μM for sample preparation. The concentration of μDNA stock solutions was confirmed by measuring their absorbance at 260nm[1] on an IMPLEN Nano UV/Vis Spectrophotometer. Extinction coefficients were calculated using the IDT OligoAnalyzer tool. All stock solutions (lipids or DNA) were aliquoted and kept at -20°C, and thawed to room temperature before use.

Salts, buffers, solvents, and consumables were purchased from Sigma-Aldrich, Waters, and Fisher Scientific and used as received.

**Giant Unilamellar Vesicles (GUVs) formation.** GUVs were formed by electroformation as previously reported.[2] Specifically, 5 μL of 50 mM lipid stock solution was diluted in chloroform to a final volume of 50 μL (5 mM). The diluted solution was spread on the conductive side of an indium tin oxide (ITO) glass slide. When needed, 0.8% Texas Red-PE was added to the diluted lipid solution before spreading it onto the glass slide. The ITO glass slide was covered with aluminium foil and placed in a desiccator for at least 1 hour to ensure complete solvent evaporation. The electroformation chamber was built by depositing a ~1.5 mm hollow polydimethylsiloxane (PDMS) spacer between the lipid-coated ITO glass slide and a non-coated ITO glass slide with the conductive sides facing each other. The hollow PDMS spacer was filled with ~500 μL of sucrose solution (300 mM). When fatty acids were included in the lipid mixture, equimolar amounts of sodium hydroxide (relative to fatty acid content) were added to the solution. The electroformation chamber was connected to a generator (Aim TTi TG315) and subjected to a sinusoidal alternating current with a 2 $V_{p-p}$ voltage amplitude and a 10 Hz frequency for 2 hours, followed by a 2 $V_{p-p}$ voltage amplitude and a 0 Hz frequency for 0.5 hours. When phospholipids with a melting temperature above room temperature were tested, the electroformation process was performed in a fan-ventilated oven at 65°C. At the end of the electroformation process, the solution containing GUVs was collected from the hollow PDMS spacer and stored at room temperature in the dark to avoid photooxidation. The electroformed GUVs solution had a final total lipid concentration of 0.5 mM (quantified by HPLC). GUVs were prepared one day before use.

**Large Unilamellar Vesicles (LUVs) formation.** LUVs were formed by lipid film rehydration followed by extrusion as previously reported.[2] Specifically, a solution of the desired lipid in chloroform was evaporated overnight in an amber vial inside a desiccator. A fluorescent lipid, e.g., Texas Red PE, was added to the solution before evaporation if needed. Half of the required volume of HEPES buffer (5 mM, pH 7.5) was used to rehydrate the lipid film. Note that, to minimise lipid degradation (e. g., hydrolysis, oxidation and polymerisation) catalysed by heat exposure, membranes composed of high-melting-temperature ($T_m$) lipids (14:0, 16:0, 18:0, and 20:0) were simply rehydrated and vortexed but not extruded at temperatures above their $T_m$. When fatty acids were included in the lipid mixture, equimolar amounts of sodium hydroxide (relative to fatty acid content) were added to the buffer solution. The solution was then rehydrated at 65°C for 15 minutes, vigorously vortexed for 1 minute, sonicated for 1 hour, and tumbled for 1 hour, after which the remaining required volume of HEPES buffer was added. The lipid solution was then extruded 9 times through a 200 nm membrane using a mini extruder (Avanti Polar Lipids).

**Confocal microscopy.** The procedure for sample preparation for confocal microscopy was adapted from a previously reported method.[2] Specifically, 20 μL of GUVs (final concentration: 0.1 mM) were added to 80 μL of a solution containing fluorescently-labelled cholesterol-tagged DNA (final concentration: 0.1 μM), HEPES buffer (final concentration: 5 mM), pH 7.5, and $MgCl_2$ as required. The osmolarity of the solution was measured using a Gonotec Osmomat 3000 basic and was maintained equal to the osmolarity in the lumen of the liposomes upon addition of a glucose solution (340 ± 10 mOsm/kg). The solutions containing GUVs and DNA were vortexed for ~3 s and incubated for 1 hour before observation. Experiments were conducted at room temperature unless otherwise specified.

Micrographs were obtained using a Leica TCS SP5 confocal microscope equipped with a 63× oil objective. Cy3 was excited with the DPSS 561 nm laser at 20% intensity, and emission was collected between 575 nm and 639 nm; FAM was excited with the Argon laser at 10% power and 40% intensity, and emission was collected between 502 nm and 530 nm; Texas Red-PE was excited with the DPSS 561 nm laser at 5% intensity, and emission was collected between 601 nm and 651 nm. Micrographs were collected using a 200 Hz scan speed, HyD detector with gain = 500, pinhole = 95.4 μm, 2-frame accumulation for FAM-labelled samples and 2-frame average for Cy3-labelled samples. DNA-liposome binding was quantified using ImageJ.

**Quantitative image analysis and GUV segmentation.** We performed image analysis and fluorescence quantification of GUVs using a custom-built image segmentation and analysis pipeline. The workflow consists of automated two-pass vesicle detection, sub-pixel radial membrane profiling, Pratt circle re-fitting, and numerical intensity integration.

To identify GUVs across varying signal-to-noise ratios without introducing detection bias, we processed images in two sequential passes. Initially, confocal fluorescence micrographs were smoothed using a 2D Gaussian filter ($\sigma = 2$). Vesicle candidates were identified on binarised images via a Circular Hough Transform across a radius range of 25 to 300 pixels. To capture dim GUVs while avoiding duplicate detections, vesicles identified in the first pass were masked, and the remaining image was run through the detection algorithm with a higher sensitivity setting.

For every detected vesicle candidate, 361 radial profile lines (in increments of 1° from 0° to 360°) were cast from an inner radius (0.6×R) to an outer radius (1.1×R) relative to the initial centre of the GUV. Along each radial vector, sub-pixel intensity profiles were sampled and fitted to a 1D Gaussian function.

To correct for imperfect primary segmentation, the set of 361 peak coordinates identified as membrane locations was fitted to a circle. Outlier coordinates deviating by more than 5% from the fitted radius were removed. A final fit was then performed on the remaining membrane locations to determine the refined vesicle centre and radius.

Membrane-associated fluorescence intensities were sampled from membrane locations and integrated across a ±2 pixel window centred on the fitted peak to account for point spread function blurring across the bilayer width. The integrated membrane intensity at every angle $\theta$, denoted as $I(\theta)$, was calculated by normalising the integral over the window width. We defined the total membrane fluorescence for an individual GUV as the mean across valid locations sampled along the perimeter.

After automated segmentation, we visually inspected all images using an interactive curation tool. Images were displayed with contrast enhancement (0.5% saturation limits), colour-matched circle overlays and vesicle identification numbers. We then manually flagged and removed misidentified vesicles, aggregates, or overlapping vesicles from the dataset before statistical analysis.

**Zeta (ζ) potential studies.** The Zeta potential of LUV samples was measured using a Malvern Panalytical Zetasizer Ultra Red.[3] The total lipid concentration in LUV samples used for these measurements was 0.5 mM in 300 mM sucrose in the presence or absence of 5 mM HEPES buffer, pH 7.5. The required amounts of $MgCl_2$ were added to the LUV solutions after liposome formation, and the samples were sonicated for 1 minute before measurement to remove air bubbles. The high-voltage ZEN1010 cuvette was used for the measurements, with the solvent dielectric constant set to 78.4, viscosity to 1.33 mPa·s, and refractive index to 1.35. Measurements were recorded at 25°C with a 120 s equilibration time. Each sample was measured at least 3 times.

**Lipid packing studies.** Lipid packing was assessed by measuring the fluorescence spectrum of the dye Laurdan[4] (6-dodecanoyl-2-dimethylaminonaphthalene) upon its incorporation into the membrane, using a spectrofluorometer (Jasco FP-8350). Laurdan general polarisation (GP) was calculated by the following equation:

$$GP = \frac{I_{430} - I_{500}}{I_{430} + I_{500}}$$

where $I_{430}$ and $I_{500}$ refer to the fluorescence intensity of Laurdan at 430 nm and 500 nm, respectively.

LUVs used for these measurements were prepared with 1% (m/m) Laurdan at a total lipid concentration of 0.5 mM in 300 mM sucrose containing 5 mM HEPES buffer, pH 7.5. Laurdan was excited at 350 nm, and the emission spectrum was recorded between 400 nm and 520 nm with a 1000 nm/min scan speed, 350 V voltage and 5 nm excitation and emission bandwidths. All experiments were performed at 25ºC.

**Membrane fluidity measurements.** Membrane fluidity was assessed through fluorescence anisotropy of the dye DPH[5] (1,6-diphenyl-1,3,5-hexatriene) upon incorporation into the membrane, via fluorescence polarisation measurements using a Plate Reader (BMG Labtech CLARIOstarplus). LUVs used for these measurements were prepared with 1% (m/m) DPH at a total lipid concentration of 0.5 mM in 300 mM sucrose containing 5 mM HEPES buffer, pH 7.5. DPH was excited at 360 nm with a dichroic filter at 410 nm, and emission was recorded at 450 nm with Gain A = 1351, Gain B = 1339, and a focal height of 1.5 mm. Each sample was measured at least 4 times.

**Single-particle Raman spectroscopy.** LUVs were prepared at a total lipid concentration of 0.5 mM in 5 mM HEPES buffer, pH 7.5 as described above. The lipid solutions were extruded 21 times through a 200 nm membrane. The procedure for sample preparation for single-particle Raman spectroscopy was adapted from the protocol reported above for confocal microscopy. Specifically, aliquots in a total volume of 100 µL comprised LUVs (final concentration: 0.5 mM), fluorescently-labelled cholesterol-tagged DNA (final concentration: 0.1 µM), HEPES buffer (final concentration: 5 mM), pH 7.5, and $MgCl_2$ as required. The solutions containing LUVs and DNA were vortexed for ~3 s and incubated for 1 hour before observation. Experiments were conducted at room temperature unless otherwise specified.

Single-particle Raman measurements were performed using the SPARTA AGIS I system. A 100 µL aliquot of each sample was deposited onto a SPARTA sample slide, and individual particles were optically trapped and measured for 10 s. A laser shut-off time of 1 s was used between acquisitions to allow particle diffusion and ensure independent measurements. Background spectra were collected using a mixture of 85 µL 5 mM HEPES buffer, pH 7.5, and 15 µL Milli-Q water. Spectral data were analysed using the SPARTA BioDiscovery software, where individual spectra were baseline-corrected and normalised to the integrated area. DNA binding was quantified using the Raman peak corresponding to the symmetric deformation of methyl groups in the Cy3-labelled DNA probe, centred around ~1395 $cm^{-1}$, and the Raman peak corresponding to the symmetric stretching vibration of the C-N bonds in the quaternary ammonium moiety, centred around ~710 $cm^{-1}$ and characteristic of the gauche conformation of the O-C-C-$N^+$ backbone of the choline headgroup. All experiments were performed in at least 3 biological replicates.

**Oligonucleotide synthesis, purification and characterisation.** DNA oligonucleotides (ONs) were assembled using standard reagents and manufacturer protocols for the ABI 394 DNA synthesiser. DMTr-removal reagent consisted of 3% trichloroacetic acid in dichloromethane, the activator consisted of 0.25 M 5-ethylthio tetrazole in acetonitrile, the oxidiser consisted of a 0.02 M solution of iodine in 8:16:76 pyridine:water:tetrahydrofuran, and the capping reagents consisted of (Cap A) a solution of 10:10:80 acetic anhydride:pyridine:tetrahydrofuran and (Cap B) a 10% (v/v) solution of *N*-methylimidazole in tetrahydrofuran. All ONs were deprotected from the solid support using a 4:1 mixture of 25% ammonium hydroxide/ethanol (total volume of 500 µL to 1 mL) for 17 h at 55°C.

Deprotected ONs were purified by Strong Anion-Exchange (SAX) chromatography using a DNApac™ PA200 column with a Vanquish™ high-performance liquid chromatography (HPLC) system with a gradient of 0-37%, 0-50%, 0-75% or 0-100% elution solvent (1 M NaCl in 10% MeCN, pH 7.61, 50 mM Tris buffer) and desalted using a Sep-Pak C18 classic cartridge. The Sep-Pak C18 cartridge was conditioned with 10 mL of MeCN, 50% MeCN/ultra-pure water (water with 18.2 MΩ-cm ionic purity), and 100 mM NaOAc (pH 7). The purified oligo was diluted to at least 2% MeCN (1:4 with ultra-pure water) and loaded onto the column at least twice. The bound oligo was washed with ultra-pure water (~25 mL) and eluted from the column with 4 mL of 50% MeCN in ultra-pure water, and then concentrated.

UV measurements of newly synthesised ONs were taken at 260 nm using an Agilent BioTek Epoch Microplate Spectrophotometer. At least three readings were taken for a given sample and averaged to a single value. The value was corrected by a blank measurement (averaged from at least three readings). Concentrations were calculated using the Beer-Lambert equation (molar extinction coefficients were estimated using the IDT OligoAnalyzer™ Tool).

Low-resolution mass spectrometry of oligonucleotides was also carried out using ThermoFisher Scientific LCQ Fleet Ion Trap with Ion Max-S Source Housing and Ion Max ESI Source. The mass spectrometer was operated in negative-ion detection mode from 400 to 2000 m/z. The oligonucleotide samples were solubilised in a 1:1 mixture of MeCN:ultra-pure water before direct injection into the instrument. The scans were recorded for 0.5 min and averaged to give the results.

**1-Decylamine modification with spacer.** First, 1 µmol of the ON strand was placed in a 1 µmol standard twist ABI column. A saturated solution of DSC (~30 mg) was made in a 1:1 pyridine (Py):MeCN mixture (1.6 mL) in a 2 mL Eppendorf tube. The solution was vortexed vigorously for 15 seconds and flowed through the column 4-5 times using disposable syringes (3 mL) attached to either side of the column. After 2 hours, the solution was removed from the column, and the CPG was subsequently washed with anhydrous MeCN (10 mL). In a 2 mL Eppendorf tube, 8 mg of amino-PEG7-alcohol was dissolved in a 1:1 Py:MeCN mixture (500 µL, 50 mM). The solution was then vortexed vigorously and flowed through the column 4-5 times using disposable syringes (1 mL) attached to either side of the column. After 30 min, the solution was removed from the column, and the CPG was subsequently washed with DMF, acetone, and MeCN (10 mL each). A saturated solution of DSC (~30 mg) was prepared in a 1:1 Py:MeCN mixture (1.6 mL) in a 2 mL Eppendorf tube. The solution was vortexed vigorously for 15 seconds and flowed through the column 4-5 times using disposable syringes (3 mL) attached to either side of the column. After 2 hours, the solution was removed from the column, and the CPG was subsequently washed with anhydrous MeCN (10 mL). In a 2 mL Eppendorf tube, 10 µL of 1-decylamine was dissolved in a 1:1 Py:MeCN mixture (500 µL, 100 mM). The solution was then vortexed vigorously and flowed through the column 4-5 times using disposable syringes (1 mL) attached to either side of the column. After 30 min, the solution was removed from the column, and the CPG was subsequently washed with DMF, acetone, and MeCN (10 mL each). The strand was deprotected as described above to produce C10-DNA-FAM.

**1-Decylamine modification without spacer.** First, 1 µmol of the ON strand was placed in a 1 µmol standard twist ABI column. A saturated solution of DSC (~30 mg) was prepared in a 1:1 Py:MeCN mixture (1.6 mL) in a 2 mL Eppendorf tube. The solution was vortexed vigorously for 15 seconds and flowed through the column 4-5 times using disposable syringes (3 mL) attached to either side of the column. After 2 hours, the solution was removed from the column, and the CPG was subsequently washed with anhydrous MeCN (10 mL). In a 2 mL Eppendorf tube, 10 µL of 1-decylamine was dissolved in a 1:1 Py:MeCN mixture (500 µL, 100 mM). The solution was then vortexed vigorously and flowed through the column 4-5 times using disposable syringes (1 mL) attached to either side of the column. After 30 min, the solution was removed from the column, and the CPG was subsequently washed with DMF, acetone, and MeCN (10 mL each). The strand was deprotected as described above to produce C10*-DNA-FAM.

**Oleylamine modification without spacer.** First, 1 µmol of the ON strand was placed in a 1 µmol standard twist ABI column. A saturated solution of DSC (~30 mg) was made in a 1:1 Py:MeCN mixture (1.6 mL) in a 2 mL Eppendorf tube. The solution was vortexed vigorously for 15 seconds and flowed through the column 4-5 times using disposable syringes (3 mL) attached to either side of the column. After 2 hours, the solution was removed from the column, and the CPG was subsequently washed with anhydrous MeCN (10 mL). In a 2 mL Eppendorf tube, 16.5 µL of oleylamine was dissolved in a 1:1 Py:MeCN mixture (500 µL, 100 mM). The solution was then vortexed vigorously and flowed through the column 4-5 times using disposable syringes (1 mL) attached to either side of the column. After 30 min, the solution was removed from the column, and the CPG was subsequently washed with DMF, acetone, and MeCN (10 mL each). The strand was deprotected as described above to produce the ON C18*-DNA-FAM.

**1-Decylamine modification through the nucleobase.** The ON was synthesised as described above, with the last two additions to the ON strand being 4-triazolyl thymidine CED phosphoramidite, which were coupled for 10 minutes. Once the synthesis was completed, 0.5

μmol of the ON strand was placed in a 1.5 mL screw cap vial with an O-ring. 16 μL of 1-aminodecane (0.08 mmol) in 200 μL of anhydrous MeCN was added, and the vial was placed at 55 °C for 17 h. After deprotection, the solution was adjusted to pH 7-8 with 1 M acetic acid, then desalted (as described above). This procedure produces the ON 2xC10*-DNA-FAM.

**Principal Component Analysis.** Principal Component Analysis (PCA) is a dimensionality-reduction algorithm used to reduce the four-dimensional membrane property data (including melting temperature, surface charge, Laurdan *GP*, and DPH anisotropy) to two-dimensional principal components. The first principal component (PC1) captures most of the dataset's variance and is calculated using least-squares linear regression to define the direction of the data trend. The second principal component (PC2) is orthogonal to PC1 and represents variation along PC1. The data can then be plotted against the principal components as axes, allowing data to be grouped and making it easier to identify similarities and differences in the dataset. The analysis was carried out in Python using the scikit-learn PCA package.

Before PCA, data for membrane fluidity, packing, surface charge, and $T_m$ were standardised (centred and normalised). This approach ensured that variables with different physical units and numerical scales contributed equally to the components, preventing high-variance parameters from disproportionately biasing the clustering.

Principal component extraction showed that PC1 was strongly influenced by lipid packing (loading = 1.01) and membrane fluidity (loading = 0.96), while PC2 was primarily driven by $T_m$ (loading = 0.46).

**Coacervate formation.** The coacervates used were composed of $R_4$ and $DNA_{12}$ ($DNA_{12}$ = ACTGACTGACTG) at a 1:1 charge ratio, where both components were added at a 10 mM charge concentration, giving a final $R_4$ concentration of 2.5 mM and a final $DNA_{12}$ concentration of 0.83 mM. The coacervates were fluorescently labelled with 0.5 μM FAM-$DNA_{12}$. The coacervates were prepared by mixing MilliQ water, HEPES buffer (25 mM final concentration, pH = 7.5) and $DNA_{12}$, followed by $R_4$ and the fluorescent oligonucleotide, with gentle mixing to induce coacervation.

**Confocal microscopy of coacervates.** A mixture of 2 μL of coacervates, 2 μL of DLPC GUV solution, 2 M glucose solution (final concentration: 300 mM), 500 mM HEPES buffer (final concentration: 25 mM) and $MgCl_2$ as required, was mixed in an Eppendorf tube and directly transferred to a PVA-coated glass slide.

Confocal imaging was performed using a Leica TCS SP5 microscope with a HCX PLAPO 63×/1.40 NA oil-immersion objective. A DPSS 561 laser was used to excite the Texas Red-PE, and an Argon laser (30% power) was used to excite the DNA-FAM. The Texas Red-PE membrane dye was excited at 561 nm at 10% power, and the fluorescence emission was collected between 622 nm and 672 nm using a PMT 4 detector. The (chol-)DNA-FAM dye was excited at 488 nm with 15% power, and the fluorescence emission was collected between 502 nm and 554 nm using a PMT-3 detector. The images were taken at 8-bit resolution, 1400 Hz frame rate, 3-frame average, and 6× zoom. The two channels were captured in a sequential scan to avoid signal overlap between the two dyes.

**Calculation of liposome-coacervate overlap percentage.** All images were processed and analysed by ImageJ. We treated the fluorescence images from the liposome and coacervate channels as separate images. Each converted each channel to a binary mask using the Make Binary tool. The Gaussian Blur Filter was applied to each image (σ = 5.00), and the Threshold tool was used to produce binary approximations of the images (below = ~85%, above = 0%). The product of these two binary masks was calculated using the Multiply tool. Similarly, the sum of both masks was calculated using the Add tool. The pixel intensity of the resulting images was obtained using the Measure tool. The channel overlap percentage was then calculated by the following equation:

$$overlap\ percentage\ = \frac{C\cap L}{C\cup L} = \frac{C\times L}{(C+L)-(C\times L)} \times 100,$$

where $C \cap L$ is given by the pixel intensity of the product of the liposome and coacervate channels, and $C \cup L$ is given by the pixel intensity of the sum of the liposome and coacervate channels with $C \cap L$ subtracted.

*Table S1 - Lipid library tested in this study. $T_m$ values were either found on the Avanti Polar Lipids website (https://www.avantiresearch.com/en-gb/support-hub/physical-properties/phase-transition-temps) or measured in-house by Differential Scanning Calorimetry (DSC).*

| | $T_m$ (°C) |
|---|---|
| 12:0 PC | -2 |
| 14:0 PC | 24 |
| 16:0 PC | 41 |
| 18:0 PC | 55 |
| 20:0 PC | 66 |
| 14:1 PC (Δ9-cis) | -47 |
| 16:1 PC (Δ9-cis) | -36 |
| 18:1 PC (Δ9-cis) | -17 |
| 20:1 PC (Δ11-cis) | -4 |
| 22:1 PC (Δ13-cis) | 14 |
| 18:0-18:1 PC (Δ9-cis) | 6 |
| 18:1-18:0 PC (Δ9-cis) | 9 |
| 18:1 PC (Δ6-cis) | 1 |
| 18:1 (Δ9-trans) PC | 12 |

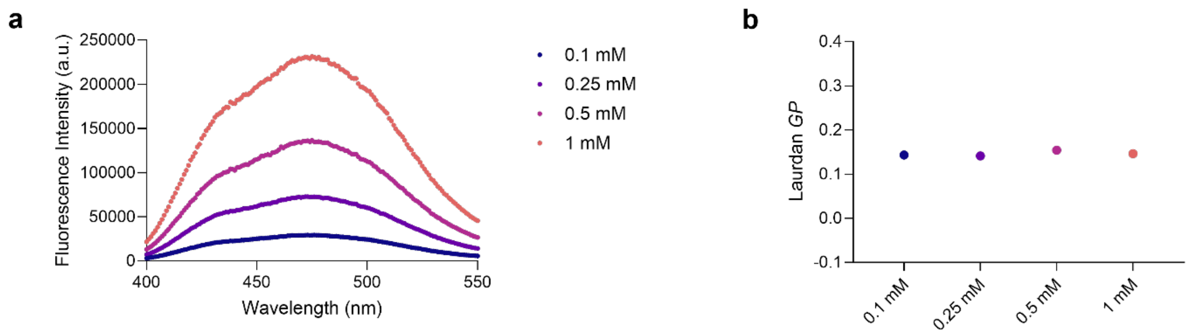


*Figure S1 - Fluorescence spectra (a) and Laurdan GP values (b) obtained for 12:0 PC LUVs at different total lipid concentrations (from 0.1 mM to 1 mM). Data represent mean values with error bars indicating standard deviations from n ≥ 3 independent measurements.*

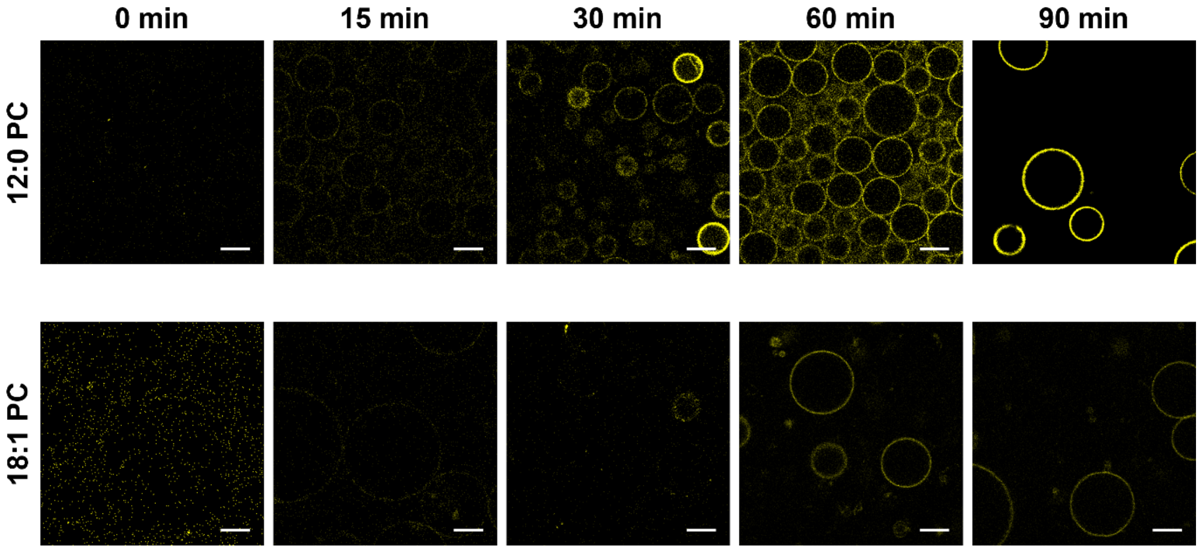


*Figure S2 - Confocal micrographs showing membrane binding of amphiphilic Cy3-DNA nanoprobes (yellow) at different timepoints in the presence of 0.5 mM $Mg^{2+}$. Scale bar: 10 µm. n ≥ 3 independent measurements.*

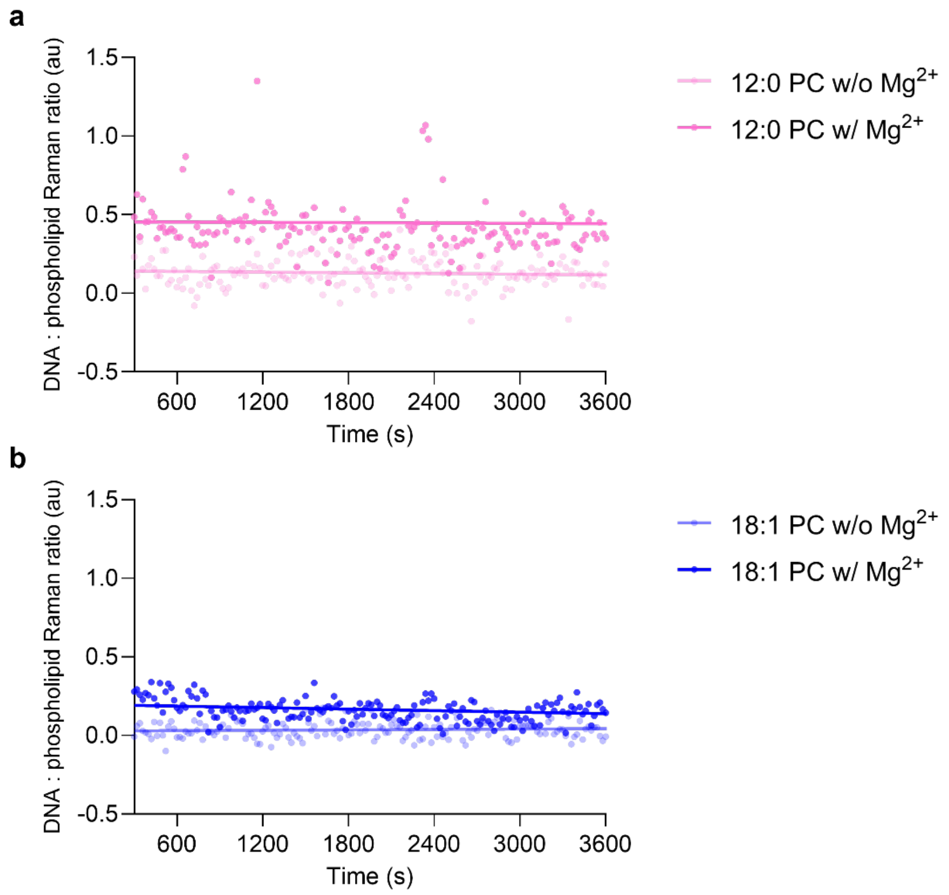


*Figure S3 - Time course of amphiphilic DNA nanoprobe binding to 12:0 PC (a) and 18:1 PC (b) LUVs in the presence or absence of 0.5 mM $Mg^{2+}$. Quantification of the Cy3-DNA Raman signal of individual particles for distinct membranes is normalised to the choline signal of PC lipids (1385-1405/700-730 $cm^{-1}$). n ≥ 3 independent measurements.*

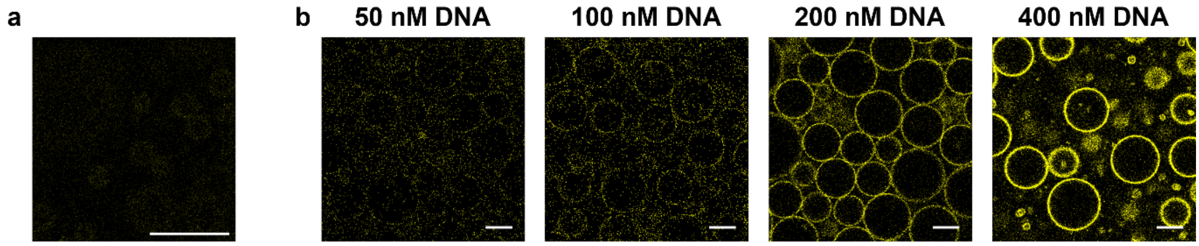


*Figure S4 - Confocal micrographs showing membrane binding (and lack thereof) of amphiphilic Cy3-DNA nanoprobes (yellow) without the cholesterol anchor (DNA concentration: 100 nM) and with the cholesterol anchor at different DNA concentrations in the presence of 0.5 mM $Mg^{2+}$. Scale bar: 10 µm. n ≥ 3 independent measurements.*

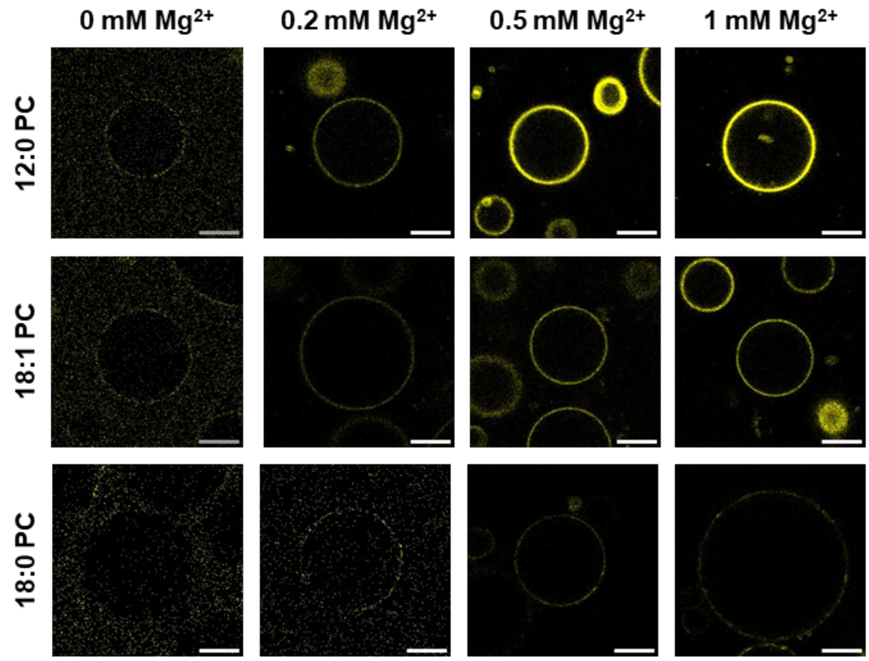


*Figure S5 - Confocal micrographs showing membrane binding of amphiphilic Cy3-DNA nanoprobes (yellow) at increasing concentrations of $MgCl_2$. Scale bar: 10 µm. n ≥ 3 independent measurements.*

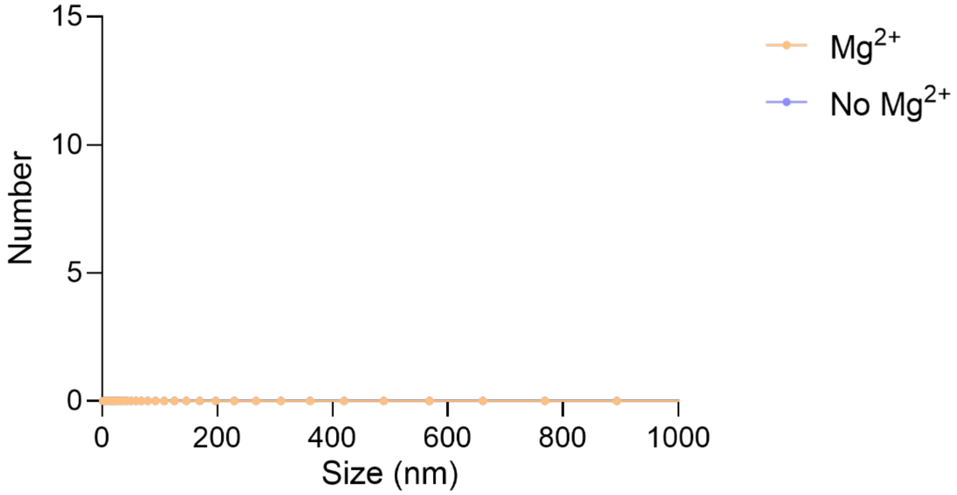


*Figure S6 - Dynamic light scattering profiles (by number) for amphiphilic Cy3-DNA nanoprobes in the presence and absence of 0.5 mM $Mg^{2+}$. Data represent mean values with error bars indicating standard deviations from n ≥ 3 independent measurements.*

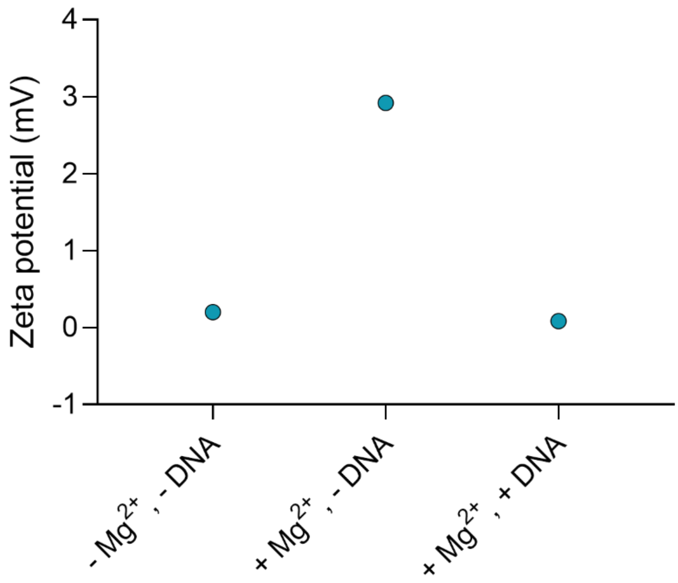


*Figure S7 - Zeta potential values for 12:0 PC LUVs in the presence and absence of amphiphilic DNA nanoprobes and $Mg^{2+}$. Data represent mean values with error bars indicating standard deviations from $n \geq 3$ independent measurements.*

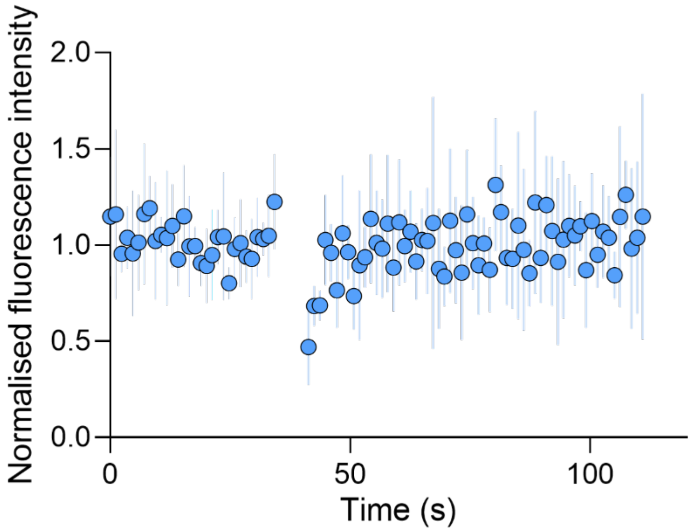


*Figure S8 - FRAP profiles of 12:0 PC GUVs in the presence of amphiphilic Cy3-DNA nanoprobes and 0.5 mM $Mg^{2+}$. Data represent mean values with error bars indicating standard deviations from n ≥ 5 independent measurements.*

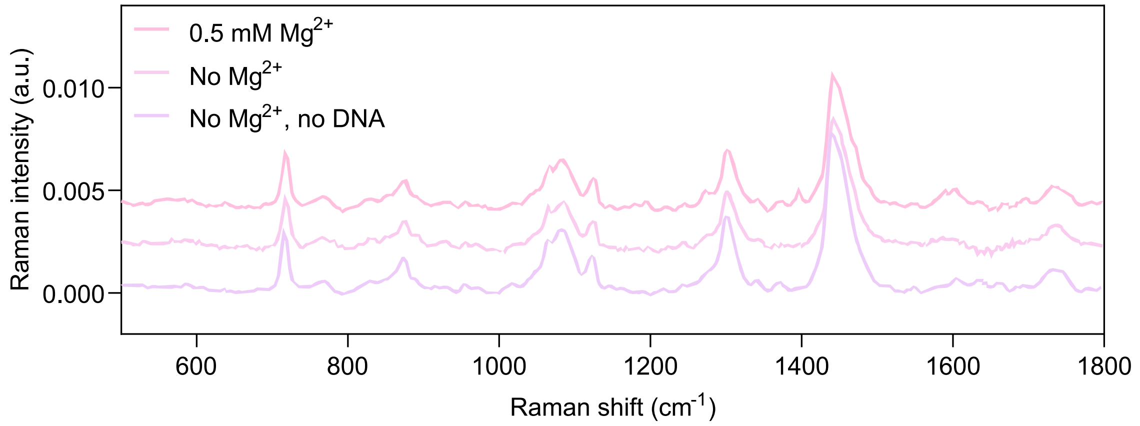


*Figure S9 - Raman spectra (median ± IQR, n ≥ 893) obtained from individual LUVs comprising 12:0 PC in the absence and presence of Cy3-labelled cholesterol-tagged DNA (with and without $Mg^{2+}$). The Cy3 peak region (1385-1405 $cm^{-1}$) is highlighted to demonstrate the detection of amphiphilic DNA binding to LUVs. n ≥ 3 independent measurements.*

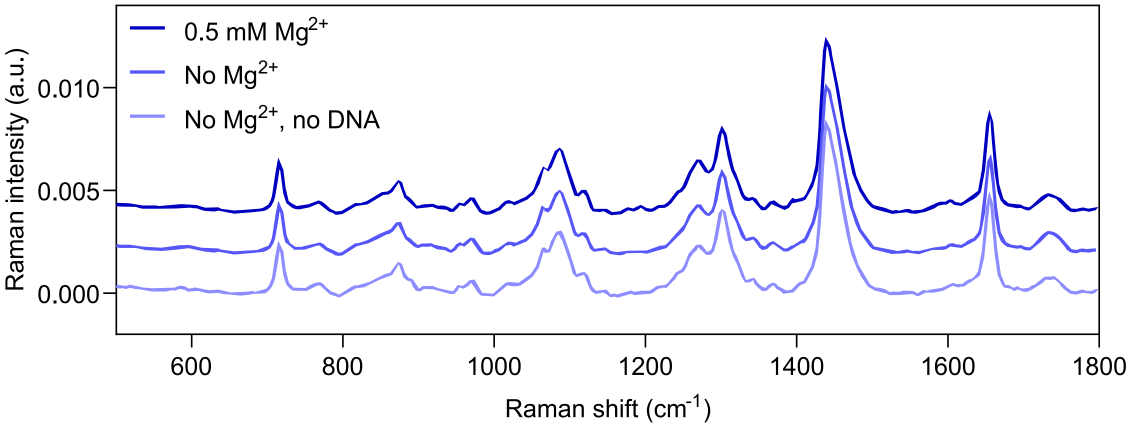


*Figure S10 - Raman spectra (median ± IQR, n ≥ 893) obtained from individual LUVs comprising 18:1 PC in the absence and presence of Cy3-labelled cholesterol-tagged DNA (with and without $Mg^{2+}$). The Cy3 peak region (1385-1405 $cm^{-1}$) is highlighted to demonstrate the (limited) detection of amphiphilic DNA binding to LUVs. n ≥ 3 independent measurements.*

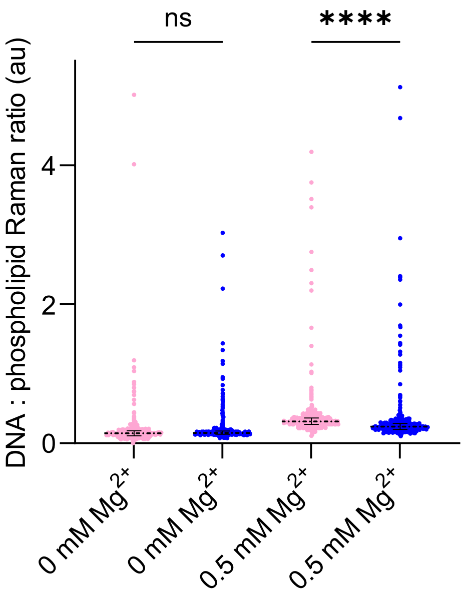


*Figure S11 - Quantification of the Cy3-DNA Raman signal of individual particles for distinct membranes in the absence or presence of $Mg^{2+}$, normalised to the choline signal of PC lipids (1385-1405/700-730 $cm^{-1}$) (median ± IQR, n ≥ 893). Full-scale picture of Figure 1.*

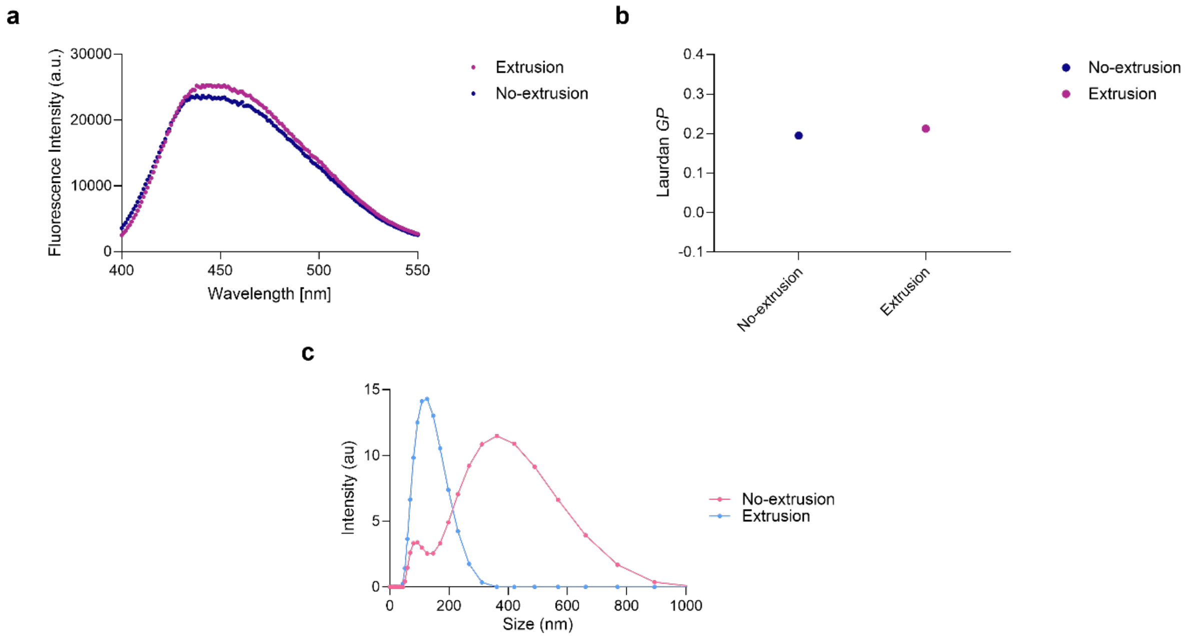


*Figure S12 - Fluorescence spectra (a) and Laurdan GP values (b) obtained for 14:0 PC LUVs with and without extrusion. Dynamic light scattering profiles (by intensity) for 14:0 PC LUVs with and without extrusion. Data represent mean values with error bars indicating standard deviations from n ≥ 3 independent measurements.*

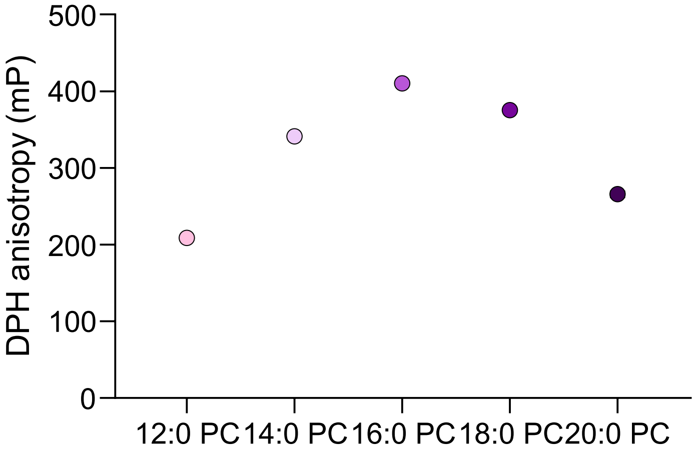


*Figure S13 - DPH anisotropy measured via fluorescence polarisation (mP) for lipid membranes composed of 12:0 PC, 14:0 PC, 16:0 PC, 18:0 PC and 20:0 PC. Data represent mean values with error bars indicating standard deviations from n ≥ 3 independent measurements.*

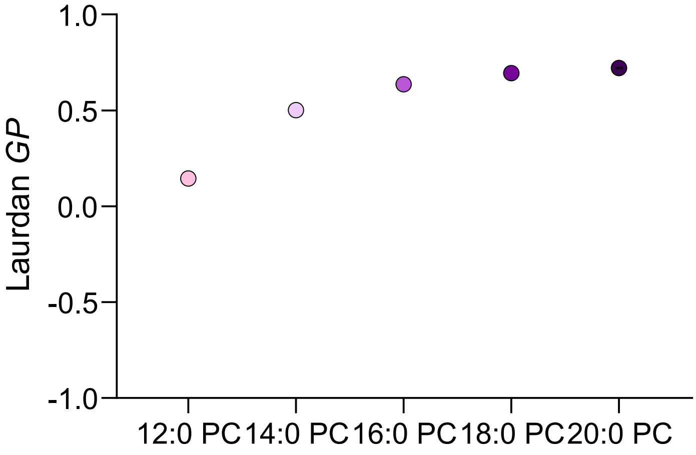


*Figure S14 - Laurdan GP for lipid membranes composed of 12:0 PC, 14:0 PC, 16:0 PC, 18:0 PC or 20:0 PC. Data represent mean values with error bars indicating standard deviations from n ≥ 3 independent measurements.*

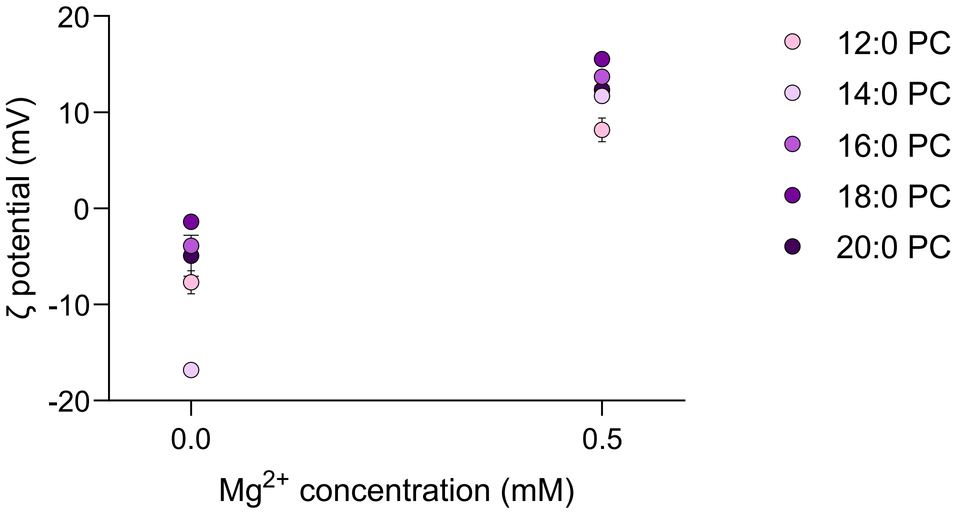


*Figure S15 - Zeta potential as a function of $MgCl_2$ concentration for lipid membranes composed of 12:0 PC, 14:0 PC, 16:0 PC, 18:0 PC or 20:0 PC. Data represent mean values with error bars indicating standard deviations from n ≥ 3 independent measurements.*

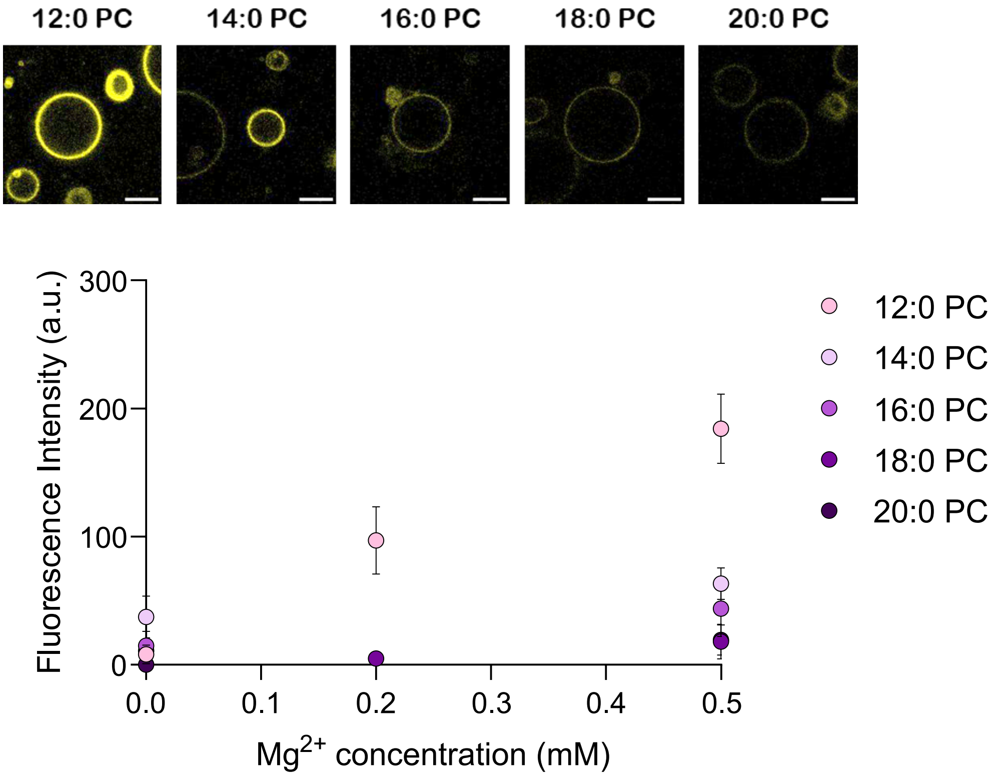


*Figure S16 - Fluorescence intensity of Cy3-labelled DNA-cholesterol conjugates (yellow) as a function of $MgCl_2$ concentration for lipid membranes composed of 12:0 PC, 14:0 PC, 16:0 PC, 18:0 PC or 20:0 PC. Membrane-associated fluorescence signal was quantified from confocal microscopy images (representative confocal micrographs are shown at 0.5 mM $Mg^{2+}$, scale bars = 20 µm). Data represent mean values with error bars indicating standard deviations from $n \geq 3$ independent measurements.*

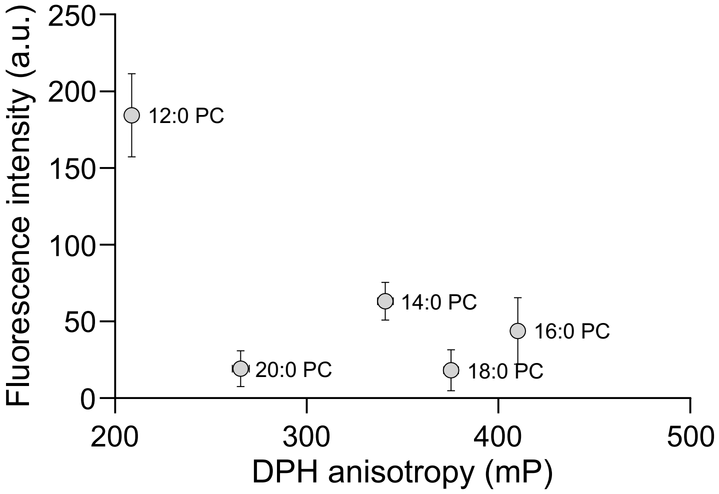


*Figure S17 - Fluorescence intensity of Cy3-labelled DNA-cholesterol conjugates at 0.5 mM $Mg^{2+}$ concentration as a function of DPH anisotropy, which correlates with membrane fluidity, for lipid membranes composed of 12:0 PC, 14:0 PC, 16:0 PC, 18:0 PC or 20:0 PC. Membrane-associated fluorescence signal was quantified from confocal microscopy images. Data represent mean values with error bars indicating standard deviations from n ≥ 3 independent measurements.*

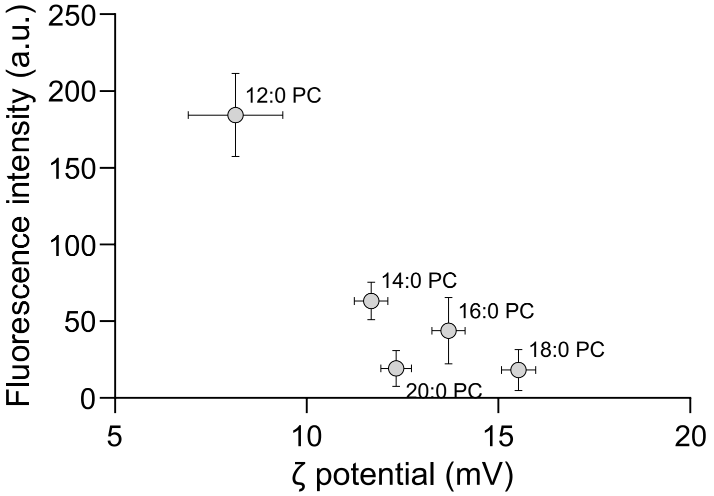


*Figure S18 - Fluorescence intensity of Cy3-labelled DNA-cholesterol conjugates at 0.5 mM $Mg^{2+}$ concentration as a function of Zeta potential, which correlates with membrane surface charge, for lipid membranes composed of 12:0 PC, 14:0 PC, 16:0 PC, 18:0 PC or 20:0 PC. Membrane-associated fluorescence signal was quantified from confocal microscopy images. Data represent mean values with error bars indicating standard deviations from n ≥ 3 independent measurements.*

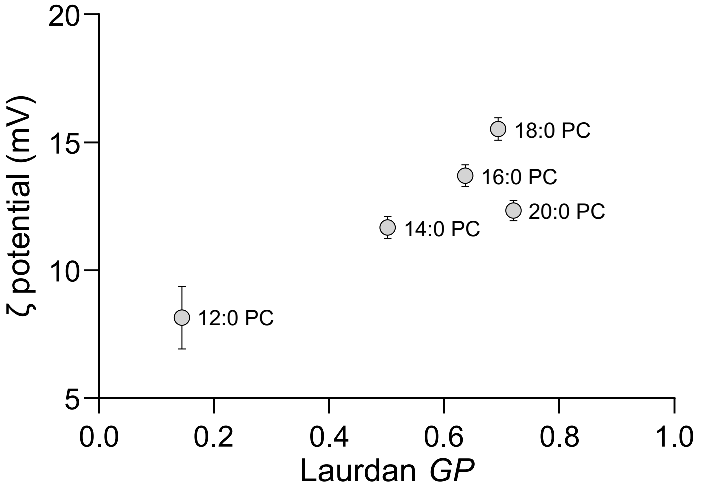


*Figure S19 - Zeta potential, which correlates with membrane surface charge, at 0.5 mM $Mg^{2+}$ concentration as a function of Laurdan GP for lipid membranes composed of 12:0 PC, 14:0 PC, 16:0 PC, 18:0 PC or 20:0 PC. Data represent mean values with error bars indicating standard deviations from n ≥ 3 independent measurements.*

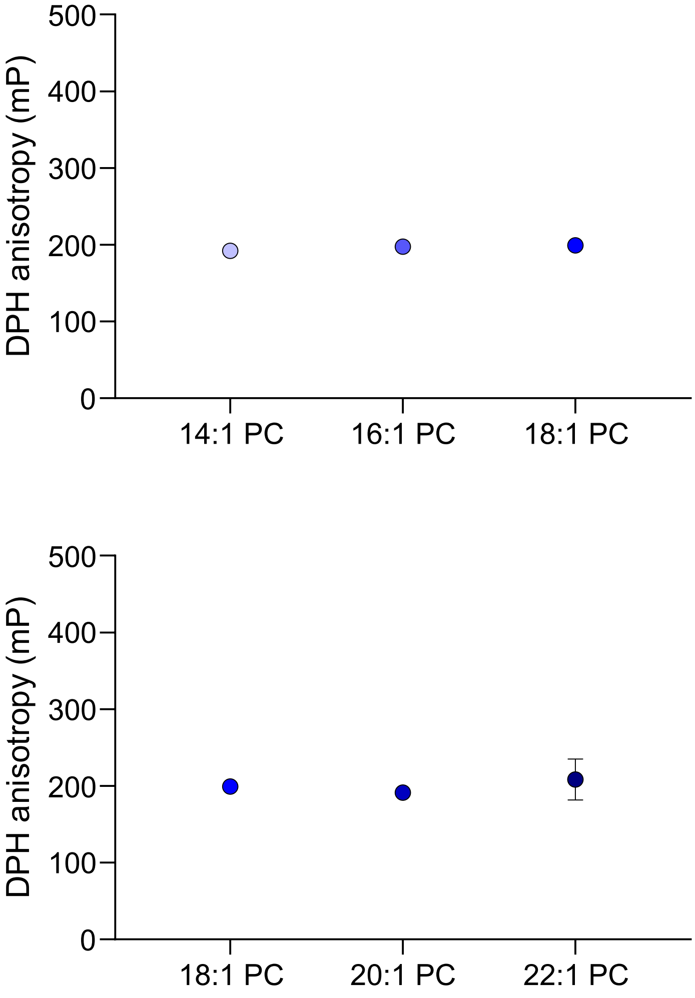


*Figure S20 - DPH anisotropy for lipid membranes composed of (top) 14:1 (Δ9) PC, 16:1 (Δ9) PC and 18:1 (Δ9) PC, and (bottom) 18:1 (Δ9) PC, 20:1 (Δ11) PC and 22:1 (Δ13) PC. Data represent mean values with error bars indicating standard deviations from n ≥ 3 independent measurements.*

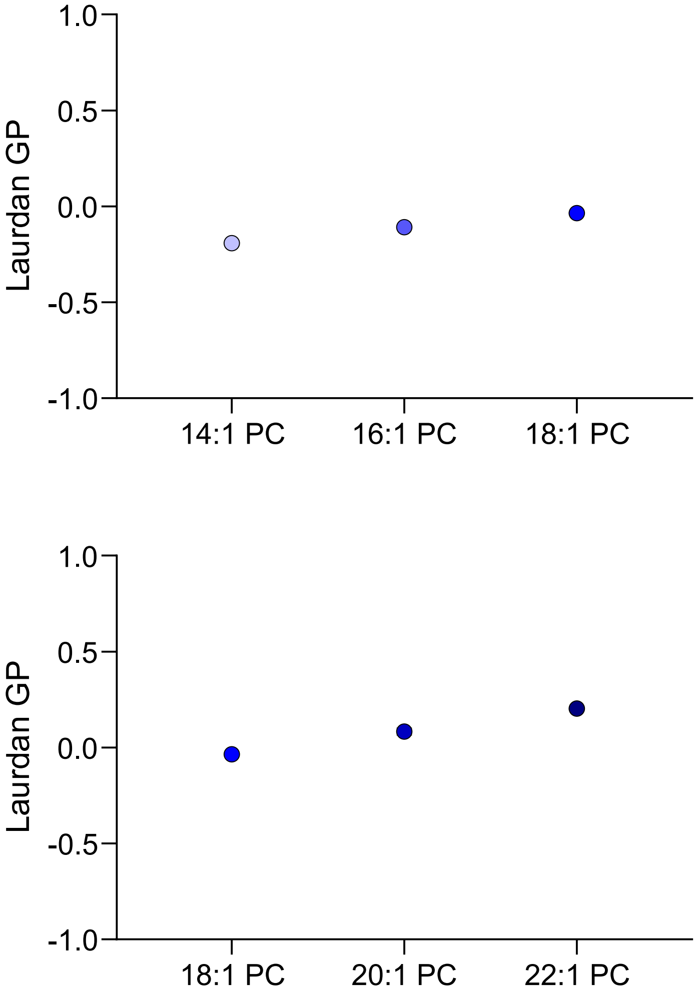


*Figure S21 - Laurdan GP for lipid membranes composed of (top) 14:1 (Δ9) PC, 16:1 (Δ9) PC and 18:1 (Δ9) PC, and (bottom) 18:1 (Δ9) PC, 20:1 (Δ11) PC and 22:1 (Δ13) PC. Data represent mean values with error bars indicating standard deviations from $n \geq 3$ independent measurements.*

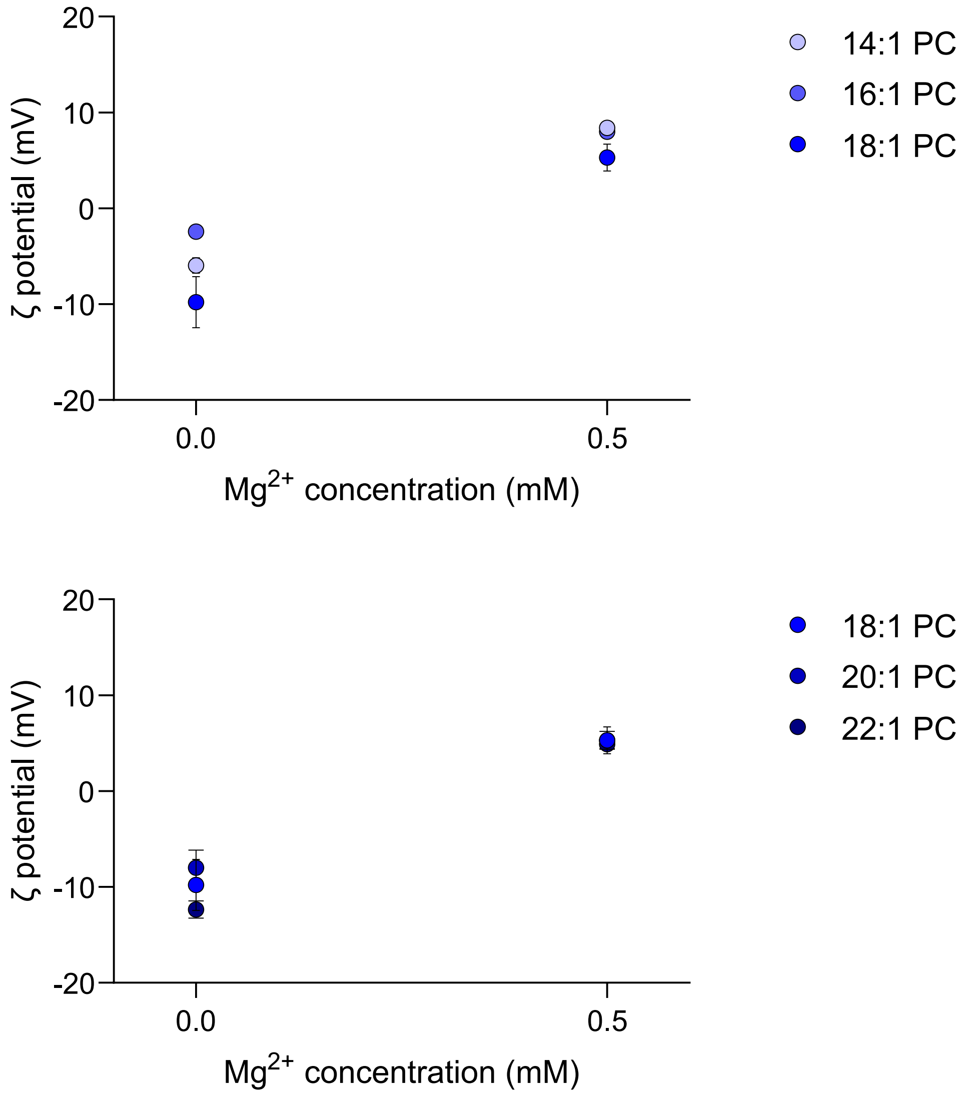


*Figure S22 - Zeta potential as a function of $MgCl_2$ concentration for lipid membranes composed of (top) 14:1 (Δ9) PC, 16:1 (Δ9) PC and 18:1 (Δ9) PC, and (bottom) 18:1 (Δ9) PC, 20:1 (Δ11) PC and 22:1 (Δ13) PC. Data represent mean values with error bars indicating standard deviations from n ≥ 3 independent measurements.*

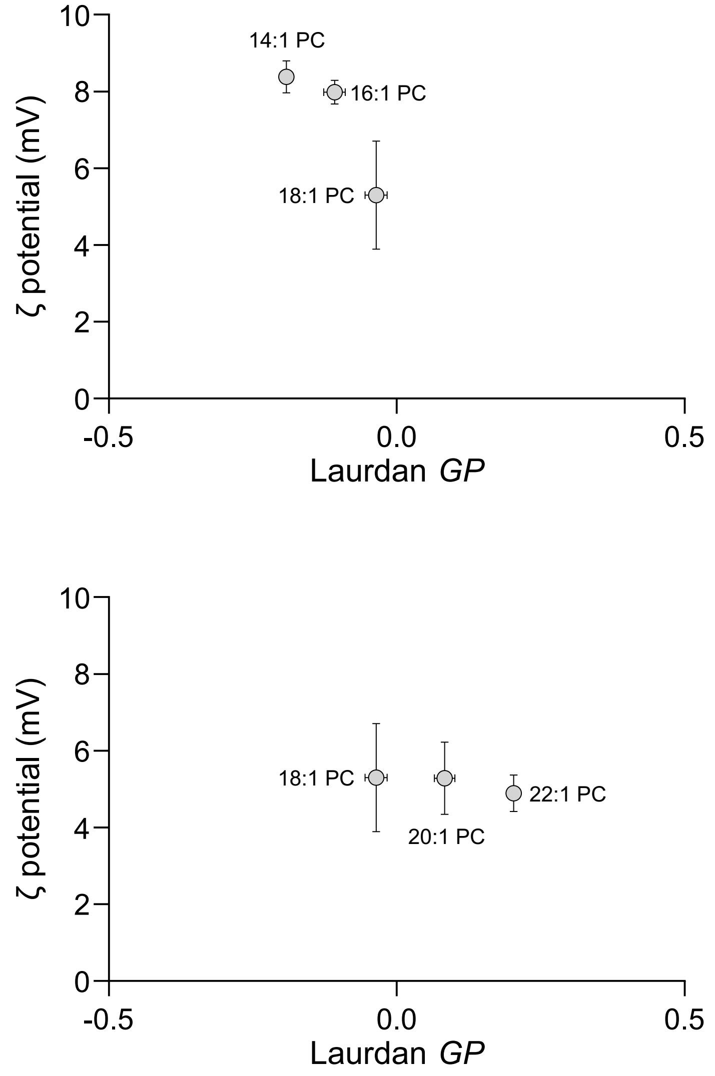


*Figure S23 - Zeta potential, which correlates with membrane surface charge, at 0.5 mM $Mg^{2+}$ concentration as a function of Laurdan GP for lipid membranes composed of (top) 14:1 (Δ9) PC, 16:1 (Δ9) PC and 18:1 (Δ9) PC, and (bottom) 18:1 (Δ9) PC, 20:1 (Δ11) PC and 22:1 (Δ13) PC. Data represent mean values with error bars indicating standard deviations from n ≥ 3 independent measurements.*

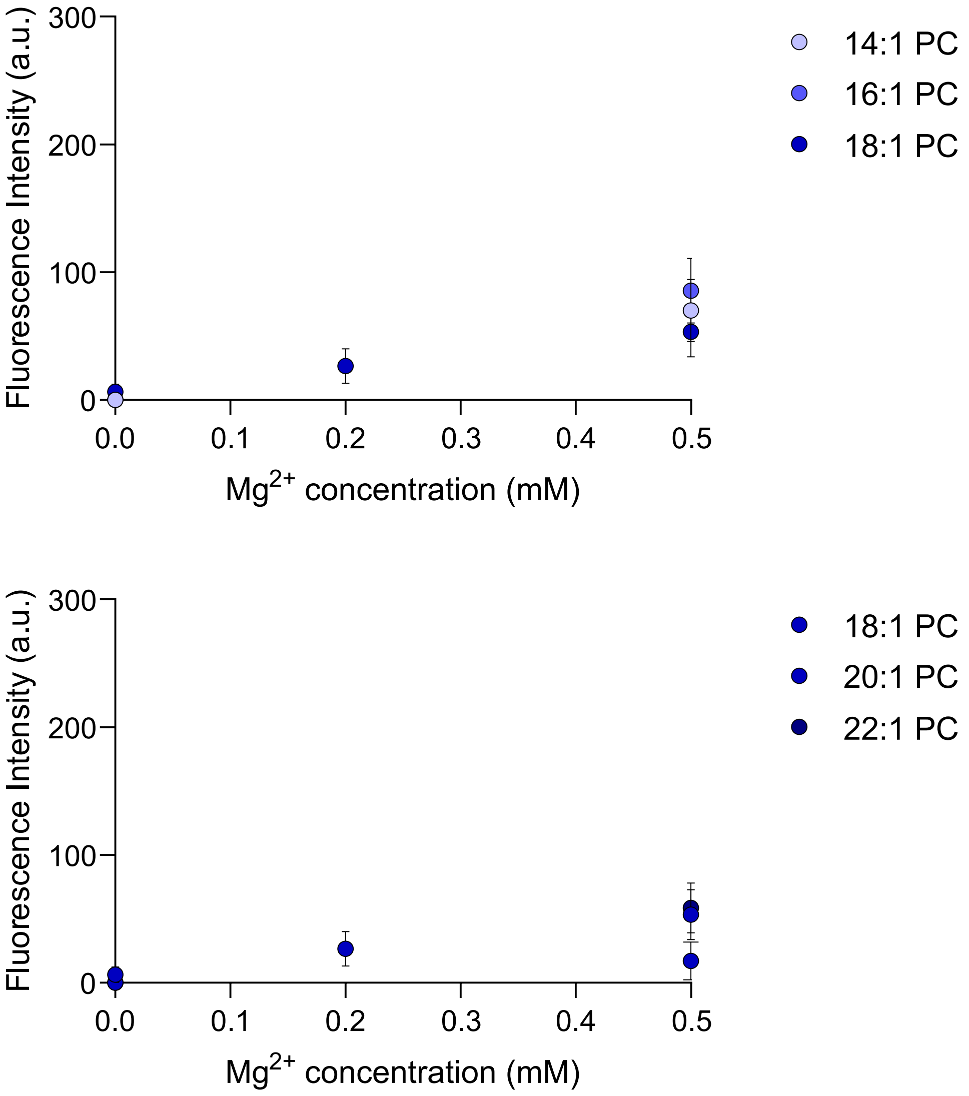


*Figure S24 - Fluorescence intensity of Cy3-labelled DNA-cholesterol conjugates as a function of $MgCl_2$ concentration for lipid membranes composed of (top) 14:1 (Δ9) PC, 16:1 (Δ9) PC and 18:1 (Δ9) PC, and (bottom) 18:1 (Δ9) PC, 20:1 (Δ11) PC and 22:1 (Δ13) PC. Membrane-associated fluorescence signal was quantified from confocal microscopy images. Data represent mean values with error bars indicating standard deviations from $n \geq 3$ independent measurements.*

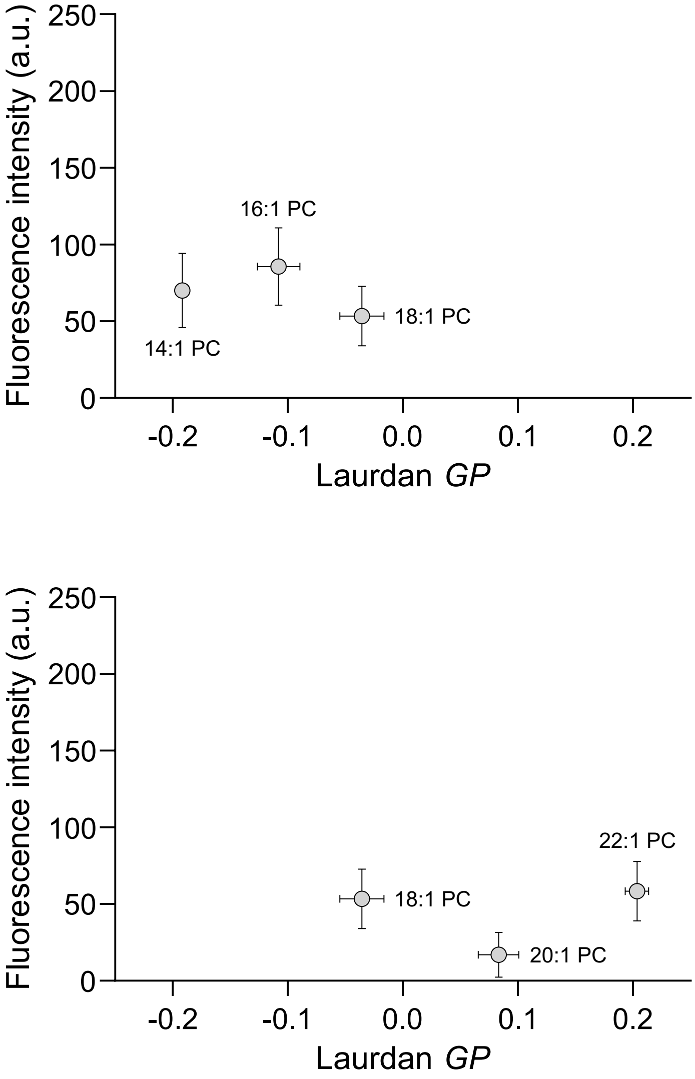


*Figure S25 - Fluorescence intensity of Cy3-labelled DNA-cholesterol conjugates at 0.5 mM $Mg^{2+}$ concentration as a function of Laurdan GP, which correlates with lipid packing, for lipid membranes composed of (top) 14:1 (Δ9) PC, 16:1 (Δ9) PC and 18:1 (Δ9) PC, and (bottom) 18:1 (Δ9) PC, 20:1 (Δ11) PC and 22:1 (Δ13) PC. Membrane-associated fluorescence signal was quantified from confocal microscopy images. Data represent mean values with error bars indicating standard deviations from n ≥ 3 independent measurements.*

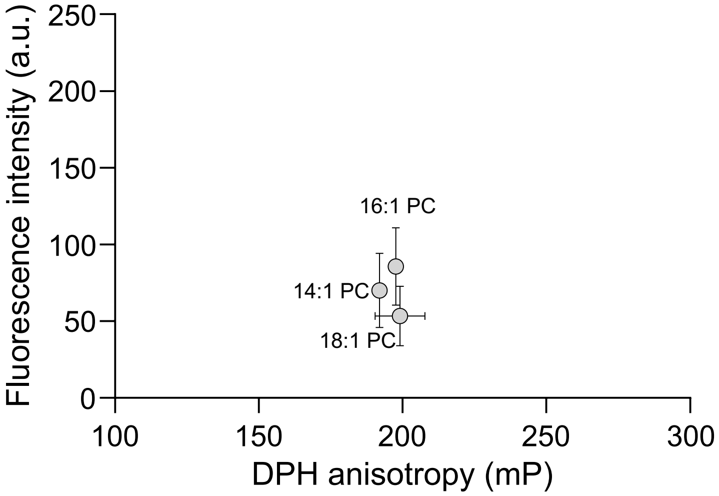


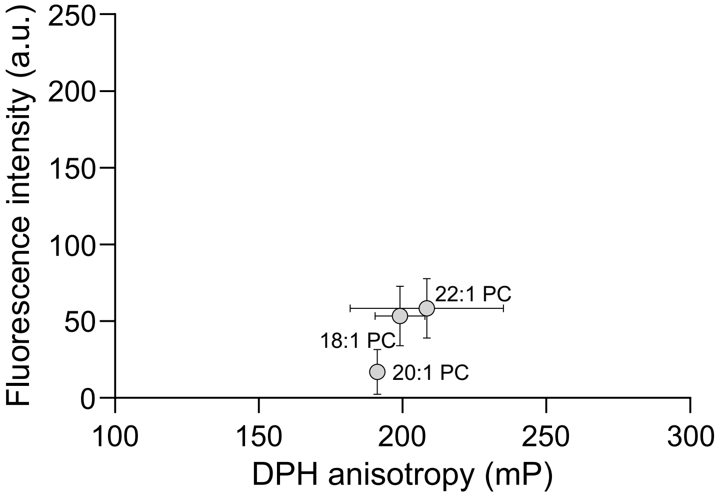


*Figure S26 - Fluorescence intensity of Cy3-labelled DNA-cholesterol conjugates at 0.5 mM $Mg^{2+}$ concentration as a function of DPH anisotropy, which correlates with membrane fluidity, for lipid membranes composed of (top) 14:1 (Δ9) PC, 16:1 (Δ9) PC and 18:1 (Δ9) PC, and (bottom) 18:1 (Δ9) PC, 20:1 (Δ11) PC and 22:1 (Δ13) PC. Membrane-associated fluorescence signal was quantified from confocal microscopy images. Data represent mean values with error bars indicating standard deviations from n ≥ 3 independent measurements.*

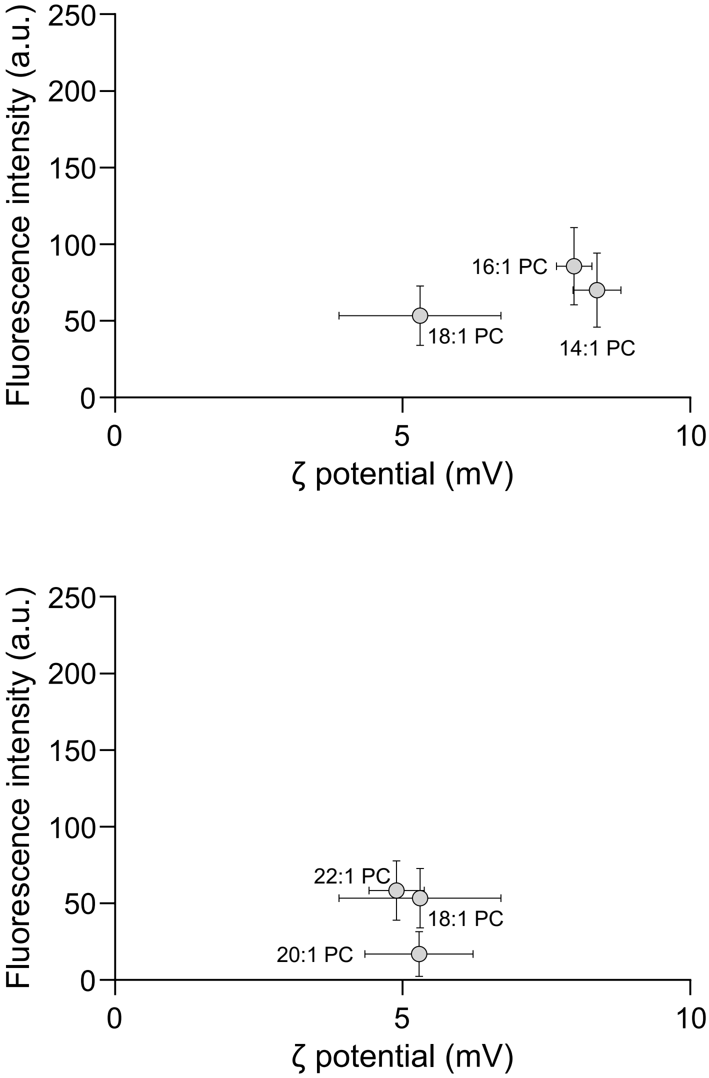


*Figure S27 - Fluorescence intensity of Cy3-labelled DNA-cholesterol conjugates at 0.5 mM $Mg^{2+}$ concentration as a function of Zeta potential, which correlates with membrane surface charge, for lipid membranes composed of (top) 14:1 (Δ9) PC, 16:1 (Δ9) PC and 18:1 (Δ9) PC, and (bottom) 18:1 (Δ9) PC, 20:1 (Δ11) PC and 22:1 (Δ13) PC. Membrane-associated fluorescence signal was quantified from confocal microscopy images. Data represent mean values with error bars indicating standard deviations from n ≥ 3 independent measurements.*

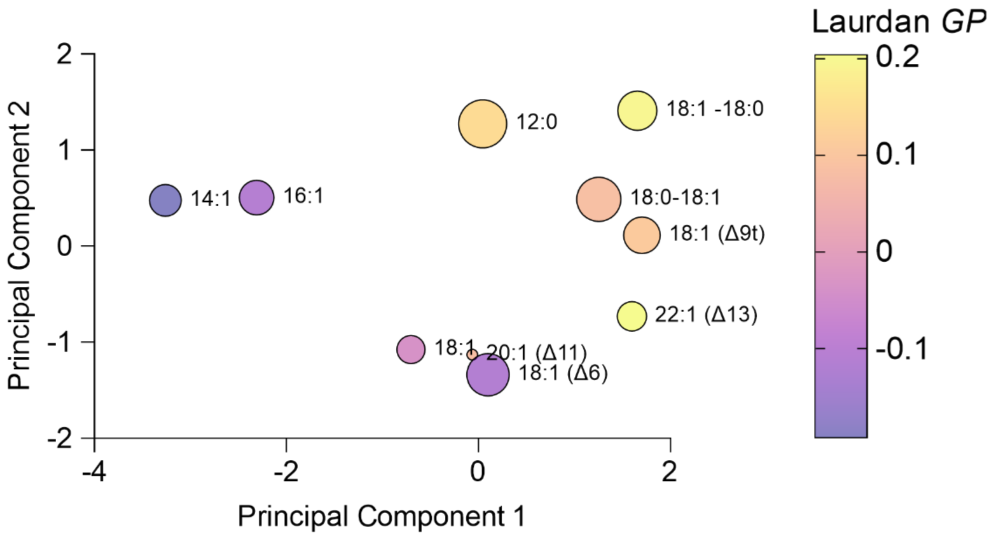


*Figure S28 - Principal component analysis (PCA) of membrane biophysical descriptors across the liquid-phase subgroup of the tested lipid library (Table S1). PCA was performed using lipid packing, membrane fluidity, surface charge, and melting temperature as input variables (PC1: 65.7%, PC2: 22.8%). Symbols are coloured according to lipid packing (inferred from Laurdan GP); the size of the symbols is proportional to the binding efficiency.*

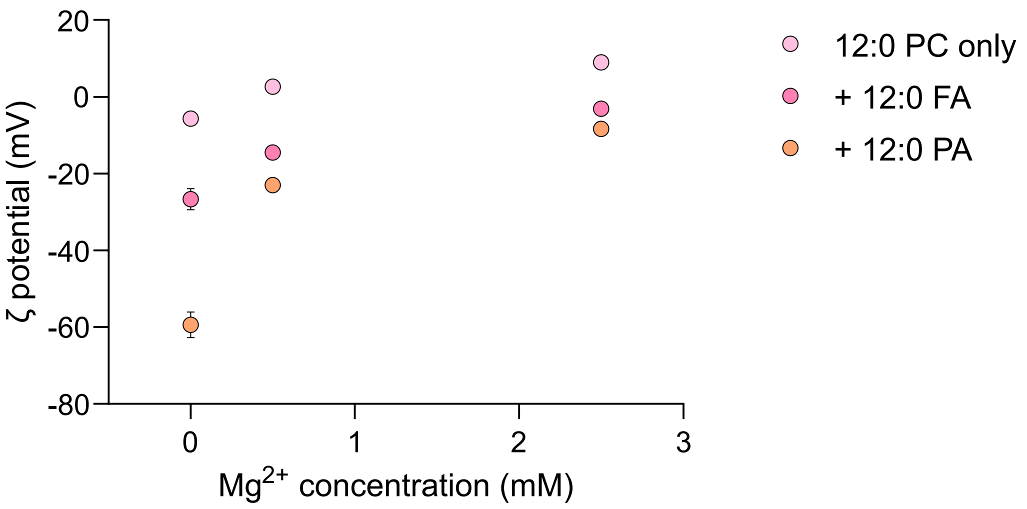


*Figure S29 - Zeta potential as a function of $MgCl_2$ concentration for lipid membranes composed of 12:0 PC, 80% 12:0 PC + 20% 12:0 FA or 80% 12:0 PC + 20% 12:0 PA. Data represent mean values with error bars indicating standard deviations from n ≥ 3 independent measurements.*

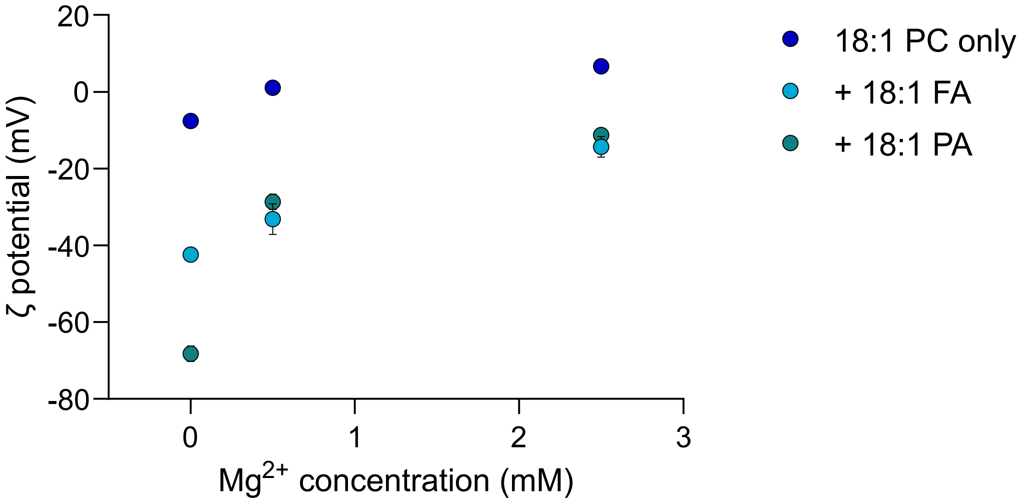


*Figure S30 - Zeta potential as a function of $MgCl_2$ concentration for lipid membranes composed of 18:1 PC, 80% 18:1 PC + 20% 18:1 FA or 80% 18:1 PC + 20% 18:1 PA. Data represent mean values with error bars indicating standard deviations from n ≥ 3 independent measurements.*

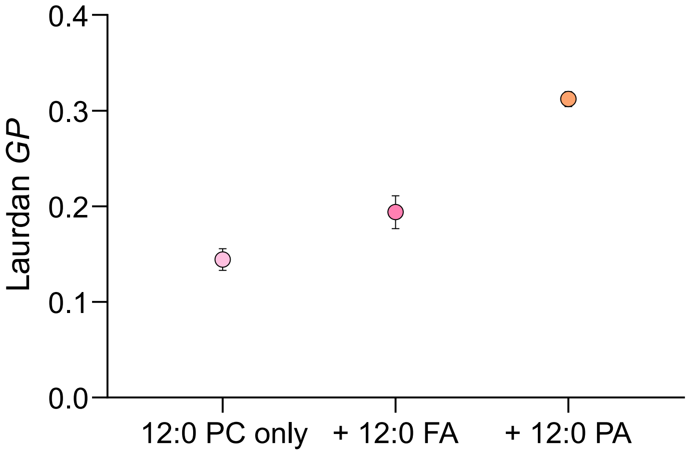


*Figure S31 - Laurdan GP for lipid membranes composed of 12:0 PC, 80% 12:0 PC + 20% 12:0 FA or 80% 12:0 PC + 20% 12:0 PA. Data represent mean values with error bars indicating standard deviations from n ≥ 3 independent measurements.*

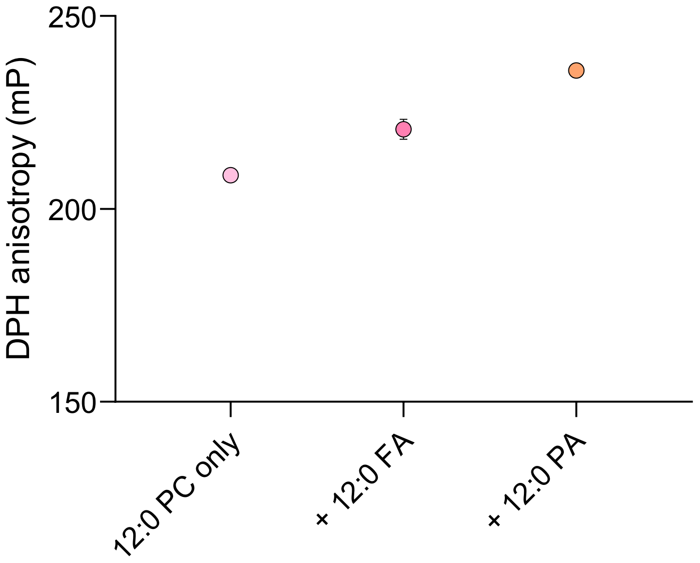


*Figure S32 - DPH anisotropy for lipid membranes composed of 12:0 PC, 80% 12:0 PC + 20% 12:0 FA or 80% 12:0 PC + 20% 12:0 PA. Data represent mean values with error bars indicating standard deviations from n ≥ 3 independent measurements.*

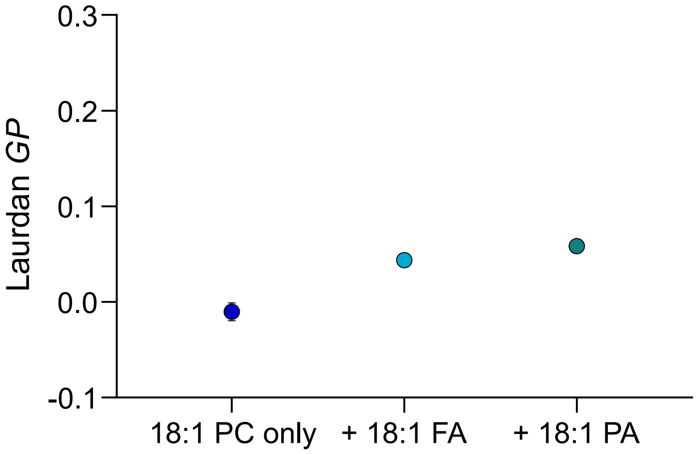


*Figure S33 - Laurdan GP for lipid membranes composed of 18:1 PC, 80% 18:1 PC + 20% 18:1 FA or 80% 18:1 PC + 20% 18:1 PA. Data represent mean values with error bars indicating standard deviations from n ≥ 3 independent measurements.*

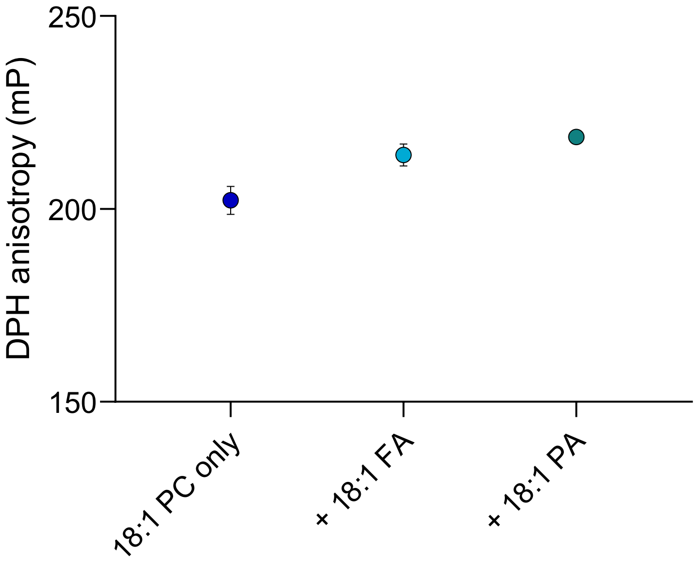


*Figure S34 - DPH anisotropy for lipid membranes composed of 18:1 PC, 80% 18:1 PC + 20% 18:1 FA or 80% 18:1 PC + 20% 18:1 PA. Data represent mean values with error bars indicating standard deviations from n ≥ 3 independent measurements.*

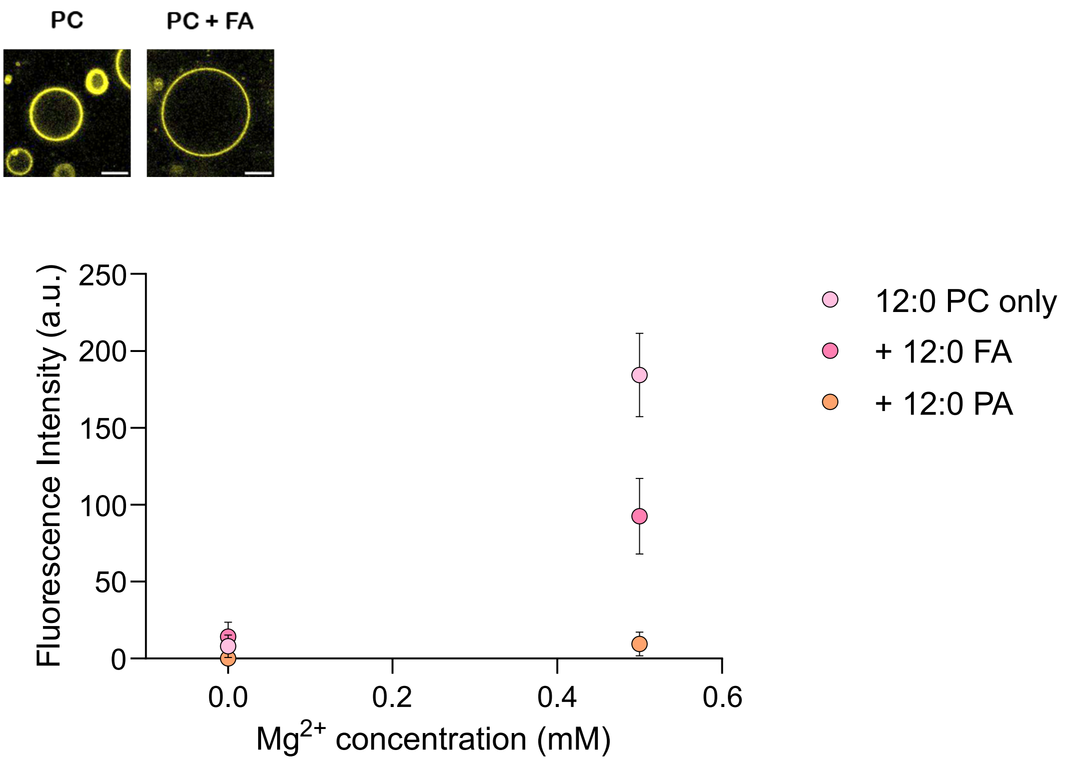


*Figure S35 - Fluorescence intensity of Cy3-labelled DNA-cholesterol conjugates (yellow) as a function of $MgCl_2$ concentration for lipid membranes composed of 12:0 PC, 80% 12:0 PC + 20% 12:0 FA or 80% 12:0 PC + 20% 12:0 PA. Membrane-associated fluorescence signal was quantified from confocal microscopy images (representative confocal micrographs are shown for 12:0 PC and 80% 12:0 PC + 20% 12:0 FA at 0.5 mM $Mg^{2+}$, scale bars = 20 μm). Trendlines are included as visual guides and do not represent any fitting. Data represent mean values with error bars indicating standard deviations from n ≥ 3 independent measurements.*

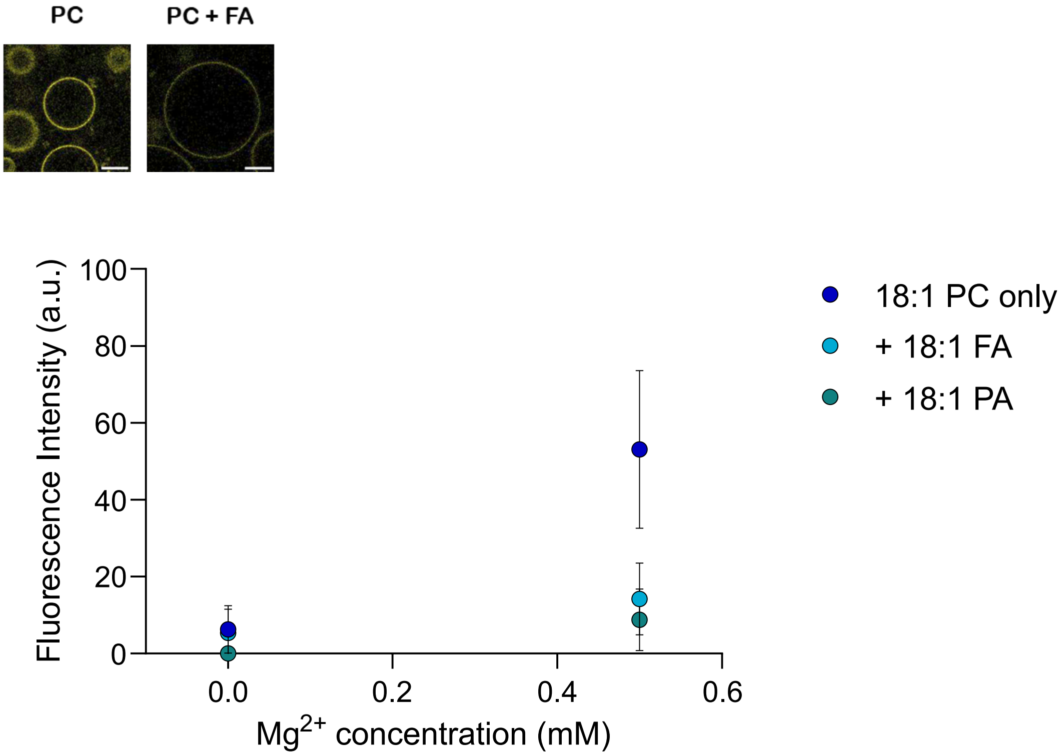


*Figure S36 - Fluorescence intensity of Cy3-labelled DNA-cholesterol conjugates (yellow) as a function of $MgCl_2$ concentration for lipid membranes composed of 18:1 PC, 80% 18:1 PC + 20% 18:1 FA or 80% 18:1 PC + 20% 18:1 PA. Membrane-associated fluorescence signal was quantified from confocal microscopy images (representative confocal micrographs are shown for 18:1 PC and 80% 18:1 PC + 20% 18:1 FA at 0.5 mM $Mg^{2+}$, scale bars = 20 µm). Trendlines are included as visual guides and do not represent any fitting. Data represent mean values with error bars indicating standard deviations from n ≥ 3 independent measurements.*

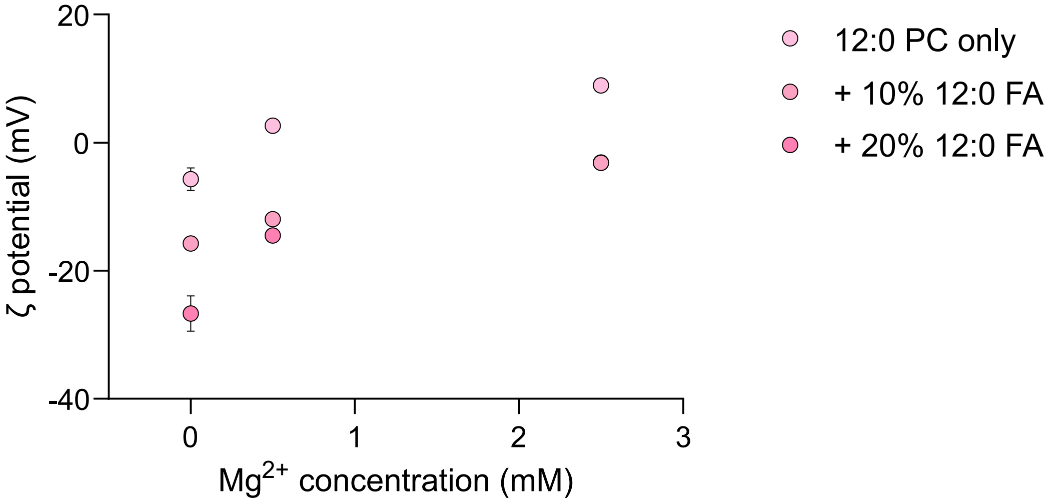


*Figure S37 - Zeta potential as a function of $MgCl_2$ concentration for lipid membranes composed of 12:0 PC, 90% 12:0 PC + 10% 12:0 FA or 80% 12:0 PC + 20% 12:0 FA. Data represent mean values with error bars indicating standard deviations from n ≥ 3 independent measurements.*

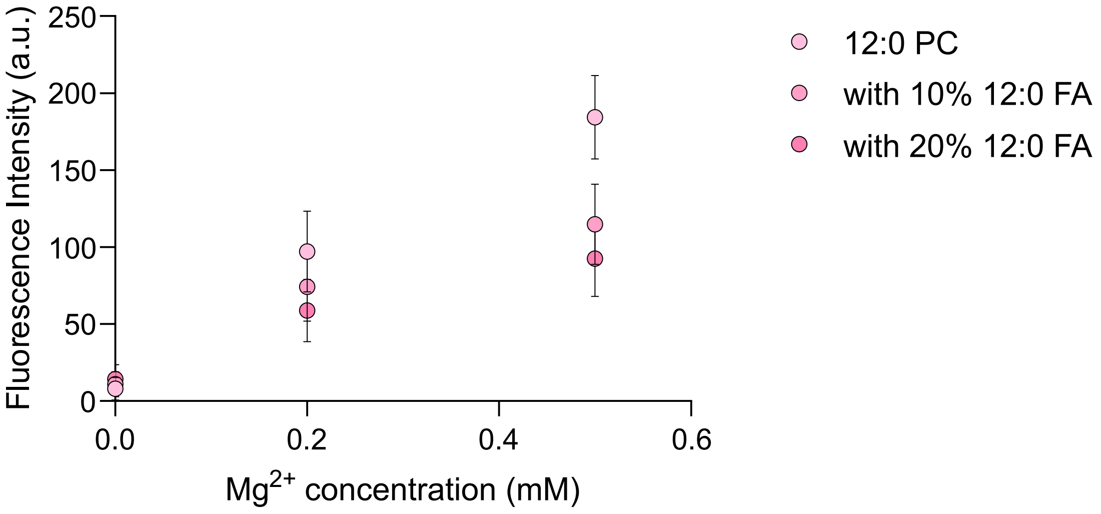


*Figure S38 - Fluorescence intensity of Cy3-labelled DNA-cholesterol conjugates as a function of $MgCl_2$ concentration for lipid membranes composed of 12:0 PC with increasing amounts of 12:0 FA. Membrane-associated fluorescence signal was quantified from confocal microscopy images. Data represent mean values with error bars indicating standard deviations from $n \geq 3$ independent measurements.*

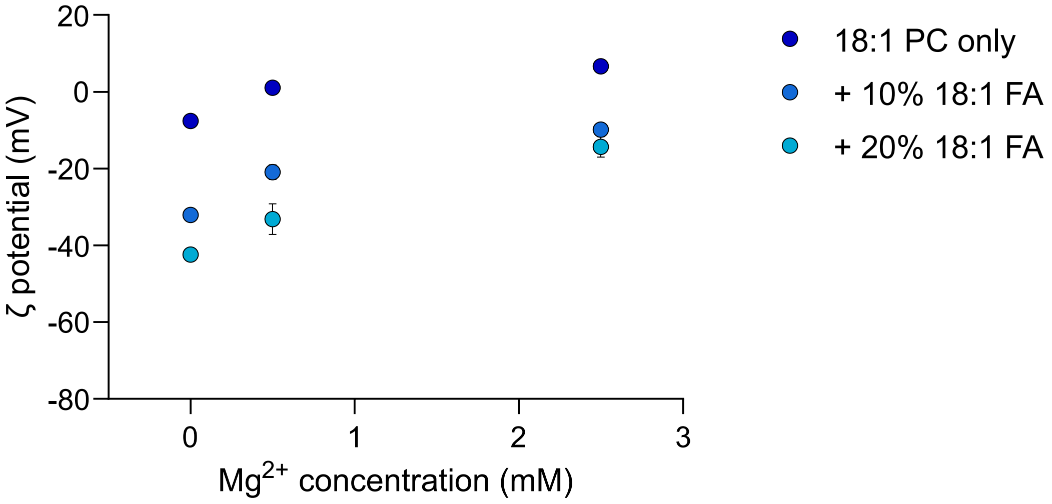


*Figure S39 - Zeta potential as a function of $MgCl_2$ concentration for lipid membranes composed of 12:0 PC, 90% 12:0 PC + 10% 12:0 FA or 80% 12:0 PC + 20% 12:0 FA. Data represent mean values with error bars indicating standard deviations from n ≥ 3 independent measurements.*

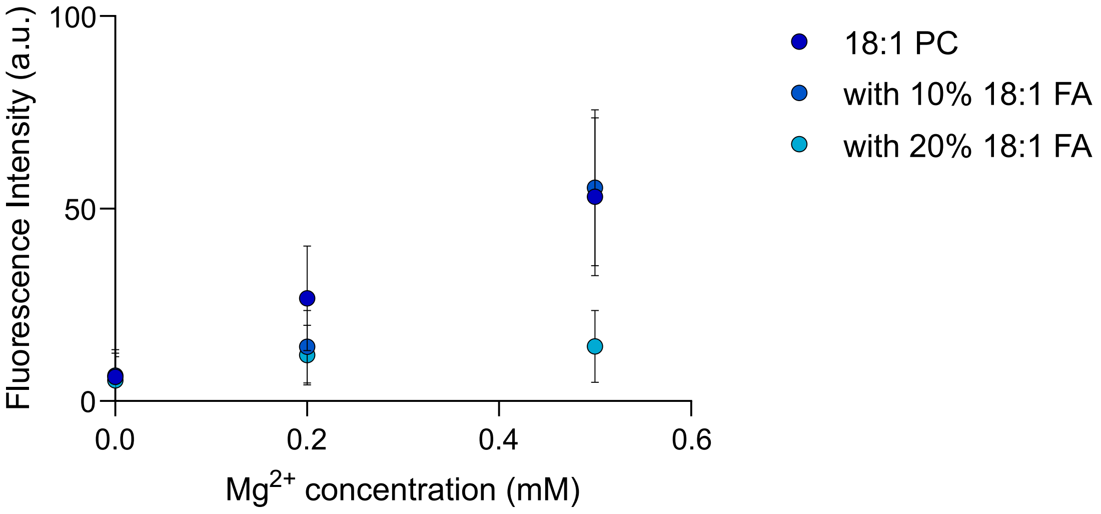


*Figure S40 - Fluorescence intensity of Cy3-labelled DNA-cholesterol conjugates as a function of $MgCl_2$ concentration for lipid membranes composed of 18:1 PC with increasing amounts of 18:1 FA. Membrane-associated fluorescence signal was quantified from confocal microscopy images. Data represent mean values with error bars indicating standard deviations from $n \geq 3$ independent measurements.*

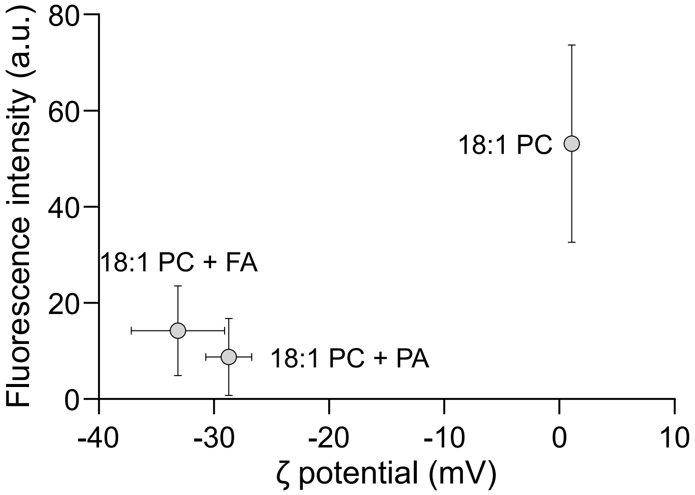


*Figure S41 - Fluorescence intensity of Cy3-labelled DNA-cholesterol conjugates at 0.5 mM $Mg^{2+}$ concentration as a function of Zeta potential, which correlates with membrane surface charge, for lipid membranes composed of 18:1 PC, 80% 18:1 PC + 20% 18:1 FA or 80% 18:1 PC + 20% 18:1 PA (Pearson correlation coefficient or r = 0.95). Membrane-associated fluorescence signal was quantified from confocal microscopy images. Data represent mean values with error bars indicating standard deviations from n ≥ 3 independent measurements.*

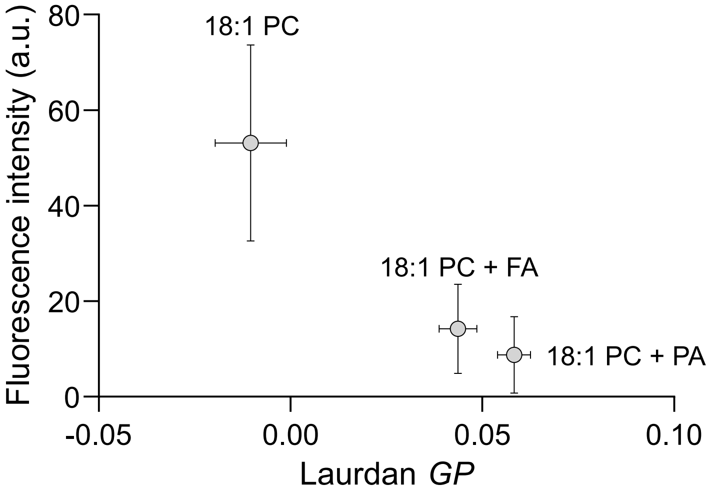


*Figure S42 - Fluorescence intensity of Cy3-labelled DNA-cholesterol conjugates at 0.5 mM $Mg^{2+}$ concentration as a function of Laurdan GP, which correlates with lipid packing, for lipid membranes composed of 18:1 PC, 80% 18:1 PC + 20% 18:1 FA or 80% 18:1 PC + 20% 18:1 PA (r = 0.99). Membrane-associated fluorescence signal was quantified from confocal microscopy images. Data represent mean values with error bars indicating standard deviations from n ≥ 3 independent measurements.*

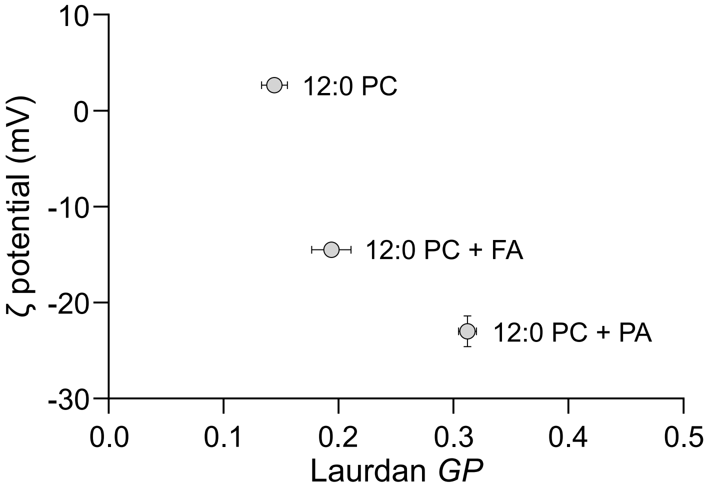


*Figure S43 - Zeta potential, which correlates with membrane surface charge, at 0.5 mM $Mg^{2+}$ concentration as a function of Laurdan GP for lipid membranes composed of 12:0 PC, 80% 12:0 PC + 20% 12:0 FA or 80% 12:0 PC + 20% 12:0 PA. Data represent mean values with error bars indicating standard deviations from n ≥ 3 independent measurements.*

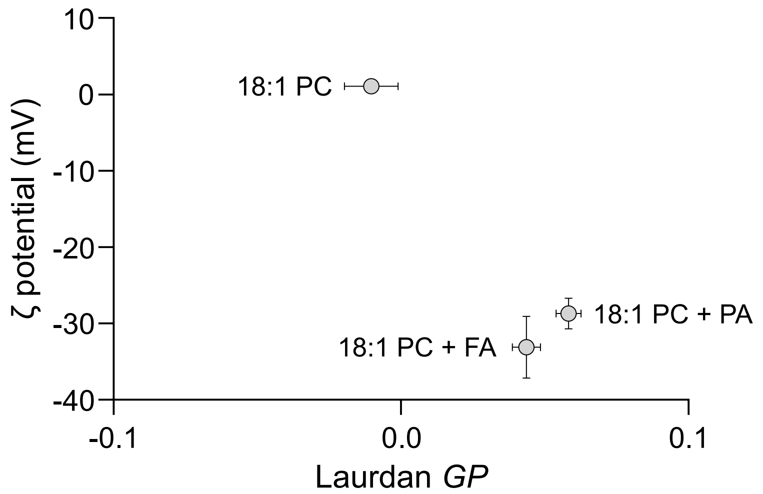


*Figure S44 - Zeta potential, which correlates with membrane surface charge, at 0.5 mM $Mg^{2+}$ concentration as a function of Laurdan GP for lipid membranes composed of 18:1 PC, 80% 18:1 PC + 20% 18:1 FA or 80% 18:1 PC + 20% 18:1 PA (r = 0.90). Data represent mean values with error bars indicating standard deviations from n ≥ 3 independent measurements.*

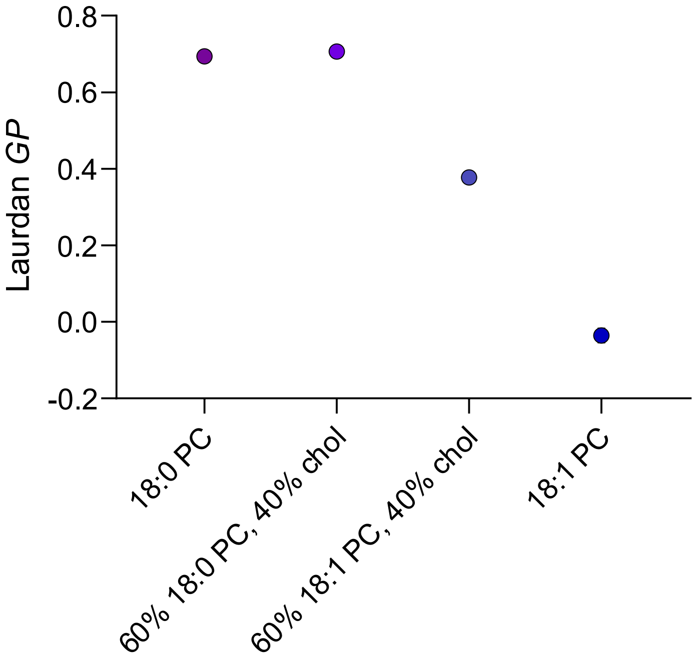


*Figure S45 - Laurdan GP for lipid membranes composed of 18:0 PC, 60% 18:0 PC + 40% cholesterol, 60% 18:1 PC + 40% cholesterol and 18:1 PC in the absence of $Mg^{2+}$. Data represent mean values with error bars indicating standard deviations from n ≥ 3 independent measurements.*

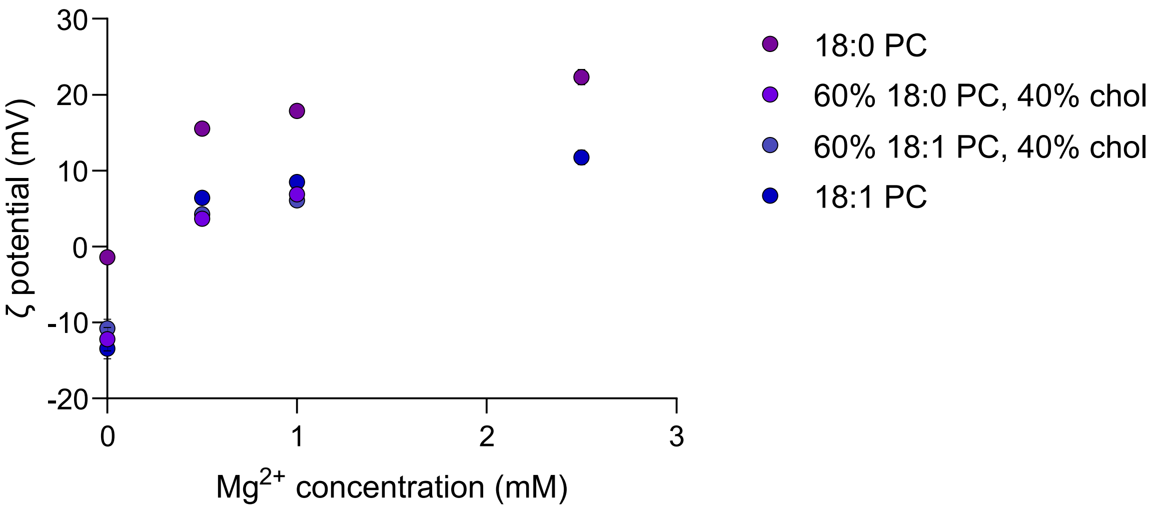


*Figure S46 - Zeta potential as a function of $MgCl_2$ concentration for lipid membranes composed of 18:0 PC, 60% 18:0 PC + 40% cholesterol, 60% 18:1 PC + 40% cholesterol and 18:1 PC. Data represent mean values with error bars indicating standard deviations from n ≥ 3 independent measurements.*

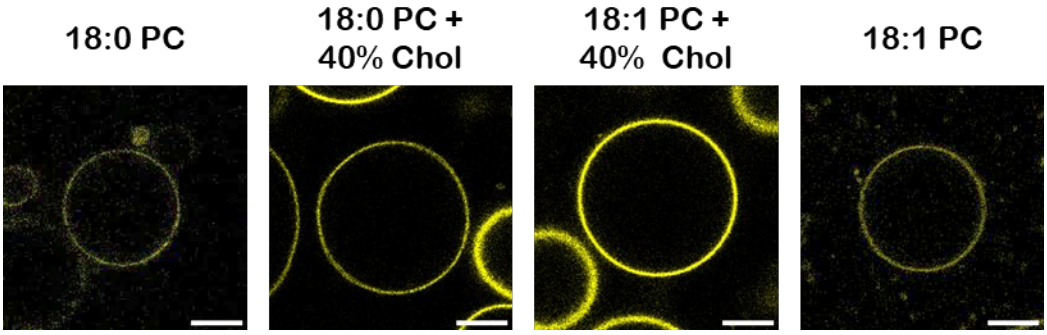


*Figure S47 - Confocal micrographs showing membrane binding of amphiphilic Cy3-DNA nanoprobes (yellow) for different membrane phases at increasing concentrations of $MgCl_2$. Scale bar: 10 µm. n ≥ 3 independent measurements.*

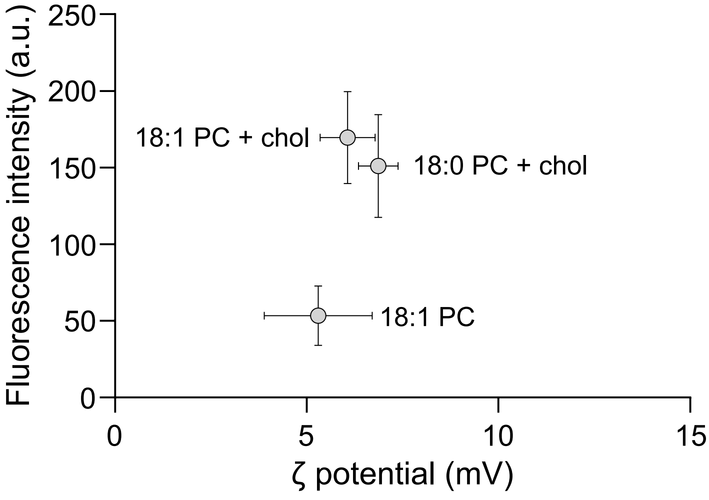


*Figure S48 - Fluorescence intensity of Cy3-labelled DNA-cholesterol conjugates at 0.5 mM $Mg^{2+}$ concentration as a function of Zeta potential, which correlates with membrane surface charge, for liquid-phase membranes composed of 60% 18:0 PC + 40% cholesterol, 60% 18:1 PC + 40% cholesterol and 18:1 PC. Membrane-associated fluorescence signal was quantified from confocal microscopy images. Data represent mean values with error bars indicating standard deviations from n ≥ 3 independent measurements.*

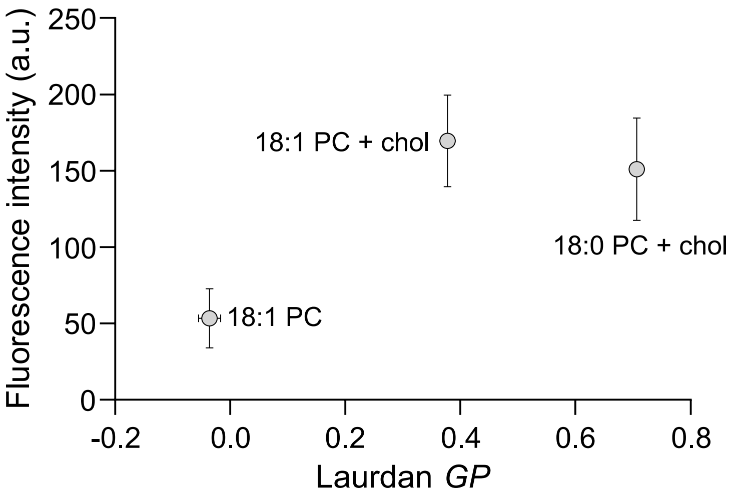


*Figure S49 - Fluorescence intensity of Cy3-labelled DNA-cholesterol conjugates at 0.5 mM $Mg^{2+}$ concentration as a function of Laurdan GP, which correlates with lipid packing, for liquid-phase membranes composed of 60% 18:0 PC + 40% cholesterol, 60% 18:1 PC + 40% cholesterol and 18:1 PC. Membrane-associated fluorescence signal was quantified from confocal microscopy images. Data represent mean values with error bars indicating standard deviations from n ≥ 3 independent measurements.*

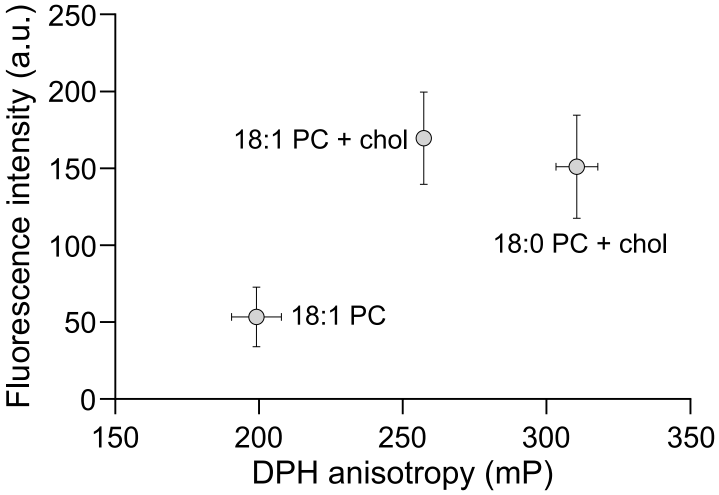


*Figure S50 - Fluorescence intensity of Cy3-labelled DNA-cholesterol conjugates at 0.5 mM $Mg^{2+}$ concentration as a function of DPH anisotropy, which correlates with membrane fluidity, for liquid-phase membranes composed of 60% 18:0 PC + 40% cholesterol, 60% 18:1 PC + 40% cholesterol and 18:1 PC. Membrane-associated fluorescence signal was quantified from confocal microscopy images. Data represent mean values with error bars indicating standard deviations from n ≥ 3 independent measurements.*

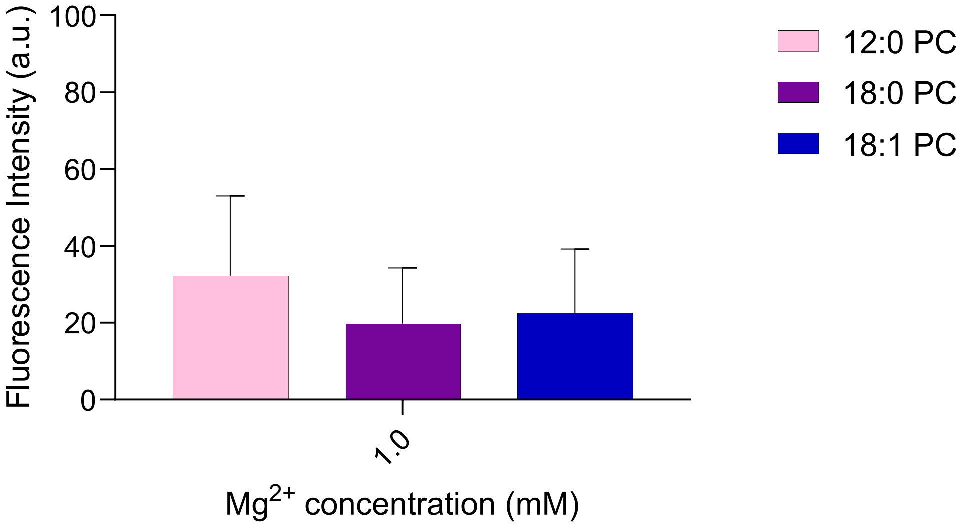


*Figure S51 - Fluorescence intensity of FAM-labelled DNA-cholesterol conjugates at 1 mM $Mg^{2+}$ concentration for lipid membranes composed of 12:0 PC, 18:0 PC and 18:1 PC. Membrane-associated fluorescence signal was quantified from confocal microscopy images. Lower fluorescence intensities are observed for FAM-labelled DNA compared to Cy3-labelled DNA throughout this study. Data represent mean values with error bars indicating standard deviations from n ≥ 3 independent measurements.*

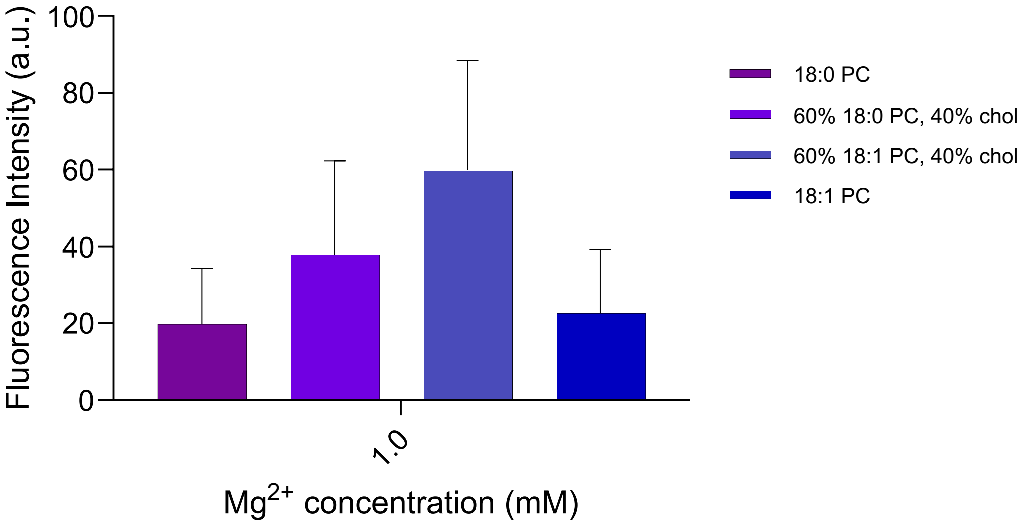


*Figure S52 - Fluorescence intensity of FAM-labelled DNA-cholesterol conjugates at 1 mM $Mg^{2+}$ concentration for lipid membranes composed of 18:0 PC, 60% 18:0 PC + 40% cholesterol, 60% 18:1 PC + 40% cholesterol and 18:1 PC. Membrane-associated fluorescence signal was quantified from confocal microscopy images. Data represent mean values with error bars indicating standard deviations from $n \geq 3$ independent measurements.*

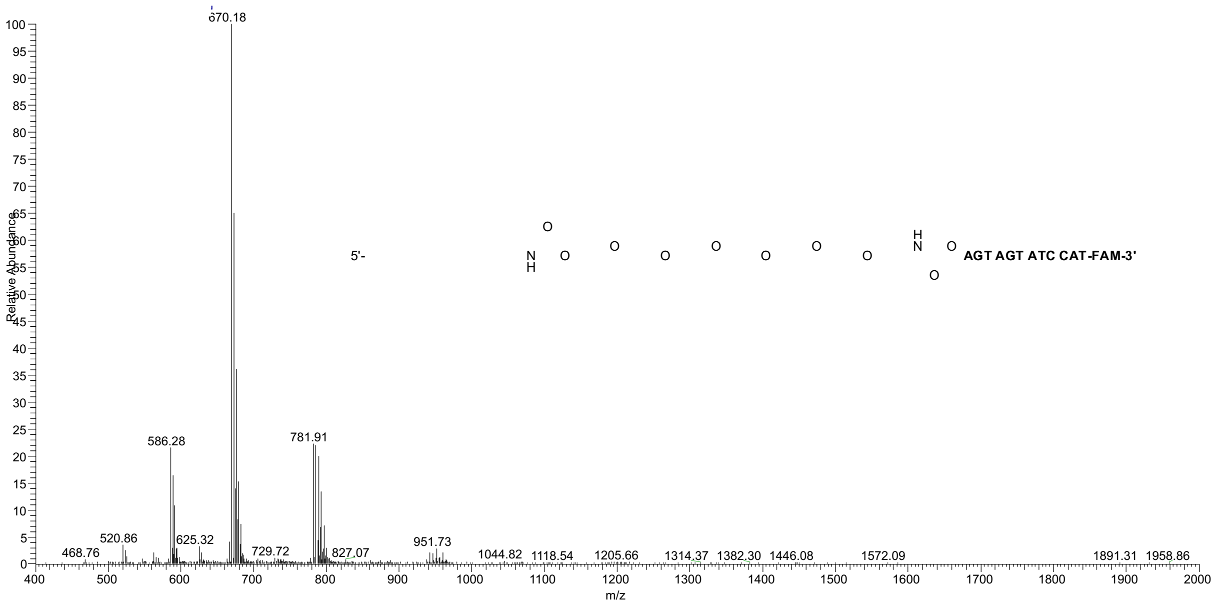


*Figure S53 - LR-ESI-MS of C10*-DNA-FAM. Expected masses: $[M-8H]^{-8}$ 586.3; $[M-7H]^{-7}$ 670.2; $[M-6H]^{-6}$ 782.0; found: 586.3, 670.2, 781.9.*

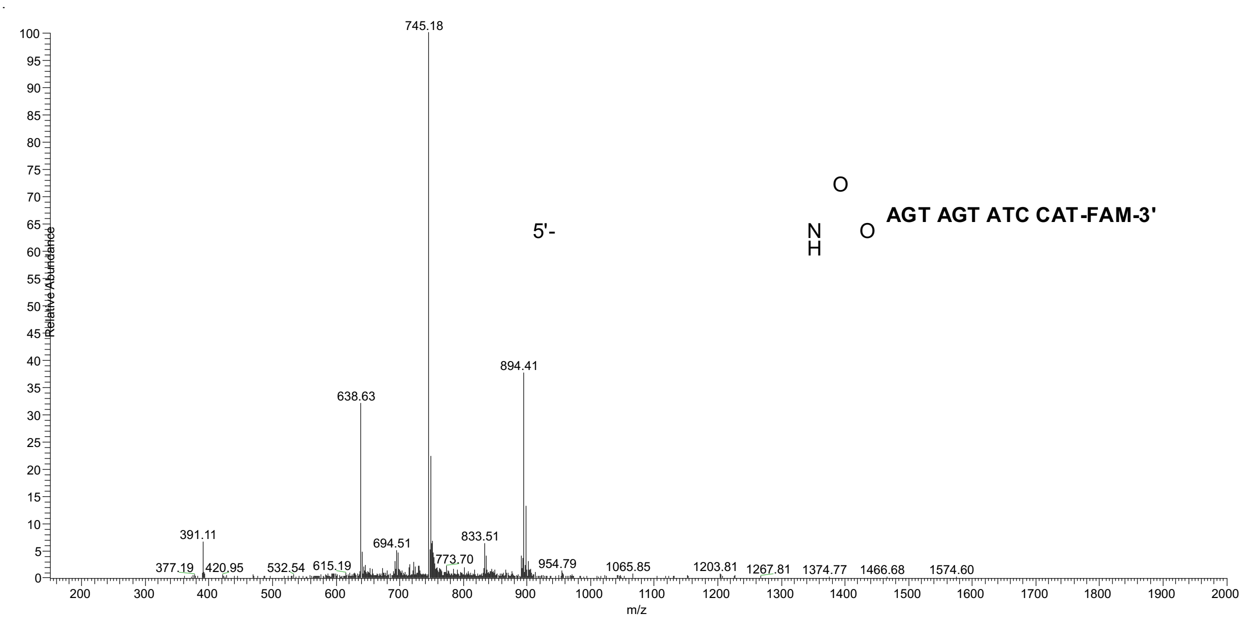


*Figure S54 - LR-ESI-MS of C10-DNA-FAM. Expected masses: $[M\text{-}7H]^{-7}$ 638.9; $[M\text{-}6H]^{-6}$ 745.5; $[M\text{-}5H]^{-5}$ 894.8; found: 638.6, 745.2, 894.4.*

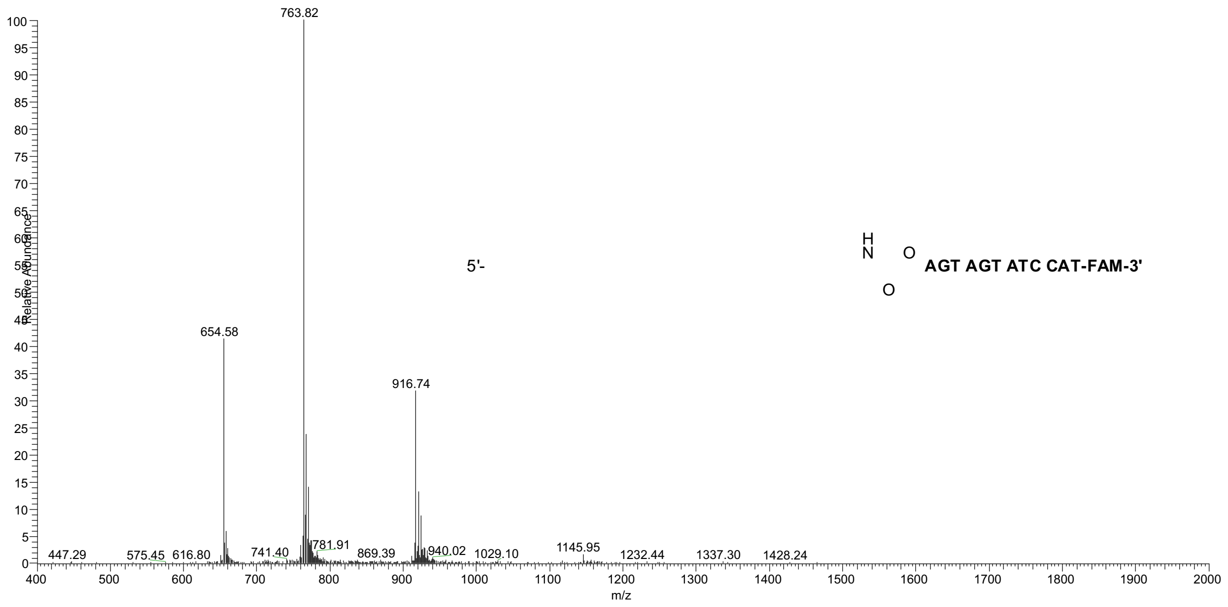


*Figure S55 - LR-ESI-MS of C18:1-DNA-FAM. Expected masses: $[M-7H]^{-7}$ 654.6; $[M-6H]^{-6}$ 763.8; $[M-5H]^{-5}$ 916.8; found: 654.6, 763.82, 916.7.*

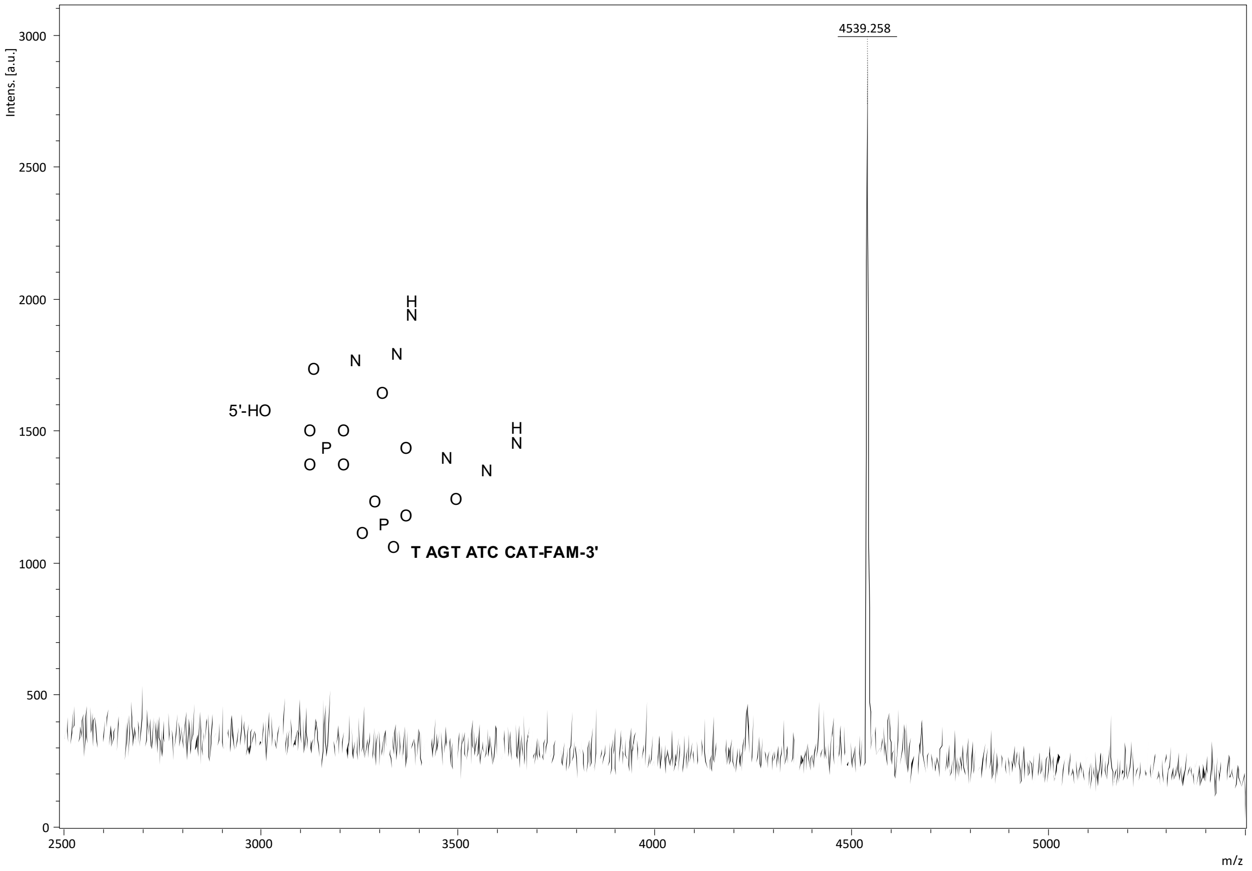


*Figure S56 - MALDI-TOF/TOF MS of 2xC10-DNA-FAM. Expected mass: [M-H]$^-$ 4539.162; found: 4539.258.*

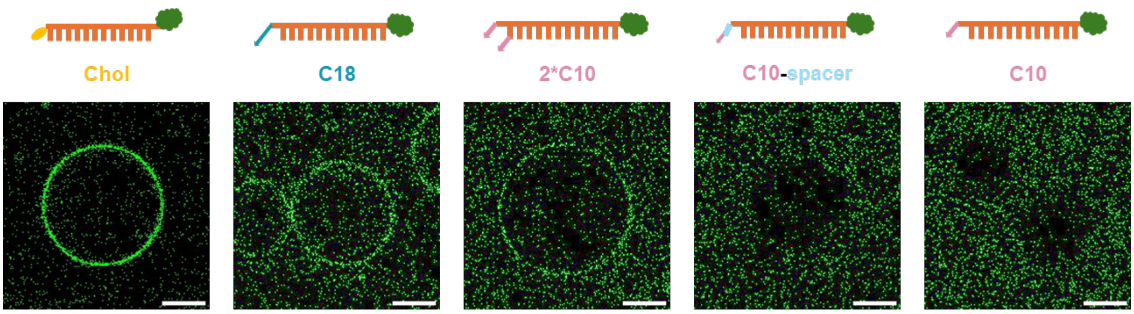


*Figure S57 - Confocal micrographs showing 12:0 PC-based membrane binding of different amphiphilic FAM-DNA nanoprobes (green) in the presence of 0.5 mM $Mg^{2+}$. Scale bar: 10 µm. n ≥ 3 independent measurements.*

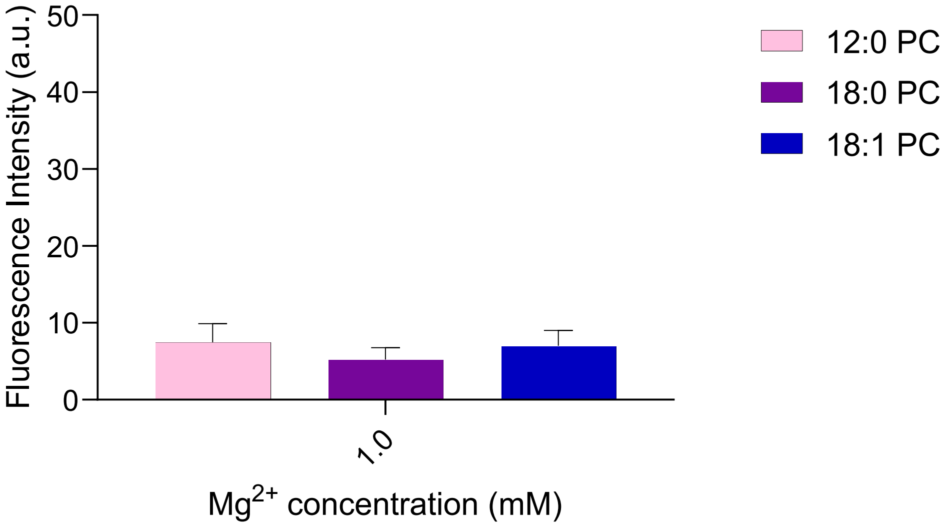


*Figure S58 - Fluorescence intensity of FAM-labelled DNA-oleic acid conjugates at 0 mM and 1 mM $Mg^{2+}$ concentration for lipid membranes composed of 12:0 PC, 18:0 PC and 18:1 PC. Membrane-associated fluorescence signal was quantified from confocal microscopy images. Data represent mean values with error bars indicating standard deviations from $n \geq 3$ independent measurements.*

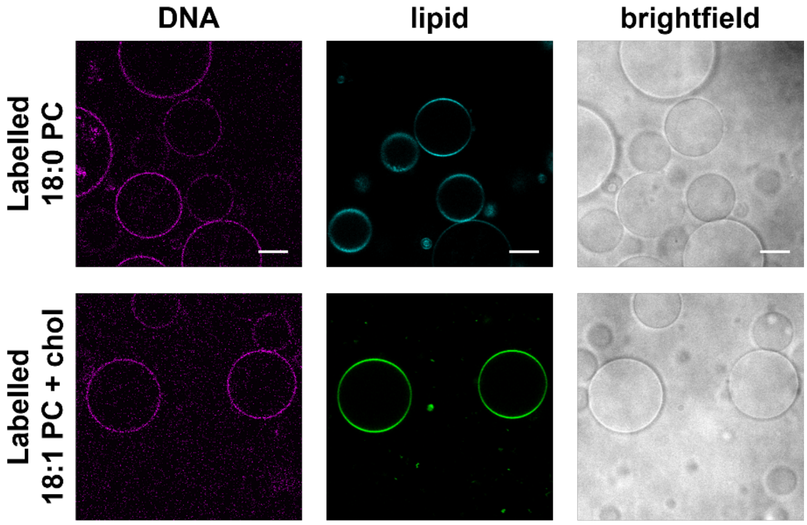


*Figure S59 - Confocal micrographs showing amphiphilic Cy3-DNA nanoprobes (magenta) preferentially interacting with 18:1 PC + chol GUVs in a solution containing both 18:1 PC + chol and 18:0 PC GUVs in the presence of 0.5 mM $Mg^{2+}$. Experiments were performed with either 18:0 PC GUVs labelled with 0.8% Texas Red-PE (cyan) and unlabelled 18:1 PC + chol GUVs or 18:1 PC + chol GUVs labelled with 0.8% Texas Red-PE (green) and unlabelled 18:0 PC GUVs. Scale bar: 10 µm. n ≥ 3 independent measurements.*

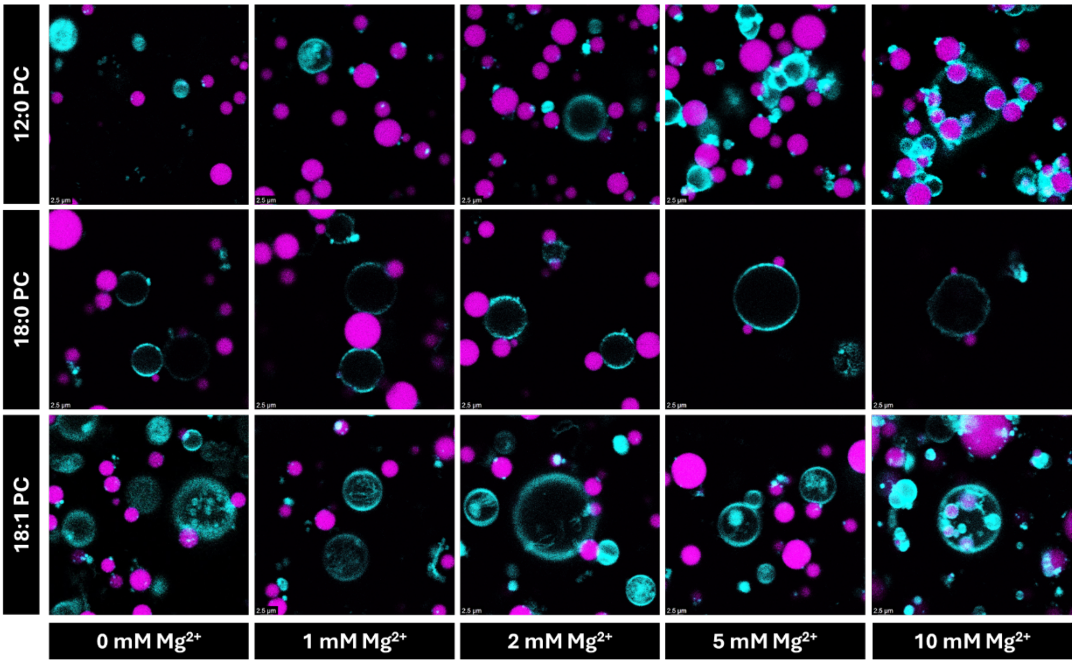


*Figure S60 - Confocal micrographs showing DNA-based coacervates-membrane interactions in the presence of DNA at increasing concentrations of $MgCl_2$. Coacervates (magenta) are labelled with 1% FAM-labelled DNA, lipid membranes (cyan) are labelled with 0.8% Texas Red-PE. Scale bar: 2.5 μm. $n \geq 3$ independent measurements.*

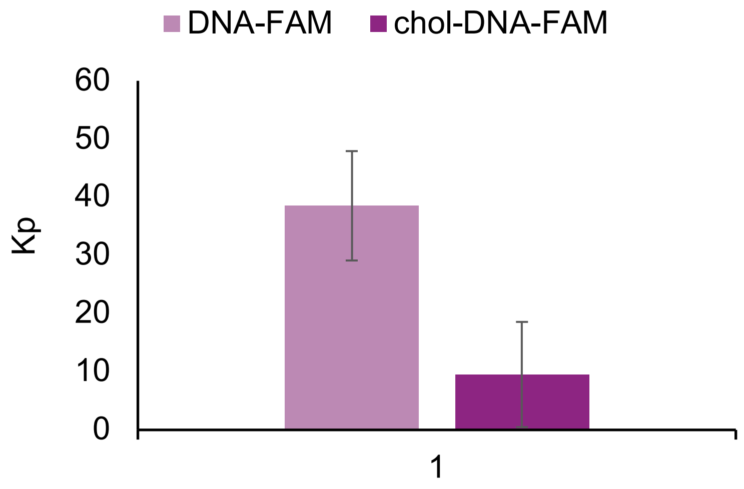


*Figure S61 - Partitioning of FAM-labelled DNA with or without cholesterol tag (1% fluorescent oligonucleotide) in $R_4$/$DNA_{12}$ coacervates. Data represent mean values with error bars indicating standard deviations from $n \geq 3$ independent measurements.*

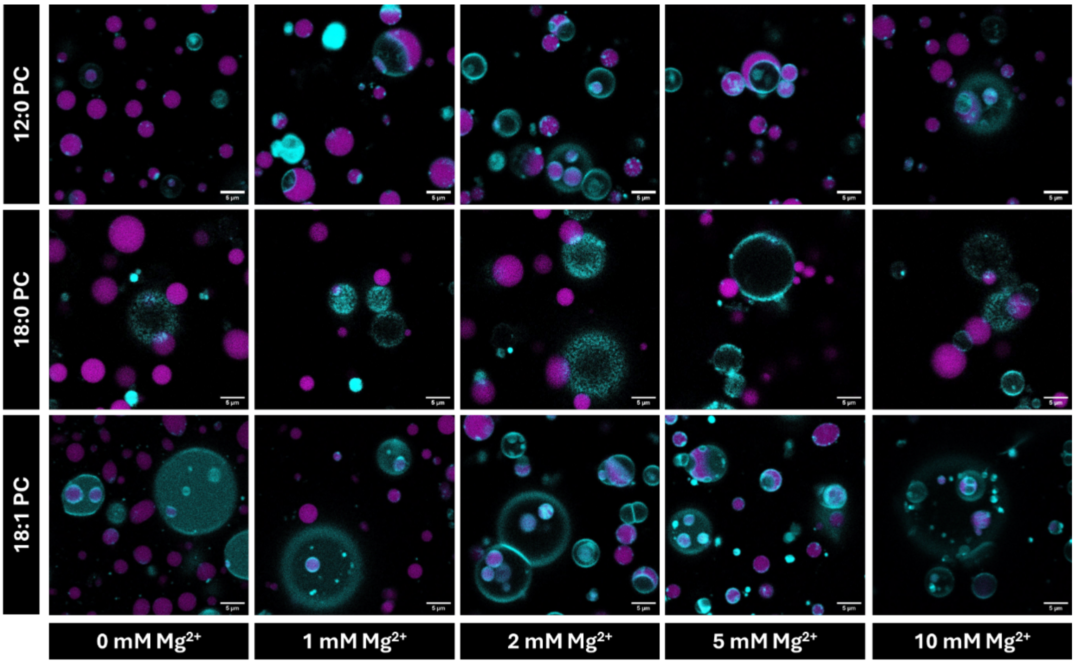


*Figure S62 - Confocal micrographs showing DNA-based coacervates-membrane interactions in the presence of cholesterol-tagged DNA at increasing concentrations of $MgCl_2$. Coacervates (magenta) are labelled with 1% FAM-labelled DNA, lipid membranes (cyan) are labelled with 0.8% Texas Red-DHPE. Scale bar: 5 μm. n ≥ 3 independent measurements.*

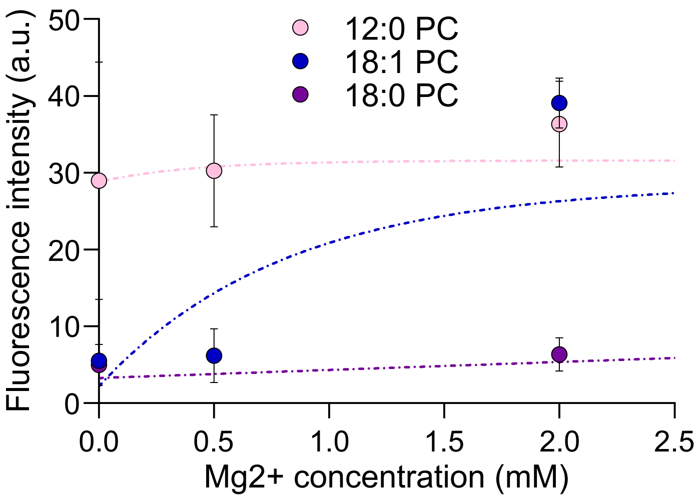


*Figure S63 - DNA-based coacervates-membrane interactions in the presence of cholesterol-tagged DNA at increasing concentrations of $MgCl_2$ determined by quantification of the fluorescence intensity overlaps between coacervates and GUVs, extracted from confocal micrographs. Data represent mean values with error bars indicating standard deviations from n ≥ 3 independent measurements.*
